# Secure and Federated Genome-Wide Association Studies for Biobank-Scale Datasets

**DOI:** 10.1101/2022.11.30.518537

**Authors:** Hyunghoon Cho, David Froelicher, Jeffrey Chen, Manaswitha Edupalli, Apostolos Pyrgelis, Juan R. Troncoso-Pastoriza, Jean-Pierre Hubaux, Bonnie Berger

## Abstract

Sharing data across institutions for genome-wide association studies (GWAS) would enhance the discovery of genetic variants linked to health and disease^1, 2^. However, existing data sharing regulations limit the scope of such collaborations^3^. Although cryptographic tools for secure computation promise to enable collaborative analysis with formal privacy guarantees, existing approaches either are computationally impractical or support only simplified analyses^4–7^. We introduce secure federated genome-wide association studies (SF-GWAS), a novel combination of secure computation frameworks and distributed algorithms that empowers efficient and accurate GWAS on private data held by multiple entities while ensuring data confidentiality. SF-GWAS supports the most widely-used GWAS pipelines based on principal component analysis (PCA) or linear mixed models (LMMs). We demonstrate the accuracy and practical runtimes of SF-GWAS on five datasets, including a large UK Biobank cohort of 410K individuals, showcasing an order-of-magnitude improvement in runtime compared to previous work. Our work realizes the power of secure collaborative genomic studies at unprecedented scale.

## Main

Genome-wide association studies (GWAS) are an essential tool for identifying genetic determinants of complex biological traits. Recent collaborative studies have demonstrated the power of jointly analyzing multiple datasets in detecting a greater number of associations than possible using any single dataset^1, 2^. However, such collaborations are still rare and typically restricted to limited types of analyses that do not require the sharing of sensitive individual-level data (e.g., based on summary statistics). These limitations are largely due to institutional policies and regulations that prevent sharing of sensitive genetic data^3^.

Secure computation techniques from modern cryptography offer promising strategies to address safety concerns in collaborative data sharing^3, 8–10^. For example, these techniques allow a group of collaborators to jointly analyze their collective data by exchanging encrypted information, while guaranteeing each party’s data is private from others. Although recent work has illustrated the potential of secure computation for collaborative GWAS^4–7^, existing methods are prohibitively slow for large datasets or implement a simplified analysis pipeline, such as one that does not correct for confounding due to population structure, which is essential for accurate GWAS^11, 12^. These issues severely limit the utility of existing methods in practice.

We introduce SF-GWAS, a secure and federated algorithm for multi-site GWAS, which overcomes these challenges to enable provably secure genetics collaborations at unprecedented scale (Fig. 1). SF-GWAS builds upon the following two key conceptual advances. First is our “federated” framework for secure computation, whereby each genomic dataset input is kept at the respective source site to minimize both computational and network communication burden. We enable this strategy using a novel combination of two cryptographic frameworks—namely, secure multiparty computation (MPC) and homomorphic encryption (HE). HE refers to data encryption schemes that allow computation to be performed directly on encrypted data, whereas MPC refers to a class of interactive protocols that allow multiple parties to perform computation on private data that is split among them with ensured data confidentiality. Our framework uses both techniques to obtain practical performance for our methods: each party uses HE to perform efficient *local* computation by utilizing the plaintext (unencrypted) input data and sharing only encrypted intermediate results with other parties to cooperatively carry out global computations. At the same time, the parties employ efficient MPC routines for complex operations (e.g., division and comparison) to mitigate the computational cost of HE on encrypted data and to enhance the accuracy of operations over a wider range of data values. Importantly, prior work based on a multiparty extension of HE^5^, including multiparty HE (MHE), did not utilize any computational MPC operations, which as our results show are critical for robust accuracy of our methods across datasets of varying sizes.

**Figure 1.**
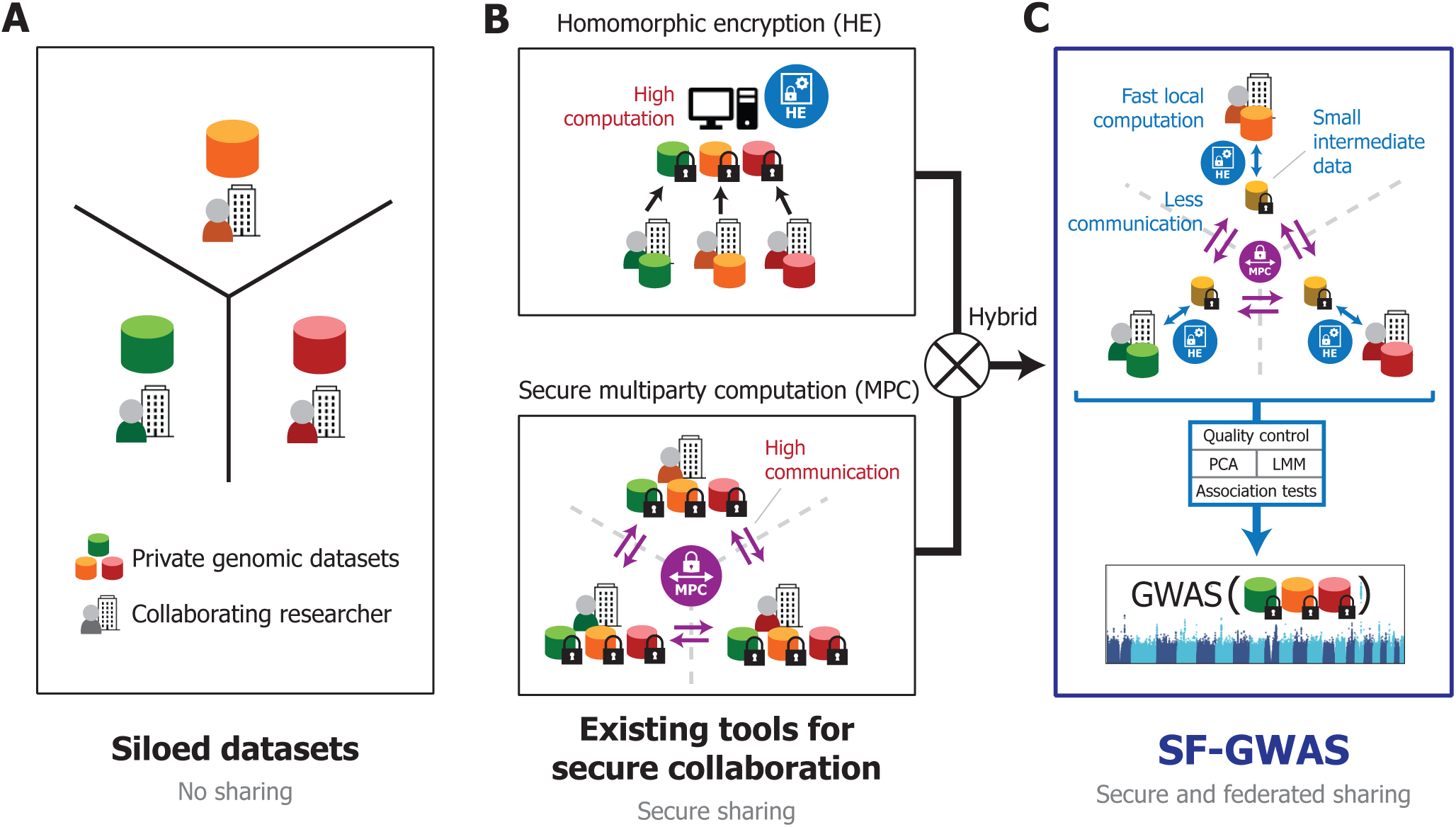
Overview of SF-GWAS. (**A**) SF-GWAS addresses a common challenge faced by collaborating researchers at different institutions who wish to conduct a joint study, but are unable to do so due to data sharing limitations based on privacy concerns. (**B**) Modern cryptographic solutions for jointly analyzing private datasets with formal privacy guarantees include homomorphic encryption (HE) and secure multiparty computation (MPC). However, existing solutions for GWAS have limited scalability due to the high costs of computation and communication incurred by these frameworks. (**C**) SF-GWAS is built upon a novel combination of HE and MPC to enable secure and federated computation, where large private datasets are locally kept by each data holder and only small intermediate data are encrypted and shared among the collaborators to carry out complex global computations. SF-GWAS introduces practical, secure, and federated algorithms to support two essential workflows for GWAS based on principal component analysis (PCA) and linear mixed models (LMMs). The final result includes GWAS association statistics, jointly computed over all private datasets while preserving data privacy. We further illustrate our novel algorithm for secure and federated LMM-based association analysis in Supplementary Figure 1.

Secondly, SF-GWAS introduces efficient algorithmic design to support the federated execution of a wide range of end-to-end GWAS pipelines. The SF-GWAS workflow includes, prior to association tests, quality control and two essential strategies to correct for population structure in the cohort^11, 12^: principal component analysis (PCA) or linear mixed models (LMMs). Both PCA and LMMs involve highly complex linear algebra computations, which thus far have limited the development of privacy-preserving algorithms that can scale to modern large-scale genomic datasets. Leveraging a range of novel algorithmic design strategies, we developed practical, secure and federated algorithms for both PCA-and LMM-based GWAS workflows, where the former supports both linear and logistic regression-based tests, illustrating the general applicability of our techniques. SF-GWAS achieves an order-of-magnitude improvement in runtime compared to the state-of-the-art approach for PCA-based GWAS^4^ while also providing significantly stronger privacy guarantees. SF-GWAS provides, to our knowledge, the first practical secure algorithms for both logistic regression-based and LMM-based GWAS (Supplementary Fig. 1). We provide the algorithmic details of SF-GWAS in Methods and Supplementary Notes 1–9.

To demonstrate the significant computational benefits of our secure and federated approach to collaborative GWAS, we compared SF-GWAS with the prior state-of-the-art method for secure PCA-based GWAS^4^, referred to as S-GWAS. We used both methods to analyze the three real datasets from the S-GWAS publication (see Methods): a lung cancer dataset (n=9,178 individuals), a bladder cancer dataset (n=13,060), and an age-related macular degeneration (AMD) dataset (n=22,683). Each dataset was randomly split into two subsets, which we distributed to different machines to emulate a joint study of two cohorts that could not be combined. Following the previous work, we used linear regression-based tests for comparison. For all three datasets, we observed a significant reduction in both runtime and communication for SF-GWAS compared to S-GWAS based on the same computing environment (Fig. 2; Methods). SF-GWAS runtimes were consistently an order of magnitude smaller than S-GWAS (e.g., 4.6 hours vs. 64.3 hours for AMD data, representing a 14x reduction). Total communication cost was three to four times lower for SF-GWAS (e.g., 173.7 GB vs. S-GWAS’ 666.6 GB for AMD data), largely due to the requirement of S-GWAS to share the entire encrypted dataset among the parties (Fig. 2), which SF-GWAS circumvents with the federated approach. The output of SF-GWAS closely matched a direct analysis of the pooled plaintext data as expected (Supplementary Fig. 2). We also note that SF-GWAS ensures a 128-bit security level, considerably higher than the 30-bit security provided by S-GWAS (Supplementary Note 7).

**Figure 2.**
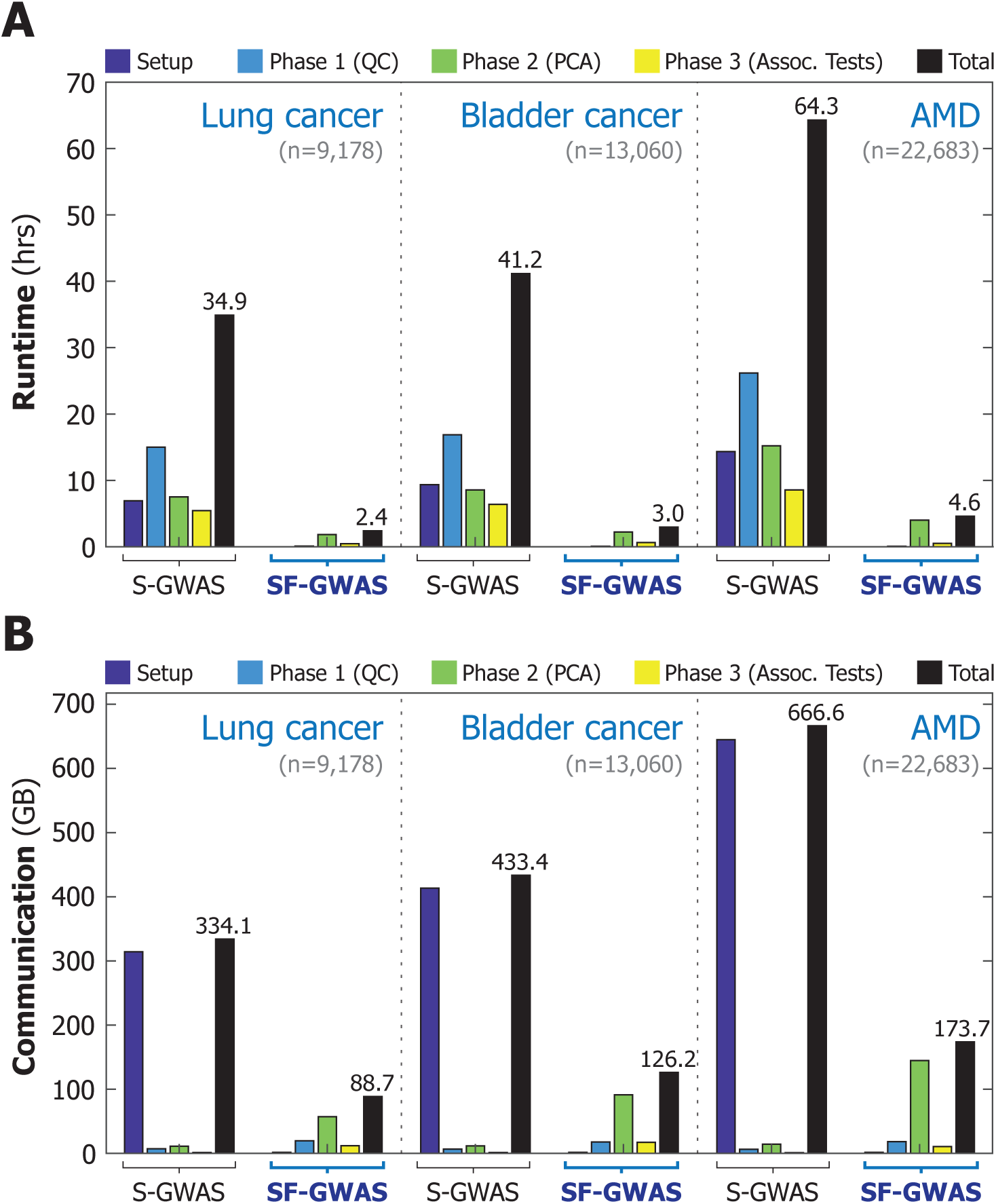
SF-GWAS is more computationally efficient than the prior art for secure collaborative GWAS. We compare the runtime (**A**) and communciation cost (**B**) of SF-GWAS (PCA-based) with those of Secure GWAS^4^ (S-GWAS). Unlike other existing cryptographic solutions for GWAS, these two methods implement the full pipeline including quality control (QC) and principal component analysis (PCA), both of which are standard steps of GWAS. We analyzed the three datasets evaluated in the S-GWAS publication for lung cancer, bladder cancer, and age-related macular degeneration (AMD) using linear regression-based association tests, following the previous work. Each dataset is evenly split into two parts and distributed between two machines to simulate a collaborative GWAS setting. In addition to the total costs (in black), we show the costs of individual steps, including the initial setup and the three phases (QC, PCA, and association tests). The setup involves key generation and network connection for both methods, and additionally encrypted data transfer (secret sharing) for S-GWAS, which is not required by SF-GWAS due to its federated nature. For all datasets, SF-GWAS reduces the overall runtime by an order of magnitude and reduces the communication by a factor of around 3.5. We also note that SF-GWAS provides a 128-bit security level, considerably higher than the 30-bit security of S-GWAS (Supplementary Note 7). Plots showing the accuracy of SF-GWAS results are provided in Supplementary Fig. 2. Unlike S-GWAS, SF-GWAS supports both linear and logistic regression-based tests, even for large-scale datasets. Supplementary Figs. 11 and 12 depict SF-GWAS’s runtime, communication cost, and accuracy for the latter.

To further illustrate the scalability of SF-GWAS, we evaluated it on two biobank-scale datasets: the eMERGE consortium and UK Biobank (UKB). The available portion of the eMERGE dataset included 31,293 individuals and 38 million imputed SNPs, which we split across seven centers according to the data providing organization of each sample, and considering each center-specific dataset as local and private. Note that this dataset is more than a hundred times larger than the largest dataset analyzed in the S-GWAS publication (the AMD dataset), which included 509K array genotypes. For UKB, we divided the largest cohort in UKB of European descent (n=275,812 after removing related individuals) into six datasets of varying sizes (n=21K to 67K) corresponding to different geographic regions from which the individual samples were collected (Methods). Each sample in UKB included 93 million imputed SNPs, representing another 20-fold increase in overall dataset size compared to eMERGE (and 2000x larger than the S-GWAS AMD data). S-GWAS could not be evaluated on either dataset due to its infeasible runtime requirement, estimated to be several months for eMERGE and several years for UKB. For both datasets, we ran the PCA-based pipeline of SF-GWAS to identify genetic associations for body mass index (BMI), over a network of seven virtual machines for eMERGE and six for UKB (plus an auxiliary machine for facilitating the computation; Methods), each holding a private dataset corresponding to a single data center.

Consistent with our previous results, the association statistics computed by SF-GWAS closely matched a plaintext analysis based on the PLINK software^13^ on each of the pooled datasets (Fig. 3 and Supplementary Fig. 3; Methods). In addition, we observed that a meta-analysis of summary statistics from individual centers can result in considerable discrepancies compared to the pooled analysis (Supplementary Fig. 4), especially on the eMERGE dataset, likely due to heterogeneity of study populations and moderate sample sizes. Since the computational steps of SF-GWAS closely emulate a centralized analysis based on a pooled dataset, this ensures that SF-GWAS results are virtually equivalent to what the collaborators would obtain if the datasets could be directly combined for a joint analysis. This equivalence holds regardless of how the data is split, e.g., even when the data distribution is heterogeneous among the parties—a challenging setting for collaborative analyses based only on summary statistics^14–16^.

**Figure 3.**
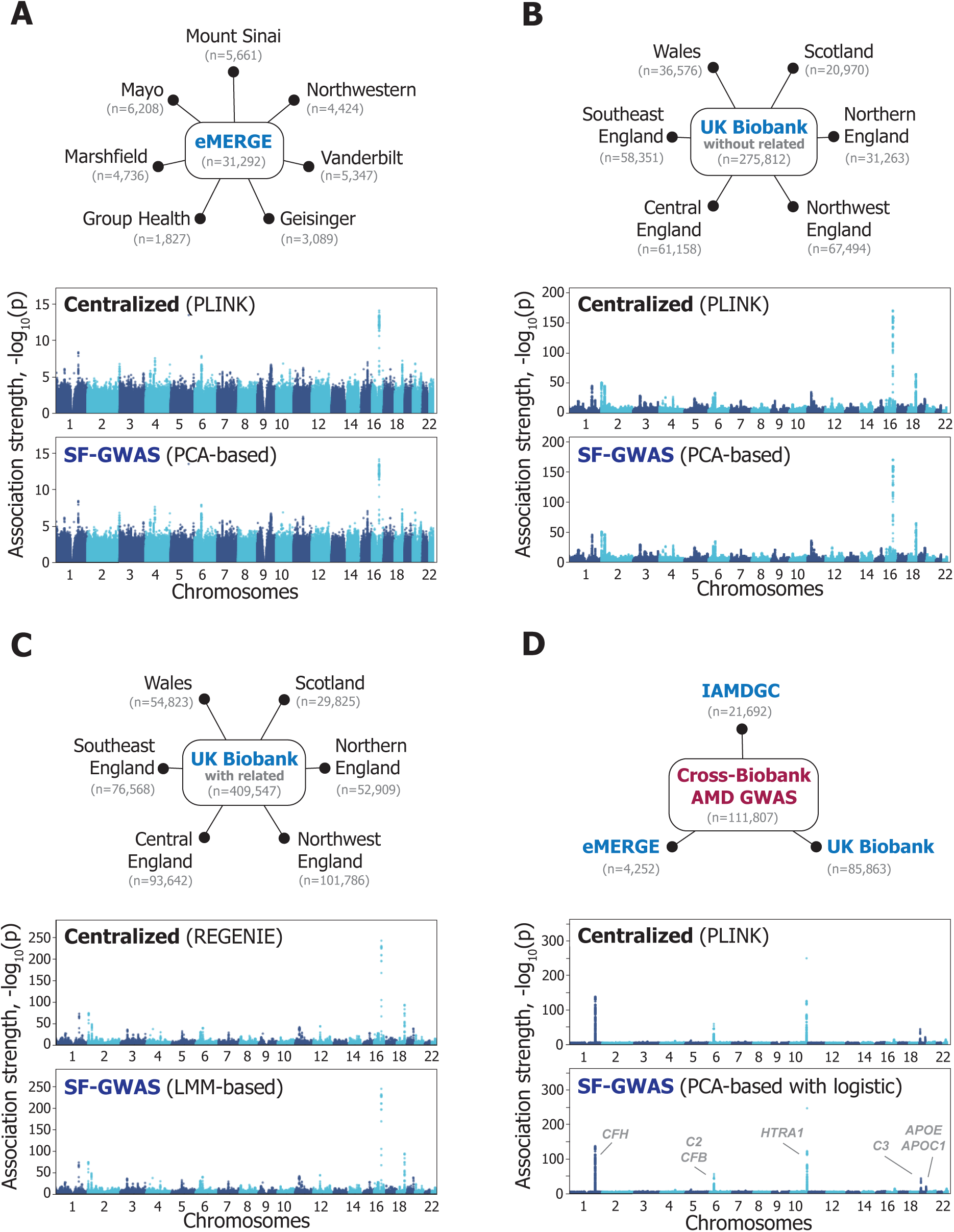
SF-GWAS accurately reproduces biobank-scale GWAS without data centralization. We evaluated SF-GWAS on eMERGE (**A**), UK Biobank (UKB) (**B**, **C**), and cross-biobank AMD (age-related macular degeneration) GWAS (**D**) datasets to demonstrate its applicability to biobank-scale collaborative GWAS. Considering both the number of individuals and the number of variants (Methods), eMERGE is at least 100x and both UKB and cross-biobank AMD are at least 2000x larger than the largest dataset considered by the prior work S-GWAS^4^. The total sample count and the sizes of individual datasets used in the federated setting are shown (left); we split eMERGE data into seven groups according to the data providing organization, and for UKB, we geographically grouped the original sample collection sites into six groups. Cross-biobank AMD represents a setting where joint analysis is performed over a heterogeneous collection of independently collected GWAS datasets. Following standard practice, we excluded individuals with a close relative in the dataset for PCA-based GWAS (**B**), whereas the full cohort was considered for the LMM-based pipeline (**C**). For eMERGE and UKB (**A**-**C**), we assessed the genetic associations of body mass index (BMI), accounting for covariates (age, sex, and center) and principal components, where the latter are globally computed over the entire dataset (securely performed in SF-GWAS). For cross-biobank AMD (**D**), we performed a logistic regression-based case-control association analysis of AMD with the same covariates. Manhattan plots visualizing the association strength of individual variants across chromosomes 1-22 are shown for a centralized, unencrypted analysis using the PLINK or REGENIE software (for PCA-based and LMM-based workflows, respectively) and a secure and federated analysis using SF-GWAS. In all experiments, SF-GWAS accurately reproduces the corresponding centralized analysis without requiring the collaborating entities to share private data. Scatter plots comparing the association statistics are provided in Supplementary Figs. 3 and 11.

The total runtime of SF-GWAS on eMERGE was 17.5 hrs, including 2.8 hrs for quality control filtering (QC), 8 hrs for PCA, and 6.7 hrs for association tests. For UKB, the runtime was 5.3 days in total, including 4.5 hrs for QC, 44 hrs for PCA, and 77.8 hrs for association tests. These runtimes are the first that are practically feasible and represent a major improvement over the prior art. A concurrent work for multi-site GWAS^17^ reported a runtime of 5 hours for 160K SNPs and 16K samples, which extrapolates to more than four months for the UKB dataset, while providing considerably weaker security guarantees than SF-GWAS and addressing only the PCA-based workflow. Since the runtime of SF-GWAS’ PCA-based pipeline scales linearly with the number of individuals and genetic variants, the expected runtime can be readily estimated in practice for different dataset sizes based on our results (Supplementary Fig. 5; Supplementary Note 8). We also expect SF-GWAS to remain practical for international collaborations with higher network communication delays (e.g., between the U.S. and the UK; Supplementary Fig. 6).

The novel LMM-based workflow of SF-GWAS builds upon the plaintext algorithm of REGENIE^18^, a recently developed method for LMM association tests (Supplementary Fig. 1). We make the key observation that the scalable design of REGENIE, based on stacked ridge regressions, together with our federated computational framework, newly allows the design of a practical secure protocol for collaborative LMM analysis. Our algorithm emulates REGENIE in the federated setting while keeping each of the input datasets provably confidential. We devise secure and federated solutions for both ridge regression and linear systems of equations through distributed and iterative optimization techniques that we newly leverage to ensure our algorithm scales to large datasets (Methods; Supplementary Note 6).

We evaluated our LMM-based workflow on the largest dataset of 409,548 individuals of European descent from UKB including related individuals. We analogously split the data across six geographic centers with the size of individual cohorts varying from 30K to 102K and analyzed associations with BMI (Methods). SF-GWAS produced association statistics accurately matching those of REGENIE, where the latter was directly run on a pooled dataset without any encryption (Fig. 3 and Supplementary Fig. 3). We also validated the accuracy of our LMM algorithm on the lung cancer GWAS dataset from the S-GWAS publication (Supplementary Fig. 7). Owing to our key algorithmic optimizations, the LMM-based SF-GWAS exhibits near-constant scaling of runtime in the size of local cohorts (Supplementary Fig. 8 and Supplementary Note 6) and maintains runtimes on the order of days for the large datasets including hundreds of thousands of individuals, realizing a practical runtime of 6 days for the UKB dataset in our experiment (Supplementary Note 8).

Collaborative analysis performed using SF-GWAS identified genetic variants with statistically significant association with BMI that are concordant with prior GWAS results. Comparing with the published summary statistics from the Pan-UK Biobank project^19^, we observed that 71 out of 73 significant loci (*p* < 5 *×* 10*^−^*^8^) identified by SF-GWAS on eMERGE coincided with previously reported associations. When we analyzed each center’s dataset independently, only one out of seven centers resulted in non-zero (two) significant loci, illustrating the benefit of collaboration in agreement with the goal of consortium-based projects like eMERGE. Similarly for UKB, 1,778 out of 2,200 significant loci for LMM-based SF-GWAS, and 21,544 out of 24,357 for our PCA-based analysis on a larger set of imputed genotypes, were previously reported to be significant, indicating a large overlap despite differences in the analysis setting. In contrast, independently analyzing each center’s private dataset led to 2,600 significant loci across all centers, considerably fewer than SF-GWAS (24,357). Moreover, given eMERGE as a study cohort and UKB as a larger validation cohort, meta-analyses of seven center-specific GWAS based on the eMERGE data resulted in fewer loci that are validated in UKB than SF-GWAS, illustrating the potential of the joint analysis enabled by SF-GWAS to increase statistical power (Supplementary Fig. 9).

GWAS analysis of binary traits (e.g., disease status) is often performed using logistic regression, requiring an iterative procedure to estimate model coefficients in the absence of an analytical solution. However, due to the cost of nonlinear operations under encryption, existing works on secure GWAS have either focused on linear regression models or provided only limited support for logistic tests, restricted to small datasets (e.g., requiring 10 days to analyze 500K variants in 100K individuals^7^). Illustrating the broad applicability of our methodology, SF-GWAS also provides a workflow for logistic tests in PCA-based GWAS, which efficiently scales to biobank-scale data. To achieve this result, we leverage a key insight that score-based tests can be efficiently performed in a secure federated setting by requiring only the shared covariate-only model (i.e., the null model) to be fitted using logistic regression while simplifying the remaining computations to those similar to the linear setting (Methods). Furthermore, we developed a secure federated Newton’s method for fitting the covariate-only model, thereby ensuring fast convergence to an accurate solution while significantly outperforming prior secure solutions for logistic regression based on first-order gradient-based methods^4, 20^ (Methods; Supplementary Figure 10). Reanalyzing the three S-GWAS datasets using logistic association tests, as newly enabled by SF-GWAS, we obtain association statistics that are consistent with the centralized analyses using a standard implementation of logistic tests in PLINK (Supplementary Fig. 11). SF-GWAS analyzed all three datasets in less than 5.3 hours (Supplementary Fig. 12), comparable to the runtime costs of linear association tests (Figure 2). Computational costs scale linearly in both settings with respect to data dimensions (Supplementary Fig. 13).

Finally, we demonstrate an application of SF-GWAS to analyzing datasets collected independently by different organizations. In addition to the AMD GWAS cohort from the International AMD Genomics Consortium (IAMDGC; n=21,692, including 9,284 cases), we selected ancestry-matched (European) AMD cohorts in the eMERGE consortium (n=4,252, including 574 cases) and the UK Biobank (n=85,863, including 2,632 cases). This resulted in a GWAS dataset of 111,807 individuals in total, split among three parties representing the original data sources. Although the three cohorts were genotyped using different array platforms, a large number of shared genetic variants (22,191,946) could be chosen for joint analysis by imputing each dataset and finding an intersecting set of genomic positions and allele definitions. The total runtime of SF-GWAS (PCA-based, logistic) was three days, including 53 minutes for quality control (QC) filtering, 8 hours for PCA, and 68 hours for association tests (Methods). For comparison, we estimate a runtime of 42 days for an existing cryptographic solution for GWAS^7^ (which does not support QC or PCA), assuming linear scaling with both the number of computing cores (inversely) and the number of variants. Consistent with previous experiments, SF-GWAS obtained accurate results compared to the results of a centralized analysis on a pooled dataset (Fig. 3D and Supplementary Fig. 11). Top five genomic loci with the strongest associations identified by SF-GWAS correspond to genes with well-established roles in AMD, including *CFH*, *C2*/*CFB*, *HTRA1*, *C3*, and *APOE*/*APOC1* in chromosomes 1, 6, 10, 19, and 19, respectively. These results indicate that SF-GWAS can enable the analysis of large-scale datasets across independent repositories or biobanks.

There are several factors to consider when using SF-GWAS to support real collaborative studies. Data harmonization steps, such as identifying a common set of genetic variants and unifying allele or phenotype definitions, must be performed cooperatively prior to SF-GWAS execution. These steps do not require sharing sensitive raw data and can be achieved by communicating only non-private metadata (e.g., a list of genomic positions) between the parties, as illustrated in our cross-biobank experiment. Furthermore, SF-GWAS may improve data quality by allowing QC filters to be applied globally; for instance, some variants that are discarded due to their rarity at individual sites could be rescued and analyzed by SF-GWAS by jointly considering all datasets. Regarding computational infrastructure needs, our open-source software is designed to run in any environment with minimal dependency. Our sfkit web server (https://sfkit.org) further streamlines the deployment of SF-GWAS and related tools with the option to use either cloud computing resources or the users’ private machines^21^. Although establishing network connectivity between parties may require organizational approval in highly restrictive environments (e.g., hospitals), the formal security guarantees of SF-GWAS, together with its federated analysis approach where the shared encrypted data encode only aggregate-level information, can facilitate this process by minimizing risks and supporting regulatory compliance^22^.

In summary, our work demonstrates a secure federated approach to multi-site GWAS, leading to the first scalable algorithms for conducting collaborative studies without the need to share private data. We demonstrated the practical performance of our methods on five datasets of varying sizes, including the analysis of a large biobank dataset with 410K individuals as well as a cross-biobank analysis involving independent data sources. Further extending our techniques to support other useful analysis pipelines, including logistic mixed-effects models and statistical corrections for imbalanced datasets^23, 24^, and to provide security against a broader set of adversaries, including malicious actors who may deviate from the given protocols, are meaningful directions for future research. We expect our algorithmic strategies and the open-source software library, including modular implementations of a range of GWAS workflows, to accelerate future method development efforts. Our work provides the tools needed to broaden cross-institutional collaboration involving sensitive genomic data, which is key to future progress in biomedicine.

## Methods

### Review of secure multiparty computation (MPC)

MPC techniques enable multiple entities to securely and interactively perform computation on private inputs (Supplementary Note 1). Standard MPC frameworks^25^ leverage (additive) secret sharing, where each private value is split into random (encrypted) shares, which are in turn distributed to different computing parties. While the shares collectively encode the private value, any subset of shares provably does not leak any private information. Computing parties then collaborate and use the secret shares to evaluate a function on the private input without revealing information about the private input to any entity involved. For example, the secure addition of two secretly shared numbers *x* and *y* can be executed by having each party add their individual shares for *x* and *y*. The new shares constitute a sharing of *x* + *y*, which is the desired computation result. More sophisticated functions (e.g., multiplication, division, square root, and sign) can be similarly defined over the secret shares, but require the two parties to interact by exchanging a sequence of numbers, which also do not reveal private information. These secure routines can be composed to perform arbitrary operations on private input data held by multiple entities. However, the communication cost of MPC can introduce a bottleneck in applications involving complex tasks. In addition, secret sharing requires that the entire input data be shared with all computing parties.

### Review of homomorphic encryption (HE)

HE is a form of encryption that allows for direct computations over encrypted data, without having to decrypt them. Initially developed for limited types/rounds of operations, HE is now widely applicable to many analysis tasks due to the recent introduction of fully HE (FHE) schemes, which include a *bootstrapping* routine to allow an arbitrary number of operations to be performed, and the development of efficient techniques for common scientific operations. For instance, the CKKS scheme^26^ encodes a vector of continuous values in a single ciphertext and is well-suited for calculations where a small amount of noise can be tolerated. Like other HE schemes, CKKS performs both additions and multiplications simultaneously on the encrypted values within a ciphertext (single instruction, multiple data [SIMD] property), which improves the overall scalability of the scheme. While HE uniquely enables a single party to perform computation on the encrypted data without interaction, the computational cost and flexibility of HE remain more limited than MPC for general tasks. Also, for multi-site collaboration, one needs to transfer all of the encrypted data to a single machine for joint analysis, which can be challenging for large datasets.

### Our approach: combine HE and MPC to enable practical, secure federated computation

To address the performance limitations of HE and MPC, we take a novel federated approach to secure computation leveraging both HE and MPC, where the input datasets are kept locally at the respective source sites and only small intermediate data are securely exchanged among the parties using cryptographic techniques to carry out a global computation. For the HE component, we build upon a multiparty HE (MHE) scheme (related to threshold HE) based on CKKS, which extends the CKKS scheme to the setting with multiple data providers by secret-sharing the decryption key and constructing a shared encryption key (Supplementary Note 2). Under our scheme, any party can encrypt the data and perform HE computations locally, but decryption can be performed only if all parties cooperate. Our approach allows each party to perform local computations involving the unencrypted input dataset and a small amount of encrypted data, whereas certain global computations are performed by sharing intermediate results among the parties, encrypted under the shared encryption key. At the end of the protocol, all parties collaborate to decrypt the final results. By keeping each input dataset local, we minimize the communication and enable local plaintext computation, which are significantly more efficient than corresponding ciphertext computation. We also leverage the fact that an efficient *interactive* protocol for bootstrapping exists in MHE^27^ to reduce the overall computational burden of HE.

Improving upon existing works on MHE^5, 28–30^, we switch between the MHE and secret sharing representations of intermediate data, which enables efficient state-of-the-art MPC routines to be used in conjunction with MHE operations to carry out the global computations (Supplementary Note 3). We convert between the two schemes to perform key operations under the most efficient scheme for each of the computational steps of the GWAS pipeline. For example, we perform large-scale matrix and vector operations using MHE to exploit its SIMD (single instruction, multiple data) property, but evaluate non-polynomial functions (division, square root, and comparison) with compact bit-wise MPC protocols, which are more efficient and numerically stable than the MHE counterparts. Any operation involving the local unencrypted data is performed using MHE to avoid secret sharing of the large input datasets.

### Our algorithmic design strategies for enabling secure population structure correction

Our federated framework for secure computation allows us to develop efficient and provably secure algorithms for collaborative GWAS. In another key advance beyond prior work, we introduce practical methods for two standard approaches to account for population structure, namely, principal component analysis (PCA) and linear mixed models (LMMs). We adopt the following algorithmic design strategies to obtain accurate and efficient performance in a secure setting. First, we closely adhere to the highlevel computational pipeline of the desired centralized algorithm to obtain accurate results, while using optimized cryptographic routines to securely and jointly operate over private datasets held by multiple parties. This is in contrast to other collaborative approaches that simplify or approximate the analysis to address the lack of access to a pooled dataset (e.g., meta-analysis). Next, we restructure the algorithm while maintaining its equivalence to the original computation to both maximize the use of low-cost operations and minimize communication by leveraging local plaintext data. We switch between MPC and MHE routines throughout the protocols to improve the efficiency and numerical robustness of secure computation routines. We also optimize the vectorized encoding of data in encrypted representations for efficient composition of linear algebra operations. We detail our algorithmic strategies and optimization techniques in Supplementary Notes 4-6.

On top of enabling a significant performance improvement for PCA compared to prior work^4^, our techniques allowed us to design the first practical protocol for secure and federated LMMs (Supplementary Fig. 1; Supplementary Note 6). LMMs typically require operations involving the genetic relatedness matrix (GRM), which scales with the number of individuals in the dataset and imposes a heavy computational burden for large cohorts even in a centralized analysis setting. Our new approach builds upon REGENIE^18^, a recently developed algorithm for LMM association tests based on a stacked ridge regression approach, which directly models the ancestry-related confounding effect as the output of a genome-wide regression model, thus circumventing the use of a GRM. This approach brings substantial scalability improvements while providing accuracy comparable to other LMM methods such as BOLT-LMM^31^, fastGWA^32^, and SAIGE^33^. But unfortunately REGENIE cannot be directly used in a secure federated setting. Although ridge regression can be efficiently performed in plaintext on a pooled dataset, implementing standard algorithms (e.g., based on the closed-form solution) with secure computation techniques leads to impractical runtime requirements due to the complex matrix operations (e.g., inversion) that need to be performed on large encrypted matrices. We overcome this challenge by comprehensively redesigning REGENIE’s stacked regression procedure with novel, secure federated algorithms for ridge regression based on conjugate gradient descent (CGD) and alternating direction method of multipliers (ADMM). Although the inclusion of private covariate features renders a direct application of the latter approach^34^ impractical in our setting, our reformulation of the ADMM algorithm (referred to as ADMM-Woodbury) uses a matrix identity to enable plaintext precomputation of costly operations (e.g., large-matrix inversion) at each computing node; thus, our approach maximizes efficient local computation and, as a result, obtains near-constant scaling of runtime with respect to cohort size (Supplementary Fig. 8).

### Our scalable techniques for secure association tests based on linear and logistic models

Performing association tests at genome-wide scale involves intricate linear algebra operations (e.g., projection, inversion, and factorization) applied to large-scale matrices and vectors. Following the strategies in the previous section, we designed efficient algorithms for secure federated association tests that can analyze millions of genetic variants within practical runtimes. For example, we developed optimized routines for matrix multiplications between the local genotype matrix and an encrypted low-dimensional matrix (or a vector) of covariates and phenotypes by identifying and precomputing intermediate terms that involve the unencrypted local dataset, which are more efficiently computed than operations involving only encrypted data. This approach also helps to distribute the workload among the parties according to the size of the local datasets. For complex, iterative operations such as matrix factorization and inversion, we securely convert between MHE and MPC frameworks to take advantage of efficient MPC routines for costly operations such as comparison and division. Furthermore, after the initial precomputation, we process genetic variants in non-overlapping genomic blocks in parallel to accelerate the computation.

Prior to our work, logistic regression-based association tests were considered more challenging and could be performed using cryptographic techniques only on small datasets. This limitation has mainly been due to the costly iterative optimization procedure required to estimate the coefficients of logistic regression models (e.g., genetic effect sizes). To address this challenge, we developed an efficient secure federated algorithm for score-based tests (Supplementary Note 5). Our algorithm securely obtains statistics that are asymptotically equivalent to alternative tests (i.e., Wald and likelihood-ratio) in a manner that efficiently scales to large biobank datasets. Unlike the Wald or likelihood-ratio tests, both of which require a separate logistic regression model to be fitted for every variant being tested, score-based tests fit a single null model including only the covariates, subsequently allowing all variants to be tested using a closed-form expression without refitting the model. To obtain the initial null model efficiently and accurately, we devised a federated algorithm for Newton’s method harnessing both MHE and MPC for efficient matrix multiplications and Hessian matrix inversion, respectively. This approach enables faster convergence to precise estimates compared to standard first-order gradient-based methods (Supplementary Fig. 10). Once the null model is obtained, the computation of association statistics largely reduces to linear algebraic operations, for which our secure federated approach provides a scalable computation protocol. As our results demonstrate, these novel techniques enable SF-GWAS to obtain accurate logistic regression-based association statistics within practical runtimes that are comparable to the linear regression setting.

### Secure federated GWAS (SF-GWAS) pipelines

Our SF-GWAS algorithm implements the full GWAS pipeline, including quality control (QC), correction for population structure (PCA and LMM), and association tests. Collaborating parties first agree on the phenotype, covariates, and a list of genetic variants to analyze, as well as study parameters, such as filtering thresholds and other algorithmic parameters. They also agree on the security parameters and generate the required cryptographic keys for HE and MPC, e.g., encryption keys and shared pseudorandom number generators (Supplementary Notes 1 and 2). They then proceed with the interactive protocol to securely perform the desired computation. For QC, the parties independently filter their subset of samples based on public thresholds (e.g., for heterozygosity and missing genotype rate), then utilize MPC routines to jointly and securely filter the variants based on global statistics (e.g., minor allele frequencies and Hardy-Weinberg equilibrium). The variant filtering output is shared with all parties so that the rest of the protocol can proceed with the reduced variant set.

For the PCA-based workflow, we implemented a secure federated algorithm for randomized PCA^4, 35^ to compute the top principal components (PCs) over the entire dataset without constructing the pooled matrix (Supplementary Note 5). The jointly computed PCs are kept encrypted for use in the following steps. For linear regression-based association tests, we use both HE and MPC operations to compute the covariate-corrected association statistics based on a linear model. This step includes: (i) a secure federated QR factorization for computing the joint orthogonal basis of PC and the observed covariates; (ii) secure matrix multiplication based on HE to project the genotypes onto the covariate subspace for correction and to compute genotype-phenotype covariances; and (iii) MPC routines to inversely scale the statistics by the standard deviations of the genotypes and the phenotype for normalization. For logistic regression-based association tests, we use our secure federated algorithm for score-based tests (Supplementary Note 5). The first step is to fit a null logistic regression model across the parties considering only the covariates, for which we introduce a federated Newton’s method based on both HE and MPC. Then, we compute the components of the score-based test statistic with respect to each tested variant, which include (i) the score (i.e., the derivative of the log-likelihood function with respect to the coefficient of the genetic effect) and (ii) the Fisher information (related to the standard error of the coefficient). Both terms and the resulting statistics are computed securely in a federated manner using a series of linear algebra operations in HE and nonlinear operations in MPC. Finally, the association statistics are collectively decrypted and shared among the parties as the final output of the analysis.

For the LMM-based workflow, we implemented a secure federated version of REGENIE^18^ that is based on stacked ridge regression models (Supplementary Note 6). After the QC step, the genetic variants are first grouped into fixed-size blocks. For each block, a ridge regression model is jointly trained across the parties using our ADMM-Woodbury algorithm to obtain encrypted phenotype predictions, which leverage only the variants within the block while accounting for linkage disequilibrium. Subsequently, the block-wise local predictions are provided as input features to a genome-wide ridge regression model for phenotype prediction, jointly trained across the parties using our CGD algorithm. We adopt a cross-validation scheme (CV) to determine an appropriate choice of variance parameter, representing the genomic heritability of the target phenotype. We selected K-fold cross validation as it is typically more efficient than other methods such as leave-one-out cross-validation and achieves almost identical accuracy in this framework^18^. Association tests are performed by calculating the correlation between each target variant and the phenotype residuals excluding the contribution from the genome-wide regression model, estimated without the chromosome including the tested variant. Analogous to the PCA-based workflow, we efficiently compute the association statistics using optimized secure matrix multiplication routines along with MPC routines for data normalization.

We detail the security properties of SF-GWAS workflows in Supplementary Note 7 and provide runtime and complexity analyses in Supplementary Note 8 and Supplementary Tables 2–5.

### Related work

Several works have proposed methods to securely perform collaborative GWAS over private datasets using secure computation frameworks (HE or MPC)^4, 5, 7^, but are limited by runtimes or accuracy of GWAS analysis without correction for population structure. As noted, MPC frameworks based on secret sharing^4^ are costly with respect to communication between parties and require the entire dataset to be encrypted and distributed among all parties. Although HE enables non-interactive computation over encrypted data, joint analysis based on HE^7^ still requires the collaborating entities to centralize the encrypted data for a single party to analyze, which leads to impractical communication costs for large-scale genetic datasets and significant computational overhead for performing complex analysis tasks. Prior work applying multiparty HE (MHE) to GWAS^5^ addresses the limitations of centralized HE for only association testing and none of the other essential components of GWAS, i.e., quality control filtering and correction for population structure (PCA or LMMs). Algorithms for PCA and LMMs involve highly sophisticated computational pipelines, presenting a major challenge for existing cryptographic tools for secure computation in obtaining practical performance. Our work overcomes this challenge to realize scalable and provably-secure algorithms for the full GWAS pipeline. In addition, this earlier work solely relied on HE computations and did not consider jointly leveraging MPC protocols (as we do in our work) to obtain robust accuracy across a wide range of datasets and improved efficiency for non-polynomial operations. Secure hardware-based approaches to GWAS (e.g., based on Intel SGX) have also been proposed^6^; however, existing methods are limited to association testing only (without PCA or LMMs) and do not provide formal guarantees of privacy like HE or MPC, leading to known security risks^36–38^.

Another line of work for collaborative GWAS is based on federated learning techniques, which iteratively aggregate intermediate results among the parties to carry out global computations^16, 39^. These solutions are generally more accurate than meta-analysis as they more closely emulate a pooled analysis^16^. They also achieve efficient performance in general because the plaintext (unencrypted) data are locally held and directly analyzed by each party. However, these methods require intermediate results to be shared among the parties (or with a trusted third-party) in plaintext, which may lead to private data leakage^40, 41^. Although differential privacy techniques can be employed to mitigate such leakage, existing methods do not provide a practical solution for GWAS where releasing the statistics for a large number of variants would require a significant amount of noise to be added for privacy^42^. In SF-GWAS, the parties keep their local data in plaintext and exchange only intermediate results that remain encrypted throughout the study; our approach efficiently emulates a pooled analysis while ensuring a formal notion of privacy during the entire process.

### Benchmark datasets

We obtained the three datasets used in the original S-GWAS publication^4^ for comparison. These include a lung cancer dataset (n=9,178, including 5,088 cases; 612,794 SNPs), a bladder cancer dataset (n=13,060, including 9,684 cases; 566,620 SNPs), and an age-related macular degeneration (AMD) dataset (n=22,683, including 6,211 cases; 508,740 SNPs). We followed the steps in the prior work to prepare the data, then evenly and uniformly split each dataset between two parties to emulate a multi-site study.

For the eMERGE data, we obtained a cohort of 31,292 individuals split across seven study groups: Geisinger Health System (n=3,089), Group Health (University of Washington; n=1,827), Marshfield Clinic (Pennsylvania State University; n=4,736), Mayo Clinic (n=6,208), Icahn School of Medicine at Mount Sinai (n=5,661), Northwestern University (n=4,424), and Vanderbilt University (n=5,347). We used a total of 38,040,168 imputed biallelic SNPs for the analysis and chose body-mass index (BMI) as the target phenotype. Four covariates were included in the analysis: membership to each study group, age (at time of assessment), sex, and age^2^.

For the UK Biobank (UKB) data, we obtained a cohort of 409,547 individuals of European descent. For use with the PCA-based GWAS pipeline, we also constructed a subset of 275,812 unrelated individuals (King^43^ relatedness coefficient less than 0.062). This cohort represented 22 different health centers across the United Kingdom. To simulate a federated study, we grouped the health centers into six study groups based on geographic location: Scotland (n=29,825 in total; 20,970 unrelated), Northern England (n=52,909; 31,263), Northwest England (n=101,786; 67,494), Central England (n=93,642; 61,158), Southeast England (n=76,568; 58,351), and Wales (n=54,823; 36,576). We provide the list of centers in each group in Supplementary Table 1. We used a total of 92,248,310 imputed biallelic SNPs for the PCA-based pipeline, and a subset of 581,927 non-imputed genotyped SNPs for the stacked regression models in the LMM-based pipeline, following the recommendation in the REGENIE software documentation^44^. We analyzed BMI as the target phenotype. Six covariates were included in the analysis: membership to each study group, age (at time of assessment), sex, age *∗* sex, age^2^, and age^2^*∗* sex.

For the cross-biobank AMD GWAS analysis, we used data from three independent sources: eMERGE (n=4,252, including 574 cases), UK Biobank (n=85,863, including 2,632 cases), and International AMD Genomics Consortium (IAMDGC; n=21,692, including 9,284 cases) for a total of 111,807 individuals (12,490 cases). We used the imputed genotype data provided by eMERGE and UKB. For IAMDGC, we used Michigan Imputation Server to impute the samples using the Haplotype Reference Consortium reference panel. The total number of imputed biallelic variants shared among the three datasets was 22,191,946. For UKB, AMD cases were ascertained based on the ICD10 code H35.3, and the control samples were randomly sampled from the remaining cohort. For eMERGE and IAMDGC, we retained the AMD case and control labels provided in the original datasets. To demonstrate the scalability of our approach, we considered individuals of European ancestry in each dataset, determined based on self-reported ethnicity for UKB and GrafPop-inferred ancestry for eMERGE and IAMDGC (*P_e_* > 90%). Four covariates were included in the analysis: membership to each study group (two independent features), age (at time of assessment), and age^2^. Other covariates of potential interest, such as sex, were not available in all datasets and thus were excluded from the analysis.

### GWAS details

For the lung cancer, bladder cancer, and AMD datasets, we used the same quality control (QC) filters as applied in the prior analysis of these datasets using S-GWAS^4^. For the eMERGE data, we used the following QC parameters: genotype missing rate per SNP <0.1, minor allele frequency (MAF) >0.05, and Hardy–Weinberg equilibrium chi-squared test statistic <23.928 (*p*-value >10*^−^*^6^). We used the same set of filters for the UKB and cross-biobank AMD data, except for MAF >0.001 to account for the larger size of these datasets.

For PCA-based GWAS, we adopt the standard approach of using a reduced set of SNPs with low levels of linkage disequilibrium (LD) for the PCA step. SF-GWAS achieves this by imposing a minimum pairwise distance threshold of 100 Kb after QC filtering, which we found to obtain similar results as alternatives based on a direct calculation of LD (Supplementary Fig. 14). To establish parity between the plaintext, centralized analysis and our SF-GWAS approach, we use the same set of SNPs for PCA in both analyses for our main results. Alternatively, SNP selection for PCA can be performed separately and the agreed-upon list of SNPs may be provided as input to SF-GWAS. For lung cancer, bladder cancer, AMD, eMERGE, and cross-biobank AMD datasets, we kept the top 5 principal components as covariates for the subsequent analysis. For UKB, we kept the top 10 principal components. We assess the association between each SNP and phenotype of interest based on both linear and logistic regression models including covariates.

For linear association tests, SF-GWAS first constructs an orthogonal basis (*Q*) for the subspace defined by the covariates (e.g., top principal components, age, sex, study group memberships), then computes the Pearson correlation coefficient (*r*) between the genotype and phenotype vectors where the covariate effects have been projected out using *Q*. The coefficient *r* for each SNP is revealed to the collaborating entities. The corresponding *χ*^2^ statistic with one degree of freedom is obtained as *r*^2^(*n − c*)*/*(1 *− r*^2^), based on which a *p*-value can be calculated. *n* and *c* denote the total number of individuals and the number of covariates, respectively. Note that this mapping does not reveal any additional information other than *r*. For logistic association tests, SF-GWAS computes the score-based test *t*-statistic for each SNP *i* as 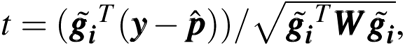 where 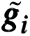 is the genotype residual vector across individuals after adjusting for covariates, ***y*** the phenotype vector, 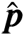 the vector of estimated mean of the trait based on the null model, and 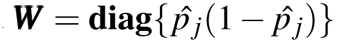 with 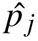 the *j*-th element of 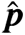 corresponding to the *j*-th individual, following the approach of REGENIE^18^. The corresponding *p*-value is estimated via a normal approximation with 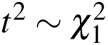.

For LMM-based GWAS, we follow REGENIE’s approach to first apply ridge regression to obtain the best predictive model of the covariate-corrected phenotype within a given genomic block, accounting for genotype correlations, and then perform a second regression to obtain genome-wide phenotype predictions based on the block-wise predictors. The size of each genomic block is set to 8,192 in order to maximally leverage the vectorized encryption scheme based on our cryptographic parameters. Following REGENIE, we use a five-fold cross validation to select the best variance parameter to construct the final predictors. For the association tests, we adopt the standard leave-one-chromosome-out (LOCO) scheme, leaving out the chromosome including the tested variant to correct for the background genetic effect on the phenotype without interfering with the signal being tested. We then obtain the *χ*^2^ statistic with one degree of freedom, analogous to the PCA-based pipeline, using the residuals of the LOCO genome-wide predictors for each variant.

### Evaluation approaches

We evaluated SF-GWAS by simulating each party on a separate virtual machine (VM) with 16 virtual CPUs (vCPUs) and 128 GB of memory (e2-highmem-16) on the Google Cloud Platform (GCP). For the main results, we adopt the most-efficient local area network (LAN) setting by creating the VMs within the same zone in GCP; we illustrate the reasonable additional cost of wide-area network (WAN) setting in Supplementary Fig. 6. For the UKB and cross-biobank AMD analyses, we used the same or larger VM types to utilize more vCPUs in parallel given the large size of the dataset. Specifically, we used n2-highmem-64 (64 vCPUs with 512 GB of RAM) and n2-highmem-128 (128 vCPUs with 864 GB of RAM) depending on the size of the local dataset.

We measured the runtime and communication cost of SF-GWAS (the latter measured by the number of bytes sent among the parties) on all datasets. We compared these metrics against the prior work implementing the analogous PCA-based GWAS pipeline, S-GWAS^4^. We also evaluated SF-GWAS’s scaling with respect to the number of samples, SNPs, covariates, and computing parties (Supplementary Fig. 5 and 13). For the scaling experiments, we replicated the lung cancer dataset to produce a dataset of desired dimensions and modified GWAS parameters as needed to ensure that the amount of data at each step grew proportionally with the input dimensions, e.g., ensuring that the number of samples passing quality control doubles when the original number of samples doubles.

To evaluate the accuracy of SF-GWAS, we compared its association statistics to those obtained from a plaintext, centralized analysis where the individual datasets are combined to form a single consolidated dataset for analysis. For the three S-GWAS datasets, we used a plaintext Python implementation of the same procedure as SF-GWAS (using a standard PCA implementation in the scikit-learn package^45^); and for eMERGE, UKB, and the cross-biobank AMD analysis, we used the PLINK software (https://www.cog-genomics.org/plink/2.0/) implementing the same pipeline for PCA-based GWAS. For LMM-based GWAS, we used the REGENIE software (https://rgcgithub.github.io/ `regenie/`) on the pooled dataset with the same parameters to obtain the ground truth association results. For eMERGE and UKB, we additionally evaluated the accuracy of meta-analysis approaches by performing a separate GWAS for each study group with the same study parameters as the global analysis, and then combining the association statistics among different parties using the meta-analysis methods implemented in PLINK.

## Data availability

The three datasets used for comparison with the prior work on Secure GWAS^4^ are available via NIH dbGaP with accession numbers phs000716.v1.p1 (lung cancer), phs000346.v2.p2 (bladder cancer), and phs001039.v1.p1 (AMD; International AMD Genomics Consortium [IAMDGC]). The eMERGE consortium data is also available via dbGaP (phs000888.v1.p1). Data access applications for the UK Biobank data can be submitted at: https://www.ukbiobank.ac.uk/.

## Code availability

Our open-source software is available at: https://github.com/hhcho/sfgwas. Automated workflows for executing SF-GWAS are available on the sfkit web server (https://sfkit.org), which allows a group of collaborators to easily perform a secure joint analysis of their private datasets using either cloud computing resources or private machines.

## Ethics & Inclusion statement

The use of all controlled-access datasets in this study were approved by the respective data access committees through NIH dbGaP and the UK Biobank Access Management System.

