## Supplementary Materials for "Secure and Federated Genome-Wide Association Studies for Biobank-Scale Datasets"

##### Supplementary Figures


**Supplementary Figure 1**: Overview of SF-GWAS' secure and federated approach to LMM-based association tests.

**Supplementary Figure 2**: SF-GWAS accurately reproduces end-to-end PCA-based GWAS (linear) without data centralization.

**Supplementary Figure 3**: SF-GWAS accurately reproduces biobank-scale GWAS without data centralization.

**Supplementary Figure 4**: Meta-analysis approach for multi-site GWAS often deviates from an ideal centralized analysis.

**Supplementary Figure 5**: SF-GWAS efficiently scales with all dimensions.

**Supplementary Figure 6**: SF-GWAS remains practical for a trans-Atlantic wide-area network setting between the US and the UK.

**Supplementary Figure 7**: SF-GWAS closely reproduces REGENIE's LMM-based association test results on the lung cancer dataset.

**Supplementary Figure 8**: LMM-based SF-GWAS scales efficiently to large datasets with near-constant runtime over a range of dataset sizes.

**Supplementary Figure 9**: Collaborative GWAS of eMERGE data using SF-GWAS leads to a greater number of validated associations than meta-analysis approaches.

**Supplementary Figure 10**: Faster convergence of our secure federated Newton's method for logistic regression compared to standard first-order gradient descent methods.

**Supplementary Figure 11**: SF-GWAS accurately reproduces end-to-end PCA-based GWAS (logistic) without data centralization.

**Supplementary Figure 12**: The runtime and communication costs of logistic regression-based SF-GWAS on the three S-GWAS datasets.

**Supplementary Figure 13**: Efficient scaling of logistic regression-based SF-GWAS.

**Supplementary Figure 14**: Alternative strategies for selecting a reduced set of SNPs for PCA yield similar GWAS results.


##### Supplementary Tables (in this document)


**Supplementary Table 1**: List of UK Biobank assessment centers grouped by geographic region.


##### Supplementary Tables (included as Excel spreadsheets)


**Supplementary Table 2**: Runtime and Communication Costs of Core Operations in SF-GWAS.

**Supplementary Table 3**: Complexity of PCA-based SF-GWAS.

**Supplementary Table 4**: Complexity of LMM-based SF-GWAS.

**Supplementary Table 5**: Complexity of Linear Algebra Routines in SF-GWAS.

**Supplementary Notes**

**Supplementary Note 1**: MPC Definitions and Operations

**Supplementary Note 2**: MHE Definitions and Operations

**Supplementary Note 3**: MHE-MPC Conversion

**Supplementary Note 4**: Algorithm Design Techniques

**Supplementary Note 5**: SF-GWAS Algorithm for PCA-based GWAS

**Supplementary Note 6**: SF-GWAS Algorithm for LMM-based GWAS

**Supplementary Note 7**: Security of SF-GWAS

**Supplementary Note 8**: Runtime Estimation and Complexity Analysis

**Supplementary Note 9**: Derivation of Our ADMM-Woodbury Algorithm

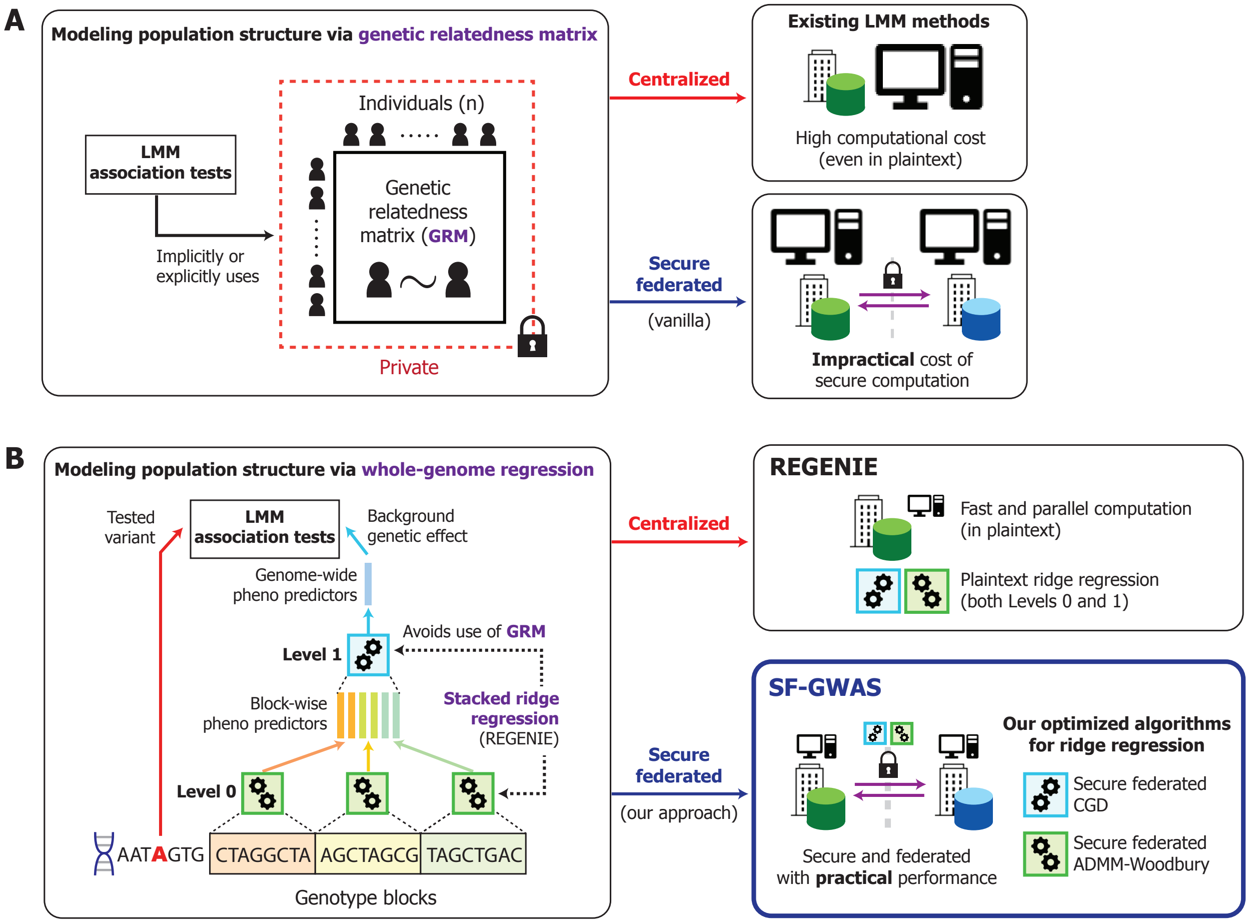


Supplementary Figure 1: **Overview of SF-GWAS' secure and federated approach to LMM-based association tests.** (**A**) Conventional methods for LMM-based association analysis utilize a genetic relatedness matrix (GRM), either explicitly or implicitly, during the computation to model population structure (*left*). This leads to considerable computational costs for large datasets even in a centralized analysis setting (*top right*). Secure and federated execution of such methods with guaranteed privacy for the input data (including the GRM) is infeasible due to the overhead of cryptographic operations (*bottom right*). (**B**) The LMM-based workflow of SF-GWAS draws from the plaintext algorithmic pipeline of REGENIE, a recently proposed scalable approach for LMMs (*left*). As in REGENIE, we adopt a stacked ridge regression approach, where phenotype predictions based only on genetic variants in small genomic regions (blocks) are jointly fed into a genome-wide phenotype prediction model. The output of the whole-genome regression model represents the background genetic effect to correct for in the subsequent association tests. This approach avoids the use of GRMs and thus leads to efficient performance in plaintext (*top right*). To support this workflow for a collaborative GWAS, we introduce efficient, secure and federated algorithms for two different methods for ridge regression (neither of which is used by REGENIE): conjugate gradient descent (CGD) and our improved distributed optimization algorithm called ADMM-Woodbury (*bottom right*). Enabled by our new algorithms, SF-GWAS realizes secure and federated LMM-based association tests with practical performance.

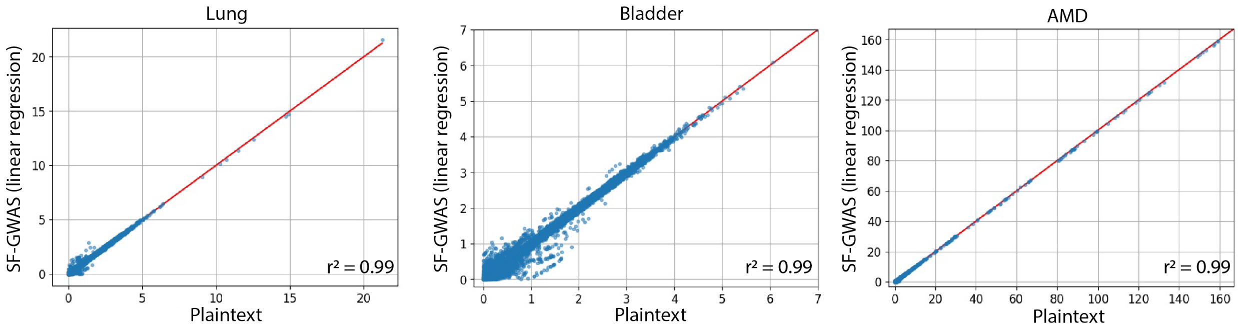


Supplementary Figure 2: **SF-GWAS accurately reproduces end-to-end PCA-based GWAS (linear) without data centralization.** We evaluated PCA-based SF-GWAS (with linear regression) on lung cancer (**left**), bladder cancer (**middle**) and age-related macular degeneration (AMD; **right**) datasets to demonstrate that it obtains similar results as a centralized study in which all plaintext (unencrypted) data are pooled and analyzed together. We evenly split the datasets between two computing parties and compared the covariate-corrected, Pearson correlation coefficients of individual variants obtained using (SF-GWAS) with those from a centralized analysis (Plaintext; using the standard scikit-learn Python library). The analysis pipeline includes both the quality control and PCA steps. Transparency is added to visualize density. While a small amount of accuracy loss can be seen due to the limited precision of cryptographic schemes, note that the squared Pearson correlation coefficients ($r^2$) between the two results are always above 0.98. Results for the same datasets using the logistic regression-based tests are provided in Supplementary Fig. 11.

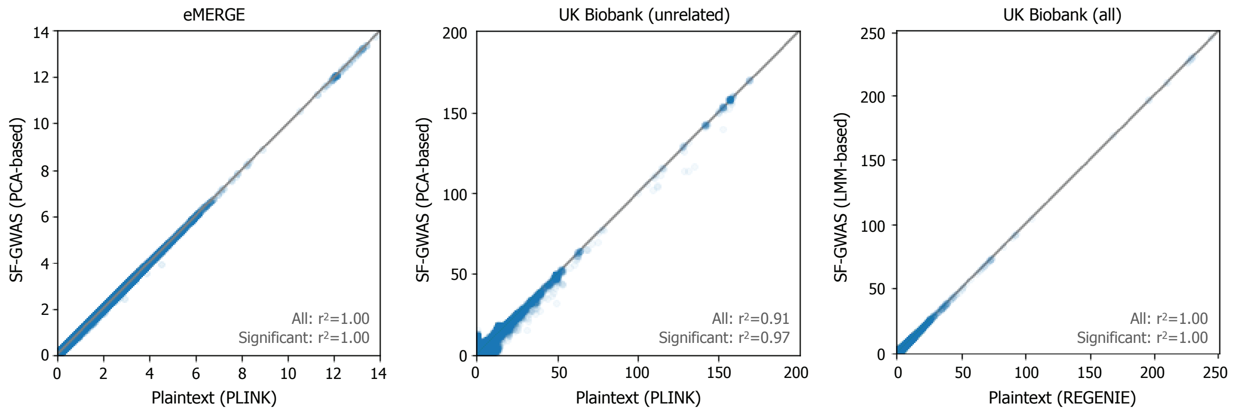


Supplementary Figure 3: **SF-GWAS accurately reproduces biobank-scale GWAS without data centralization.** We evaluated PCA-based SF-GWAS on the eMERGE consortium dataset (**left**), split across 7 data collection centers, and the UK Biobank (UKB) dataset (individuals of European descent, related individuals excluded) (**middle**), split across 6 geographically grouped centers. We additionally evaluated LMM-based SF-GWAS on the UKB dataset including all European individuals, also split across 6 centers (**right**). The plots compare the association statistics ($-\log_{10}(p)$) of individual genetic variants obtained by SF-GWAS to those from a centralized analysis using the PLINK software for the PCA-based analysis and REGENIE for the LMM-based analysis (Plaintext). We show the squared Pearson correlation coefficients ($r^2$) both for all variants and for significant variants identified in the centralized setting (with nominal $p < 5 \times 10^{-8}$). Transparency is added to visualize density. A small amount of numerical noise is attributed to the reduced precision of cryptographic operations. In all datasets, SF-GWAS results closely match the centralized analysis without requiring data centralization.

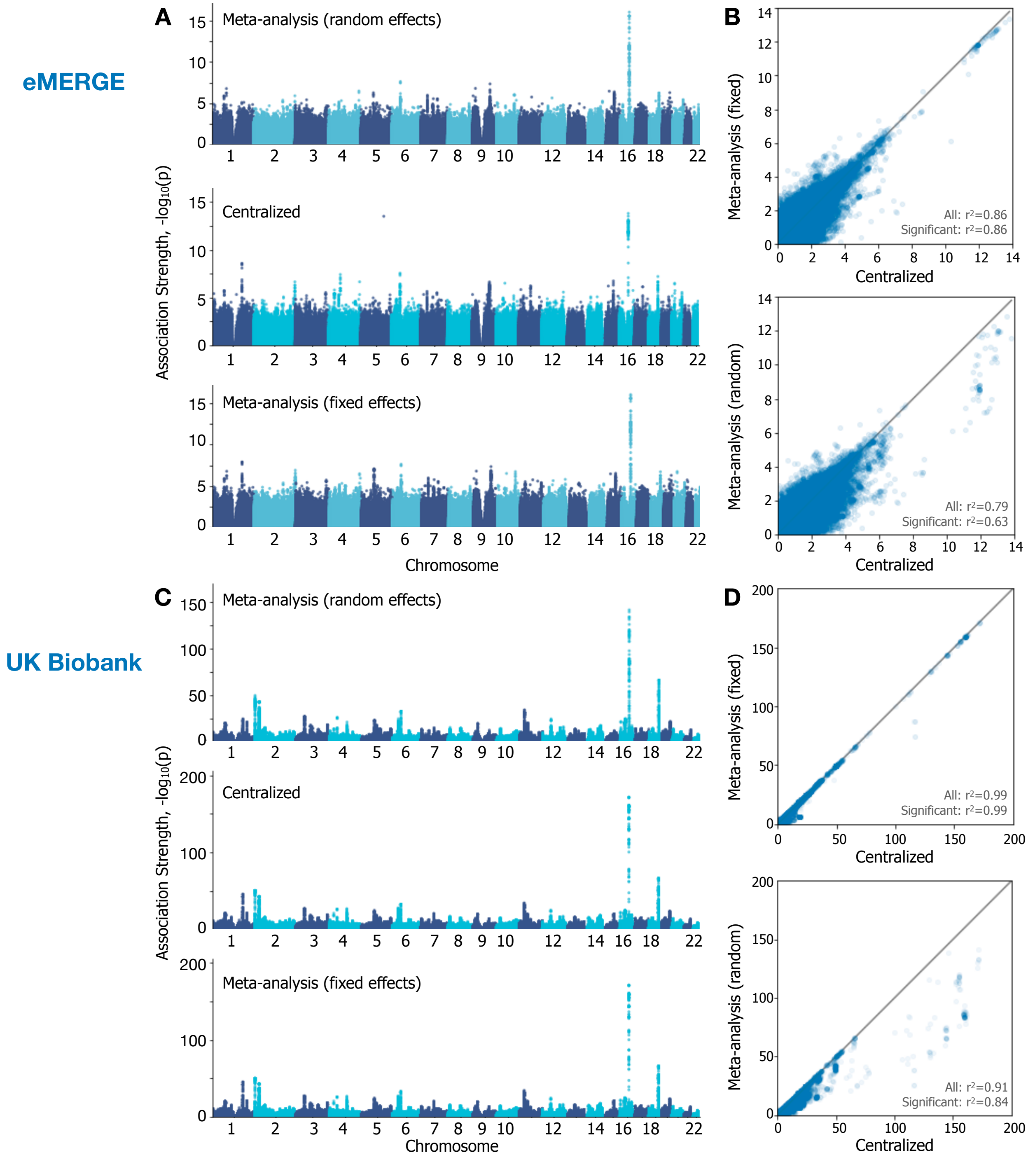


Supplementary Figure 4: **Meta-analysis approach for multi-site GWAS often deviates from an ideal centralized analysis.** Using the same federated study settings from the evaluation of SF-GWAS, we applied standard meta-analysis approaches (i.e., random-effect and fixed-effect methods provided by PLINK) to the eMERGE (**top**) and UKB (**bottom**) datasets based on the PCA-based analysis. We show both the Manhattan plots for individual methods (**A**,**C**) and the scatter plots comparing the association statistics between the centralized analysis and each meta-analysis method (**B**,**D**). Squared Pearson correlation coefficients ($r^2$) both for all variants and for significant variants identified in the centralized setting (with nominal $p < 5 \times 10^{-8}$) are reported. Transparency is added to visualize density in the scatter plots. While meta-analysis results are accurate in UKB based on the fixed-effects model, in all other settings it leads to considerable deviations from the centralized analysis.

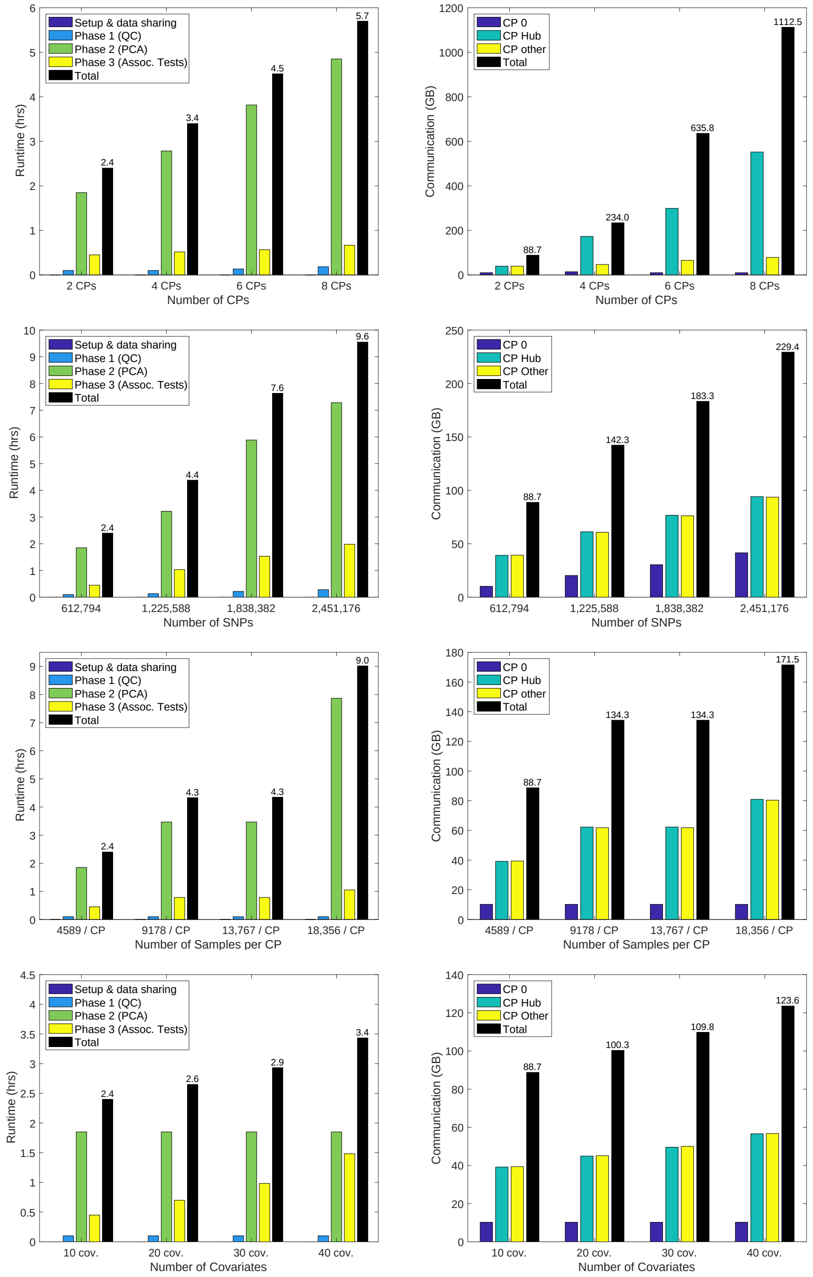


Supplementary Figure 5: **SF-GWAS efficiently scales with all dimensions.** We evaluated how the runtime (**left column**) and communication cost (**right column**) of PCA-based SF-GWAS scale with various parameters of the study setting, including: the number of computing parties (CPs) (**first row**), the number of genetic variants (or SNPs; **second row**), the number of covariates **(third row)**, and the number of individuals (or samples) per party (**fourth row**). For all experiments, we evenly split the lung cancer dataset between two CPs and replicated as needed to obtain a dataset of desired dimensions. We modified the GWAS parameters to ensure that the amount of data at each step grew proportionally with the input dimensions. For communication, we separately measured the amount of data sent by the auxiliary party in MPC (CP-0); by the 'hub' party that aggregates/broadcasts intermediate results (CP-Hub); and by each of the other CPs (CP-Other). The total communication (in black) accounts for data transfer among all parties. Both runtime and communication scale linearly with each parameter. Due to the vector-wise encryption (in groups of 8,192 in our setting), the computational cost increases in steps as the dataset grows (e.g., see the identical costs of 9,178 and 13,767 samples per party).

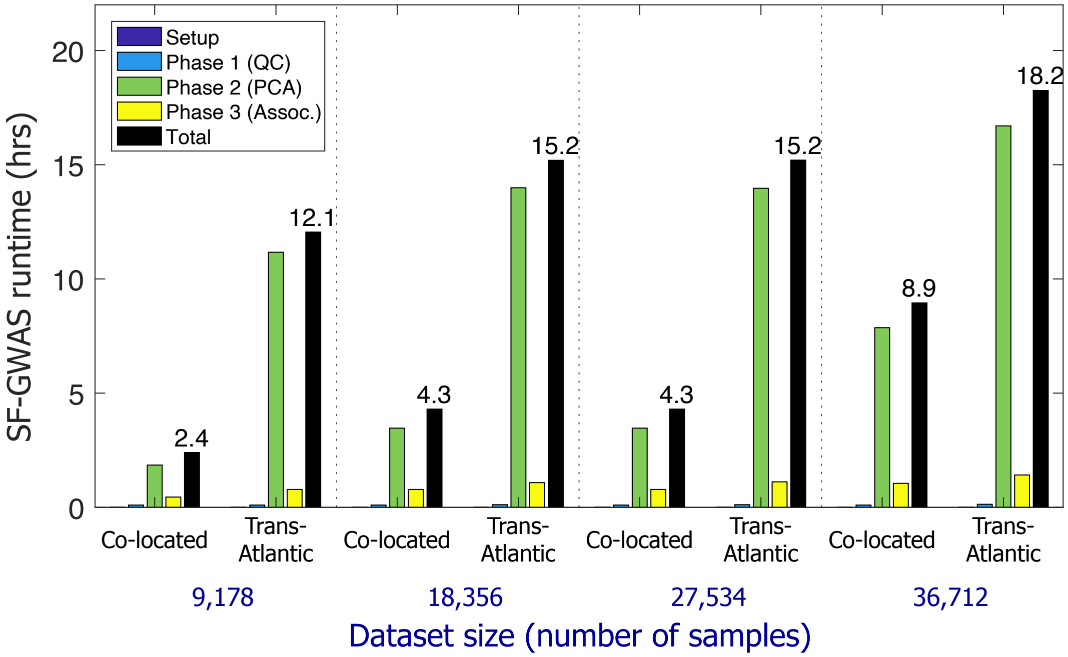


Supplementary Figure 6: **SF-GWAS remains practical for a trans-Atlantic wide-area network setting between the US and the UK.** We evaluated the applicability of SF-GWAS for international collaborations by running the PCA-based workflow on the lung cancer dataset with three computing parties (CPs) located in different geographic regions: two main CPs (each with a half of the data) are placed in Oregon (US; us-central1 in Google Cloud Platform) and London (UK; europe-west2), and the auxiliary party ($CP_0$) is placed in North Virginia (US; us-east4). This wide-area network setting (Trans-Atlantic) is compared to the original setting (Co-located) with all CPs in North Virginia (US; us-east4). We also replicated the dataset to evaluate the runtime for larger dataset sizes. We observe that SF-GWAS runtime increases by at most 5x when executed among distant machines, and this gap decreases as the dataset gets larger given the increased burden of local computation relative to the communication costs. For a large dataset with 36,712 samples, the trans-Atlantic setting incurs only a 2x slow down. We note that the observed differences are much smaller than the 475-fold increase in the underlying round-trip communication delay in the trans-Atlantic setting; the round-trip latency is 0.2ms between co-located CPs (us-east4), 25ms between different regions within the US (us-central1 and us-east4), and 95ms between the US (us-central1) and the UK (europe-west2).

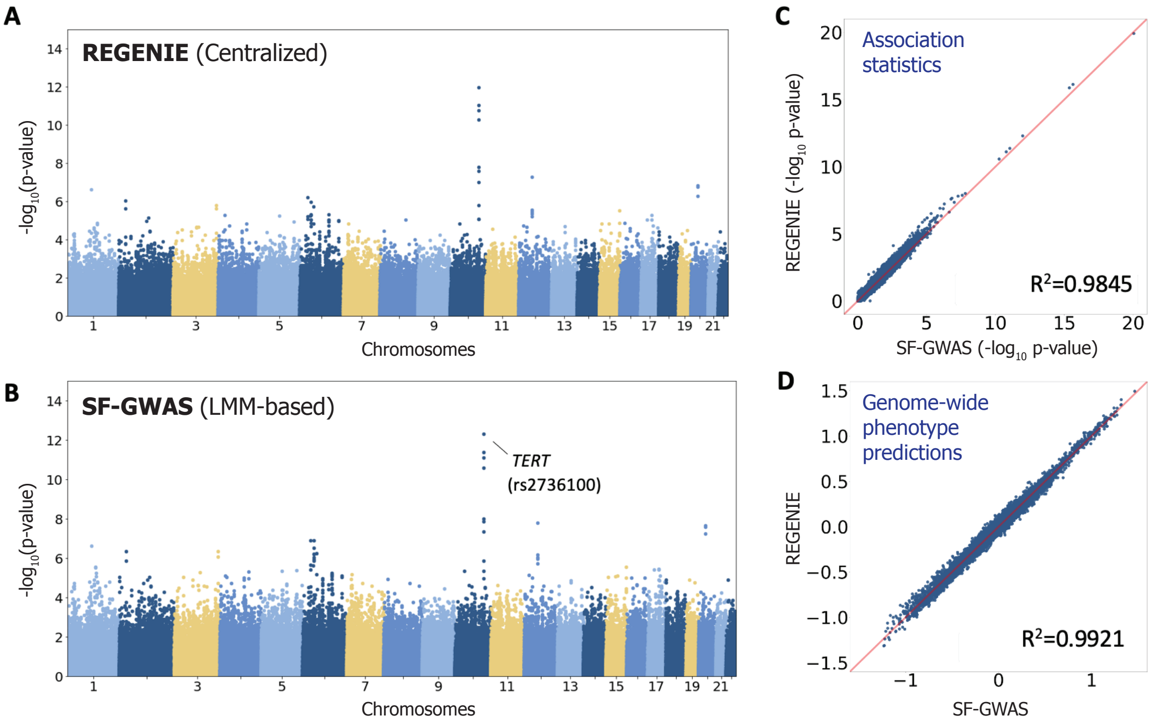


Supplementary Figure 7: **SF-GWAS closely reproduces REGENIE's LMM-based association test results on the lung cancer dataset.** We evaluated LMM-based SF-GWAS on the lung cancer GWAS dataset (including 9,098 individuals and 378,482 SNPs after quality control filters) split between two parties. The Manhattan plot for SF-GWAS (**B**) mirrors the Manhattan plot obtained from running REGENIE on the same centralized dataset (**A**). A particularly strong association is identified for SNP rs2736100, which is associated with the *TERT* gene, a known cancer gene involved in telomere maintenance. (**C**) Comparison of the negative log *p*-values for all variants in the dataset generated by REGENIE and by SF-GWAS. (**D**) Plot of the genome-wide phenotype prediction vectors obtained by both methods, which capture a portion of phenotypic variation that can be explained by genetic relatedness among individuals. These values are provided as input to the association testing pipeline in the REGENIE algorithm to correct for confounding effects. The results from SF-GWAS closely match those of REGENIE with an $R^2$ value over 0.98 in both plots.

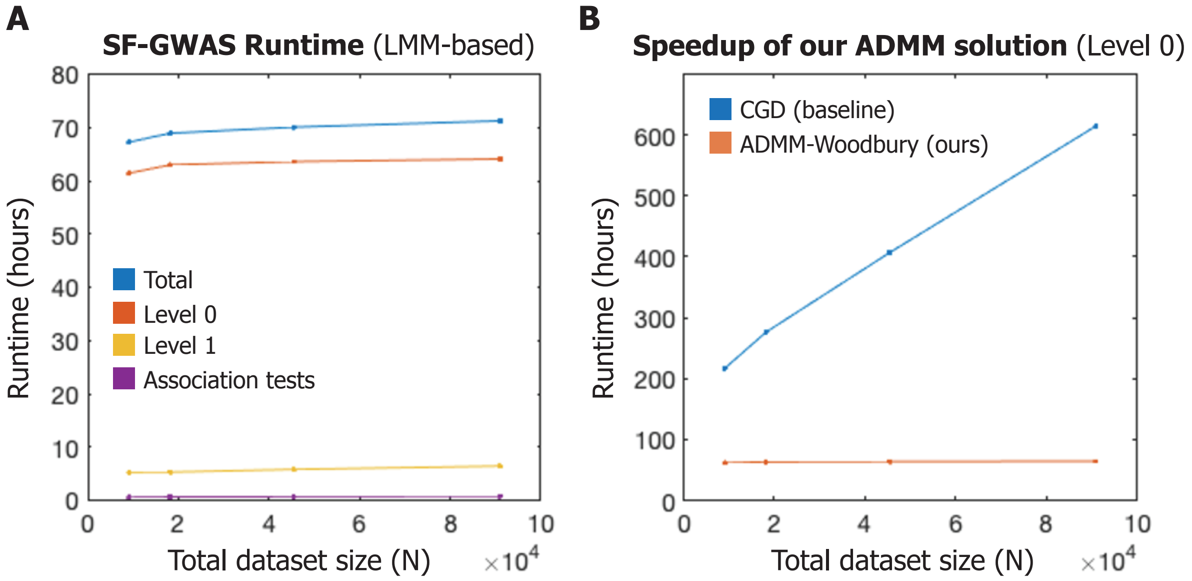


Supplementary Figure 8: **LMM-based SF-GWAS scales efficiently to large datasets with near-constant runtime over a range of dataset sizes.** (**A**) We show the runtimes of LMM-based SF-GWAS on upsampled lung cancer datasets including up to 91K individuals (ten times the original dataset). Measurements are based on a network of three co-located machines on Google Cloud with 12 cores each for parallelization. Runtimes for iterative components of the algorithm with identical computational load for every iteration are estimated based on a smaller set of iterations. SF-GWAS runtime remains near-constant as the size of the dataset grows due to our scalable design of the federated algorithm. (**B**) We compare our optimized ADMM-Woodbury algorithm for Level 0 of the REGENIE workflow (which is the main computational bottleneck) with the baseline conjugate gradient descent (CGD) algorithm, both implemented in our secure and federated framework. The comparison shows the improved asymptotic complexity of our approach (Supplementary Note 6). For the largest dataset including 91K individuals, our approach achieves a 9.6-fold speedup over the baseline.

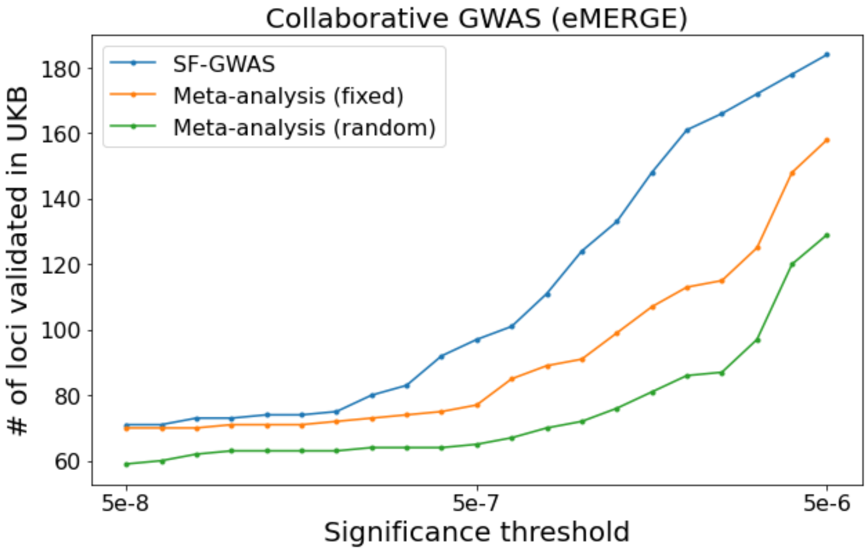


Supplementary Figure 9: **Collaborative GWAS of eMERGE data using SF-GWAS leads to a greater number of validated associations than meta-analysis approaches**. We compare SF-GWAS to both fixed-effects and random-effects models of meta-analysis implemented in the PLINK software for a GWAS of body mass index on the eMERGE data (split across 7 data collection centers) with respect to each method's ability to identify validated associations in the larger UK Biobank cohort. We counted an association as validated if the same locus received a significant $p$-value ($< 5 \times 10^{-8}$) in the summary statistics reported by the Pan-UK Biobank project resource in any ancestry-specific or aggregate analysis of the same phenotype. Plot shows the comparison of the number of validated loci for each method based on varying significance thresholds for the eMERGE analysis from $5 \times 10^{-8}$ to $5 \times 10^{-6}$. Overall, SF-GWAS identified a greater number of validated associations, indicating an increase in statistical power.

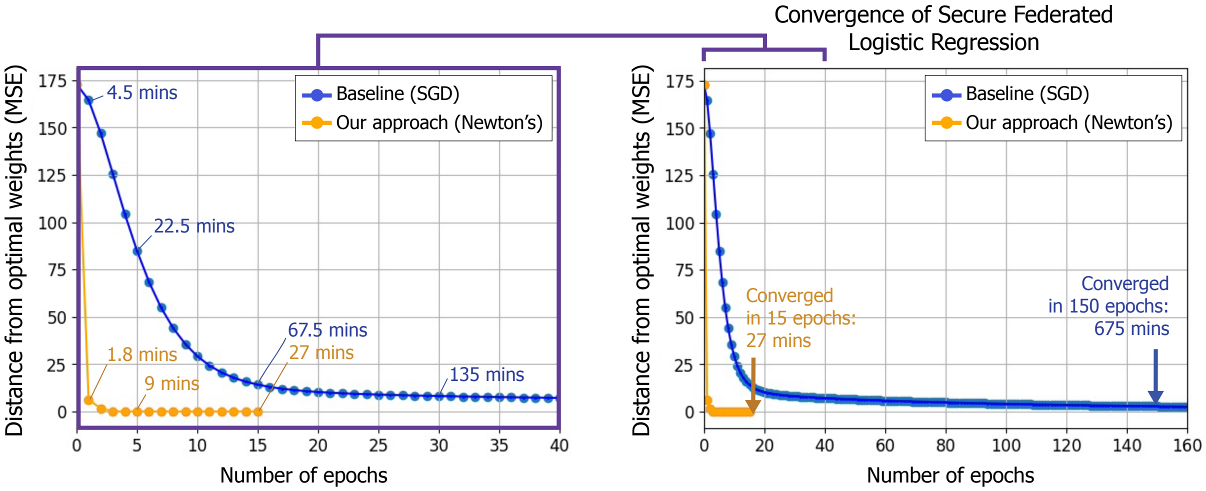


Supplementary Figure 10: **Faster convergence of our secure federated Newton's method for logistic regression compared to standard first-order gradient descent methods.** To illustrate the effectiveness of our Newton's method-based solution for secure logistic regression, we compare the convergence of our secure federated logistic regression pipeline to that of standard mini-batch stochastic gradient descent (SGD)-based optimization of logistic regression models proposed by previous works (e.g., S-GWAS). We evaluate both implementations on the S-GWAS lung cancer dataset, which includes 9,178 samples and 15 covariates. The dataset is split between 2 parties, with each party using a 16-core machine to perform the computation, analogous to our default experimental setup. The ground truth weights used to evaluate convergence are obtained by fitting the model on a centralized, non-encrypted dataset using standard Python libraries. We note that a range of alternative solvers provided by the *statsmodels* and *scikit-learn* libraries all converged to the same final weights. We implemented a secure and federated version of SGD with momentum and regularization to compare against our secure and federated approach for Newton's Method. For SGD, we empirically found that a learning rate $\alpha = 0.9$, a momentum parameter $\mu = 0.99$, a batch size $b = 200$ and a regularization parameter $\rho = 0.0$ were the most effective choices that ensure model convergence in fewer than 200 epochs across all our datasets. In contrast, our Newton's method-based algorithm consistently converged to the optimum within 15 epochs on all datasets. Moreover, our method achieves faster performance in each epoch than SGD (e.g., 1.8 vs 4.5 minutes on the lung cancer dataset). This improved efficiency can be attributed to the greater number of iterations required by SGD in each epoch, which are more costly to perform in the secure federated setting due to the overhead of cryptographic operations. MSE: mean squared error.

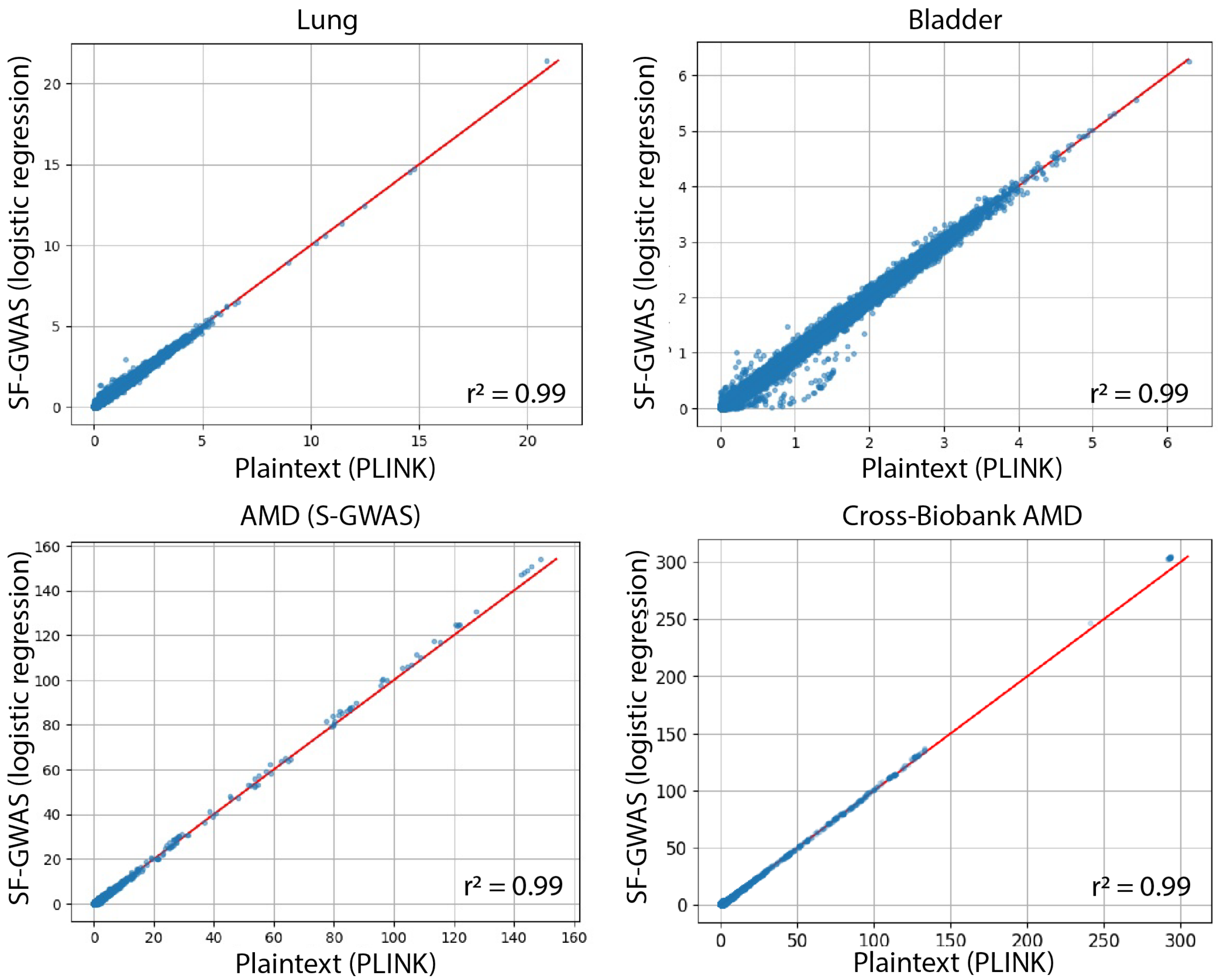


Supplementary Figure 11: **SF-GWAS accurately reproduces end-to-end PCA-based GWAS (logistic) without data centralization.** We reanalyzed the three case-control GWAS datasets presented in Supplementary Fig. 2 from the S-GWAS publication (lung cancer, bladder cancer, and AMD) using logistic regression-based association tests provided by SF-GWAS. The results are compared with those of PLINK's logistic regression-based GWAS pipeline. The same comparison is shown (bottom-right) for our cross-biobank AMD GWAS experiment, where we jointly analyzed three independently collected AMD GWAS datasets from IAMDGC, eMERGE, and UK Biobank (see Methods). SF-GWAS obtains logistic regression-based association statistics that closely match a centralized study in which all plaintext (unencrypted) data are pooled and analyzed together using a standard tool (PLINK).

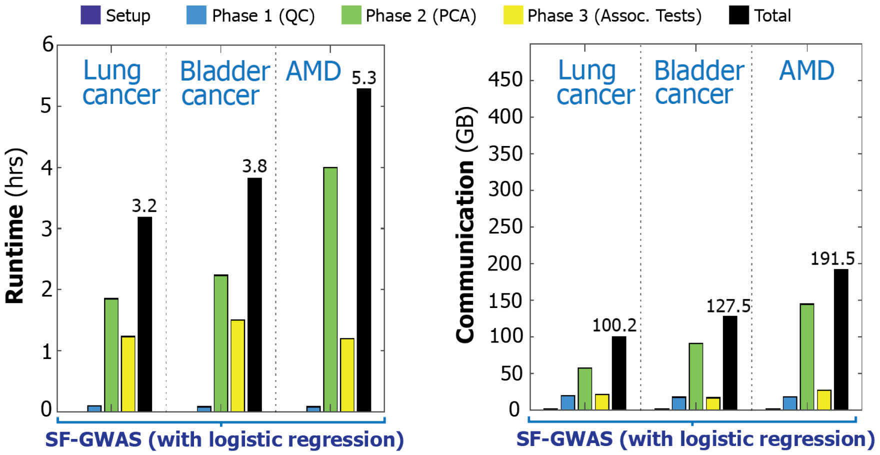


Supplementary Figure 12: **The runtime and communication costs of logistic regression-based SF-GWAS on the three S-GWAS datasets.** We repeat the performance evaluation presented in Figure 2 for the logistic regression-based SF-GWAS. Both runtime and communication costs remain comparable to the linear regression-based workflow despite the greater complexity of the logistic regression models due to our efficient algorithm design.

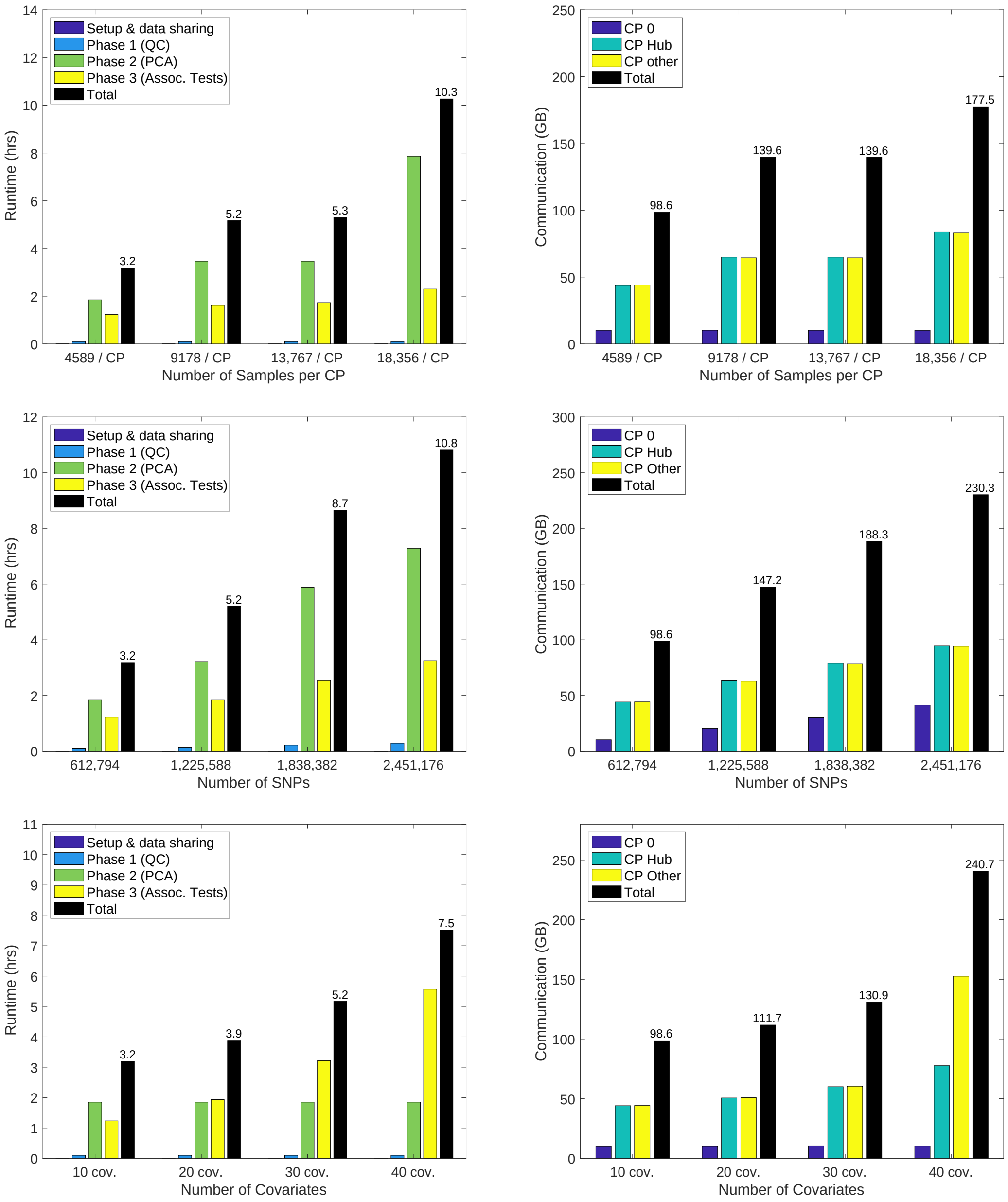


Supplementary Figure 13: **Efficient scaling of logistic regression-based SF-GWAS.** We illustrate the scaling of SF-GWAS runtime **(left column)** and communication bandwidth **(right column)** for the logistic, PCA-based workflow with respect to the number of samples **(top row)**, the number of SNPs **(middle row)**, and the number of covariates **(bottom row)**. We observe similar performance and (linear) scaling behavior compared to the linear regression-based workflow of SF-GWAS, shown in Supplementary Fig. 5. We refer to Supplementary Fig. 5 for evaluation of scaling with respect to the number of computing parties, which remains analogous in this setting.

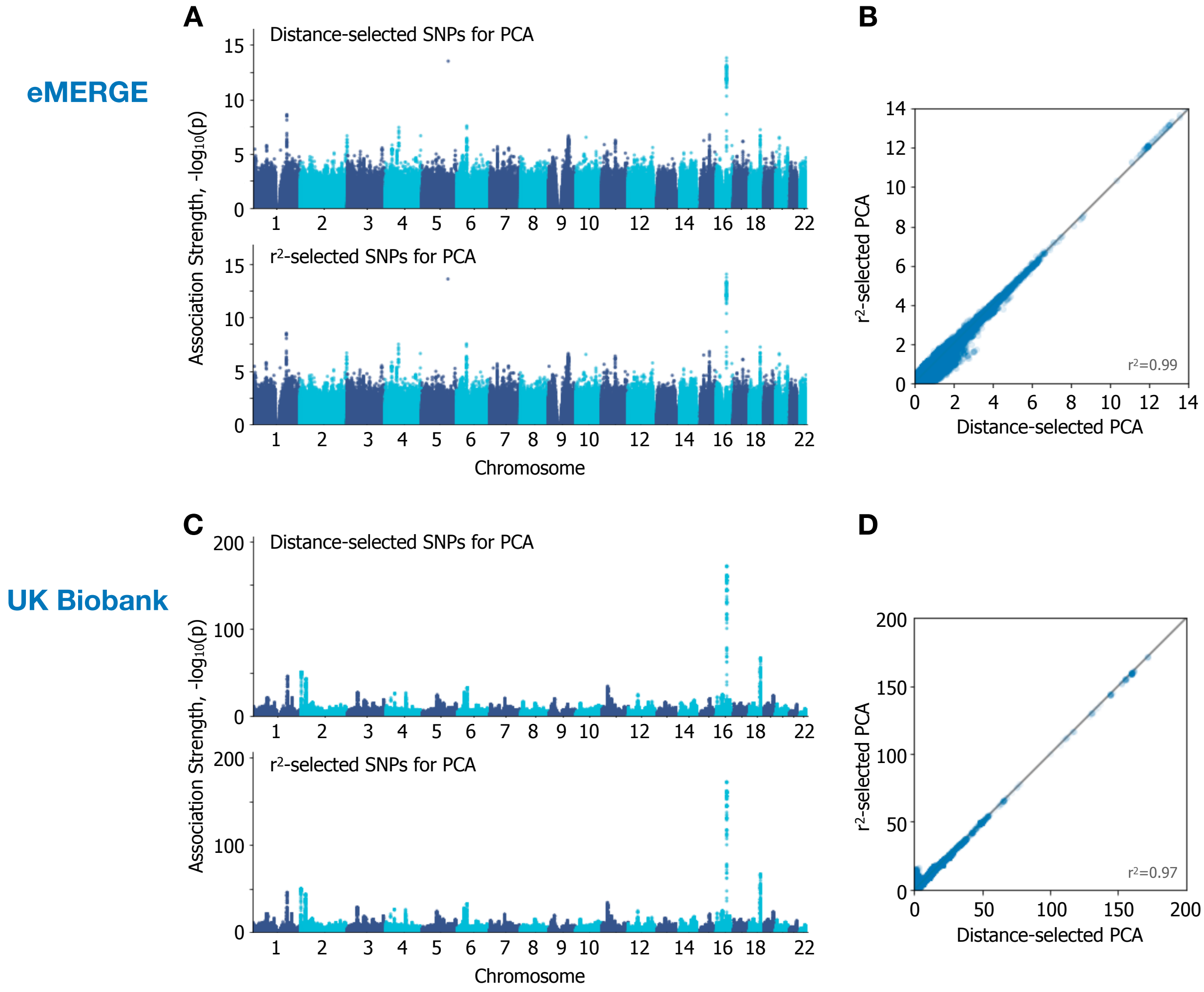


Supplementary Figure 14: **Alternative strategies for selecting a reduced set of SNPs for PCA yield similar GWAS results**. We compared the following two SNP selection strategies: (i) prune the SNPs with pairwise $r^2$ greather than 0.1 using windows of 100 kbp with a default step size of 1 ($r^2$ selection; performed using PLINK); and (ii) Impose a minimum distance threshold of 100 kbp to obtain a pruned SNP set (distance selection). We applied both approaches to the eMERGE (**top**) and UKB (**bottom**) datasets while keeping the rest of the study setting the same, and compared the final GWAS results. We show both the Manhattan plots for individual methods (**A**,**C**) and the scatter plots comparing the association statistics between the two strategies (**B**,**D**). Squared Pearson correlation coefficients ($r^2$) quantifying the agreement in association statistics are reported. Transparency is added to visualize density in the scatter plots. Overall, the difference between the two strategies have only a small impact on the GWAS results, indicating that the global population-level covariates can be reliably constructed based on different genome-wide sets of SNPs.

| Study Group | UK Biobank Assessment Centers | Total Individual Count |
|---|---|---|
| Scotland | Edinburgh<br>Glasgow | 20,970 |
| Northeast England | Middlesborough<br>Newcastle | 31,263 |
| Northwest England | Liverpool<br>Bury<br>Stockport<br>Manchester<br>Leeds | 67,494 |
| Central England | Birmingham<br>Stoke<br>Sheffield<br>Nottingham<br>Wrexham | 61,158 |
| Southeast England | Barts<br>Hounslow<br>Croydon<br>Reading<br>Oxford | 58,351 |
| Wales | Swansea<br>Cardiff<br>Bristol | 36,576 |

Supplementary Table 1: **List of UK Biobank assessment centers grouped by geographic region.** To simulate a federated study on the UK Biobank dataset, we grouped the data collection centers into six study groups based on their geographic location within the UK.

### Supplementary Note 1: MPC Definitions and Operations

The MPC component of SF-GWAS's hybrid secure computation framework generally follows that of S-GWAS [1] with several notable improvements, which we describe later in this section. We first provide key notations and definitions to facilitate a high-level understanding of our framework. We include the implementations of all of our protocols in our open-source software release.

**Setup.** We consider $k$ collaborating entities interacting to perform computation on private data that is not known to any entity other than the data holder. For simplicity, we assume that each pair of parties have a secure and authenticated channel to exchange data. In practice, we adopt a hub-and-spoke network to streamline the communication in aggregation operations. In addition to the $k$ parties, our MPC framework involves an auxiliary coordinating party that is involved only in preprocessing a sequence of random values (generalized Beaver tuples; see [1] for details) in order to distribute correlated randomness to the main parties. These precomputed data facilitate the MPC operations in an efficient manner at the cost of a relaxation in security guarantees (Supplementary Note 7). This setup is commonly known as the trusted-dealer model for preprocessing in MPC [2]. In the following, we refer to the main parties as $\mathsf{CP}_1, \ldots, \mathsf{CP}_k$ (for computing party) and the auxiliary party as $\mathsf{CP}_0$.

**Secret sharing.** We adopt an additive secret sharing scheme based on a finite field $\mathbb{F}_q$ with a base prime $q$ (i.e., a set of integers modulo $q$, equipped with modular addition and multiplication for arithmetic operations). Each real data value $x \in \mathbb{R}$, is encoded as an element in $\mathbb{F}_q$ by scaling it with a factor $2^f$ then mapping it to the closest integer. Depending on the usage, $f$ may be set to different values (e.g., $f = 0$ for encoding integers directly in the field). Also note that the sign of $x$ is represented in $\mathbb{F}_q$ similar to how signed integers are represented in machines: positive values are mapped to $[0, \frac{q-1}{2})$ and negative values are mapped to $[\frac{q-1}{2}, q-1]$ (with larger negative values being farther away from zero).

Given this encoding scheme, we can represent a secret sharing of $x \in \mathbb{F}_q$ as a tuple

$$[x] := ([x]_1, \ldots, [x]_k),$$

where $k$ is the number of collaborating entities and $\sum_{i=1}^{k} [x]_i = x$ (modular addition in $\mathbb{F}_q$). The tuple represents the view of each party: the $i$-th coordinate $[x]_i$ is held by $\mathsf{CP}_i$ only and unknown to the rest. Initial construction of these shares for private data $x$ held by $\mathsf{CP}_1$ (without loss of generality) is performed by $\mathsf{CP}_2, \ldots, \mathsf{CP}_k$ setting their shares $[x]_j = r_j \in \mathbb{F}_q$ to uniformly random elements, and $\mathsf{CP}_1$ setting $[x]_1 = x - \sum_{j=2}^{k} r_j$. This ensures the sum property while hiding $x$ from anyone other than the data holder. Note that the random elements $r_j$ need not be shared over the network; they can be sampled from shared randomness between $\mathsf{CP}_1$ and each of the other parties. This optimization is discussed in S-GWAS [1]. In Supplementary Note 3, we present another way to construct these shares starting from a HE ciphertext. Additionally, we denote by $[\boldsymbol{x}]$ the secret sharing of a vector of values $\boldsymbol{x}$.

**MPC operations.** Once the input data is shared among the parties as secret shares, one can use a range of MPC routines to perform computation over the underlying private values without leaking information about them during the process. These routines are generally of the form:

$$[y] \leftarrow \mathsf{MPC\text{-}Protocol}([x_1], \ldots, [x_s]),$$

where, at the beginning of the protocol, $s$ input values are available to the parties as secret shares and the output consists of a new private value $y$ that is secret shared among the parties (thus the underlying value of $y$ is not revealed). Note that this property allows these routines to be *composed*—i.e., the output $[y]$ can in turn be considered as input to another MPC protocol to carry out further computation on $y$.

Our MPC framework includes protocols for a range of basic operations, including the following: addition, multiplication, truncation (of a specified number of least significant bits), division, square root, square root inverse, and comparison (see S-GWAS [1] for protocol definitions). These operations enable a wide range of complex MPC protocols to be built, including the eigendecomposition protocol, which SF-GWAS performs during PCA using secret shares.

**Novel improvements.** We made several key improvements upon the prior MPC framework of S-GWAS [1]. First, SF-GWAS software is based on a new implementation of the entire MPC codebase in Go, featuring a number of software engineering improvements such as representing a vector of finite field elements as a compact array of native integer types (as opposed to a separate heap object for each element represented as a big integer in S-GWAS), and better parallelization leveraging the built-in concurrency features of Go (i.e., goroutines). Second, we have improved the protocols for comparison, square root, square root inverse, and division to leverage efficient bit decomposition-based routines, which first convert the input secret shares in $\mathbb{F}_q$ to bit-wise shares in $\mathbb{F}_2$, then use a sequence of bit operations to compute the desired function. We followed the roadmap for a comparison routine introduced in [3] (see ReLU protocol) and newly extended this approach to accelerate the square root and division protocols of S-GWAS. Third, enabled by the improved efficiency of our implementations, we changed the default security parameters in our framework to provide 128 bits of statistical security, substantially higher than 30 bits provided by S-GWAS. All of our comparisons are based on the 128-bit setting of SF-GWAS and the original 30-bit setting of S-GWAS. Finally, we newly make joint use of both MPC and MHE routines to avoid secret sharing the large input datasets, as described in later sections.

### Supplementary Note 2: MHE Definitions and Operations

In addition to the MPC framework, SF-GWAS utilizes the multiparty fully-homomorphic encryption (MHE) scheme proposed by Mouchet et al. [4], which we instantiate with the Cheon-Kim-Kim-Song (CKKS) cryptosystem [5] as implemented in the Lattigo library [6]. In the following, we introduce key notations and the main MHE operations used in SF-GWAS. We also explain how we exploit key properties of MHE to efficiently compute on encrypted data. As with our MPC framework, the implementations of all of our MHE routines are included in our software release.

**Setup.** As before, we consider $k$ collaborating entities $\mathsf{CP}_1, \dots, \mathsf{CP}_k$ with secure and authenticated channels connecting them in a hub-and-spoke network. Each party $\mathsf{CP}_i$ has a share $sk_i$ of the secret key $sk$ (based on an additive secret sharing scheme) and they all collaborate to generate the corresponding collective public key $pk$. As a result, each $\mathsf{CP}$ can independently compute on ciphertexts encrypted under $pk$, but all parties have to collaborate to decrypt a ciphertext. This ensures a collective protection of the encrypted data and corresponds to a $k$-out-of-$k$ threshold encryption scheme [7].

**Homomorphic encryption (HE) with CKKS.** CKKS is a homomorphic encryption scheme that enables approximate arithmetic over $\mathbb{C}^{\mathcal{N}/2}$ and in which the plaintext and ciphertext spaces share the same domain $R_{Q_L} = \mathbb{Z}_{Q_L}[X]/(X^{\mathcal{N}}+1)$, with $Q_L = \prod_0^L q_i$ in our case and $\mathcal{N}$ a power of 2. The plaintexts/ciphertexts are represented by polynomials of degree up to $\mathcal{N}-1$ (with $\mathcal{N}$ coefficients in $\mathbb{Z}_{Q_L}$) in this domain and they encode/encrypt a vector of up to $\mathcal{N}/2$ floating-point values. Any operation on a plaintext/ciphertext is simultaneously performed on all encoded/encrypted values, i.e., in a *Single Instruction, Multiple Data* (SIMD) manner.

The CKKS parameters are denoted by the tuple $(\mathcal{N}, \Delta, \eta, mc)$. $\mathcal{N}$ is the ring dimension; $\Delta$ is the plaintext scale by which any value is multiplied before it is quantized and encrypted/encoded; $\eta$ is the standard deviation of the noise distribution; and $mc$ represents a chain of moduli $\{q_0, \dots, q_L\}$ such that $\Pi_{\iota \in \{0,\dots,\kappa\}} q_\iota = Q_\kappa$ is the ciphertext modulus at level $\kappa$, with $Q_L$ the modulus of fresh ciphertexts. Operations on a ciphertext $\boldsymbol{c}$ at level $\kappa$ and scale $\Delta$ with $\Delta < Q_\kappa$ are performed modulo $Q_\kappa$. We denote by $\{\boldsymbol{c}, L, \Delta\}$, with $\boldsymbol{c} = (\boldsymbol{c}_0, \boldsymbol{c}_1) \in R^2_{Q_L}$, a fresh ciphertext at level $L$ with scale $\Delta$ and by $\boldsymbol{x} \in R_{Q_L}$ a plaintext. In the remainder of this paper, we refer directly to an encrypted (i.e., under the public key $pk$) vector as $\boldsymbol{c}_{pk}$ and to a plaintext vector as $\boldsymbol{x}$. We refer to the work of Cheon et al. [5] for further details on the CKKS cryptoscheme. In our work, we set the default parameters to $\mathcal{N} = 2^{14}$, $\Delta = 2^{34}$, $\eta = 3.2$, and a modulus chain with $L = 9$ and $\lfloor \log_2 Q_L \rfloor = 351$ for a 128-bit security level, based on a standard configuration provided in the Lattigo library [6]. Our software allows the user to choose a different set of cryptographic parameters if desired.

An important aspect of CKKS is its addition of ciphertext noise (required for security of the system) in the least significant bits of the encrypted values. This leads to an approximate computation scheme and thus necessitates a thoughtful management of noise levels throughout the computation. In SF-GWAS, we rely on general scale management techniques [8] to control the noise in CKKS, which are implemented in the underlying cryptographic library (Lattigo [6]).

In CKKS, one can perform element-wise additions and multiplications, as well as cyclic rotations, on the vectors encoded in ciphertexts or plaintexts. Cyclic rotations shift the vector elements in a ciphertext by a desired number of positions (with wraparound). When combined with other operations, rotations enable useful vector-based operations, such as element selection (e.g., by multiplying the ciphertext with a plaintext encoding a one-hot vector to single out an element, then rotating to place it in a desired position) and inner products (e.g., by multiplying two ciphertexts, then iteratively rotating and adding to accumulate all pairwise products). A multiplication between a plaintext and a ciphertext is more efficient than a multiplication between two ciphertexts (e.g., seven times faster in the SF-GWAS setting). The complexity of a rotation is equivalent to multiplying two ciphertexts. In addition, after a certain number of multiplications on a ciphertext, the ciphertext needs to be 'refreshed' before it can be used for further multiplications—an operation known as *bootstrapping.* Bootstrapping is computationally expensive and usually at least an order of magnitude slower than a multiplication [9]. Finally, the ciphertext $\boldsymbol{c} = (\boldsymbol{c}_0, \boldsymbol{c}_1)$ is decrypted by computing $\boldsymbol{c}_0 + sk \cdot \boldsymbol{c}_1$, where $sk$ is the secret decryption key.

**Multiparty homomorphic encryption (MHE) with CKKS.** In the multiparty instantiation of CKKS, the $\mathsf{CP}$s rely on public evaluation keys, e.g., the collective encryption key $pk$, which are cooperatively generated during the setup. The $\mathsf{CP}$s can then independently perform the native CKKS operations, i.e., additions, multiplications and rotations, on collectively encrypted ciphertexts. To decrypt a ciphertext in MHE, the $\mathsf{CP}$s execute an interactive protocol in which they each partially decrypt the ciphertext using their secret key $sk_i$, then aggregate to reveal the decryption result (i.e., computing $\boldsymbol{c}_0 + (\sum_{i=1}^{k} sk_i) \cdot \boldsymbol{c}_1$). In addition, in the multiparty scheme, we have access to a lightweight, interactive protocol for bootstrapping, which can be used instead of the expensive centralized bootstrapping. In this case, the $\mathsf{CP}$s cooperatively perform a blinded decryption and re-encryption of the input ciphertext (see [4] for protocol details).

**Novel improvements.** Building upon prior applications of MHE [10], we introduce the following extensions to the framework in this work. First, we develop a hybrid framework combining MHE with secret sharing-based MPC routines by switching between the two schemes throughout the SF-GWAS algorithm (Supplementary Note 3). Using this new approach, we improve both the runtime and accuracy of nonlinear operations, such as comparison, division, and square root. These operations are costly to perform accurately using existing HE techniques, which require a high-degree polynomial approximation (involving many ciphertexts multiplications) in order to handle inputs with a wide range of possible values. In contrast, there are efficient and accurate MPC routines for the same operations leveraging bit-wise manipulation. Note that our GWAS pipelines, including the PCA and LMM algorithms, frequently invokes these routines, thus greatly benefiting from the hybrid approach.

Next, we implement a range of different matrix multiplication routines in MHE to optimize the performance of SF-GWAS, which heavily relies on matrix multiplications involving the large genotype matrix. We optimized these routines specifically for the federated setting, where the local dataset is available in plaintext and can be used to lower the computational cost. The different strategies we implemented include: (i) inner product calculations for each row-column pair from the two input matrices; (ii) an element-wise approach, where each element in the left matrix is singled out, replicated, then multiplied with the corresponding row in the right matrix to be added to the result; and (iii) optimized block matrix multiplication routine from [11] (which we extend to large matrices with multiple blocks). We choose a different strategy depending on the expected sizes of input matrices in each step of the algorithm to maximize the overall performance.

Finally, our practical application of MHE to end-to-end GWAS is enabled by a number of algorithmic design techniques aimed at taking advantage of the federated computational setting brought by MHE. The efficient MHE implementations we developed for major steps of SF-GWAS involving distributed linear algebra, e.g., QR decomposition, PCA, logistic regression, ridge regression, and solving a linear system of equations, are of independent interest. We describe our algorithm design in more detail in Supplementary Notes 4–6.

### Supplementary Note 3: MHE-MPC Conversion

SF-GWAS switches between the MHE and secret-sharing representations of the private data (encompassing intermediate computation results), which allows efficient state-of-the-art MPC routines to be used in conjunction with MHE operations to carry out the global computations in an optimized manner. For example, we perform large-scale matrix and vector operations using MHE to exploit its SIMD property, while evaluating non-polynomial functions (division, square root, and comparison) using MPC protocols, which are more efficient and numerically stable than the existing MHE counterparts. Any operation involving the local unencrypted data is performed using HE to avoid secret sharing the large input datasets.

To switch from MHE to MPC, a globally shared ciphertext in MHE can be transformed to secret shares through a collective, masked (or blinded) decryption by the $\mathsf{CP}$s as follows. Let $\mathsf{CP}_1$ be the hub party of the network. Each of the other parties $\mathsf{CP}_i$ with $i = 2, \ldots, k$ partially decrypts the ciphertext locally and adds a random number to each coordinate of the vector to mask the result. The same random numbers are then encoded in the secret sharing domain and subtracted from zero to obtain the secret shares of the corresponding party. The hub party, on the other hand, aggregates all partial decryption results from other parties (with the masks added) and adds to its own partial decryption result to obtain the masked private numbers in the plaintext domain. These are then re-encoded in the secret sharing domain to obtain the secret shares of the hub party. Correctness follows from the fact that the addition of all shares cancel out the masks applied by the individual parties to reveal the private values corresponding to the real decryption results of the input ciphertext. This procedure is formally described in Protocol 1.

Similarly, to switch from MPC to MHE, the $\mathsf{CP}$s first reveal the blinded input by masking the shares with random numbers and aggregating them. Then, each party encrypts its own masks in the CKKS domain before all parties aggregate the encrypted masks. By locally subtracting the aggregated mask from the encryption of the blinded input, the $\mathsf{CP}$s obtain the MHE ciphertext encoding the private numbers represented by the input secret shares. See Protocol 2 for details.

Both protocols rely on the statistical security of revealing blinded private numbers in the plaintext space, which are in turn re-encoded in the target scheme. SF-GWAS is based on system parameters that ensure 128-bits of statistical security in these operations. This is achieved by using random masks whose bit-length exceeds the input data range by 128 + 30 bits, accounting for $2^{30}$ instances of this task as a general upper bound. Note that applying a union bound over $j$ instances of a leakage increases the adversary's advantage in distinguishing the observed masked value from a uniformly random number by a factor of $j$, hence a multiplicative factor of $2^{30}$ is used as a buffer for this compounding effect.

---

**Protocol 1** Conversion from MHE to MPC

---

**Input:** All parties have the same ciphertext (e.g., via broadcasting) $\boldsymbol{c}_{pk} = (\boldsymbol{c}_0, \boldsymbol{c}_1) \in R^2_{Q_\tau}$ , an encryption of the plaintext $\boldsymbol{x}$ (i.e., $\boldsymbol{x} \approx \boldsymbol{c}_0 + (\sum_{i=1}^{k} sk_i) \cdot \boldsymbol{c}_1$); $\lambda$ is a security parameter, $sk_i$ each $\mathsf{CP}_i$'s share of the secret key, $\chi_{err}$ a noise distribution over $R$, where each coefficient is independently sampled from a centered Gaussian distribution with standard deviation $\sigma = 3.2$, and bound $\lfloor 6\sigma \rfloor$. We denote by Encode$(\cdot)$ the mapping from a plaintext encoded in $R$ to the equivalent encoding in $\mathbb{F}_q$. Let $T$ be the bound on all possible coefficients in the polynomial representation of $\boldsymbol{x}$ encoding real data values.
**Output:** Each $\mathsf{CP}_i$ obtains $[\boldsymbol{x}]_i$ such that $\sum_{i=1}^{k} [\boldsymbol{x}]_i \approx \boldsymbol{x}$
**Constraint:** $Q_\tau > (k+1) \cdot T \cdot 2^\lambda$

```
Each CP_i for i = 2, ..., k:
    Samples a_i ← Uniform(R_{T·2^λ}) and e_i ← χ_err
    Computes h_i = sk_i · c_1 + a_i + e_i  mod Q_τ, and sends h_i to CP_1
    Assigns [x]_i = Encode(−a_i)
CP_1:
    Samples a_1 ← Uniform(R_{T·2^λ}) and e_1 ← χ_err
    Computes h_1 = c_0 + sk_1 · c_1 + a_1 + e_1  mod Q_τ
    Receives h_i's from other parties and computes h' = Σ_{i=1}^k h_i // Note: h' ≈ x + Σ_{i=1}^k a_i + Σ_{i=1}^k e_i
    Assigns [x]_1 = Encode(h' − a_1)
```

---

**Protocol 2** Conversion from MPC to MHE

**Input:** Each $\mathsf{CP}_i$ (for $i \in 1, ..., k$) has $[\boldsymbol{x}]_i$ such that $\sum_{i=1}^{k} [\boldsymbol{x}]_i = \boldsymbol{x}$; Let $\lambda$ be the security parameter, and $\ell$ be the bound on the possible bit lengths of real data values $\boldsymbol{x}$ as encoded in $\mathbb{F}_q$; Let the length of $\boldsymbol{x}$ be $L$, equal to the number of plaintext slots in an MHE ciphertext; $\mathrm{Enc}(pk, \cdot)$ denotes MHE encryption with public key $pk$.
**Output:** All parties obtain $\boldsymbol{c}_{pk}$, an encryption of $\boldsymbol{x}$.
**Constraint:** $q > (k+1) \cdot 2^{\lambda+\ell}$

Each $\mathsf{CP}_i$ for $i = 1, \ldots, k$:
  Samples $\boldsymbol{b}_i \leftarrow \mathrm{Uniform}(\mathbb{Z}^{L}_{2^{\lambda+\ell}})$
  Computes $\boldsymbol{r}_i = [\boldsymbol{x}]_i - \boldsymbol{b}_i \mod q$
  Encrypts $\boldsymbol{b}_i$ to obtain $\boldsymbol{c}^{(i)}_{pk} = \mathrm{Enc}(pk, \boldsymbol{b}_i)$
Each $\mathsf{CP}_i$ for $i = 2, \ldots, k$: // Excluding $\mathsf{CP}_1$
  Sends $\boldsymbol{r}_i$ and $\boldsymbol{c}^{(i)}_{pk}$ to $\mathsf{CP}_1$
$\mathsf{CP}_1$:
  Receives $\boldsymbol{r}_i$'s from other parties and computes $\boldsymbol{r}' = \sum_{i=1}^{k} \boldsymbol{r}_i \mod q$ // Note: $\boldsymbol{r}' = \boldsymbol{x} - \sum_{i=1}^{k} \boldsymbol{b}_i$
  Encrypts $\boldsymbol{r}'$ to obtain $\boldsymbol{c}'_{pk} = \mathrm{Enc}(pk, \boldsymbol{r}')$
  Receives $\boldsymbol{c}^{(i)}_{pk}$'s from other parties and computes $\boldsymbol{c}_{pk} = \boldsymbol{c}'_{pk} + \sum_{i=1}^{k} \boldsymbol{c}^{(i)}_{pk}$
  Broadcasts $\boldsymbol{c}_{pk}$ to other parties
Each $\mathsf{CP}_i$ for $i = 2, \ldots, k$:
  Receives $\boldsymbol{c}_{pk}$ from $\mathsf{CP}_1$

The conversion protocols introduce additive noise that scales linearly in the number of parties (as discussed in [4]). Because our application setting involves labs and organizations holding genomic data from a large group of individuals, the number of parties is limited in practice. Note that the addition of ciphertexts leads to an analogous increase in noise, linear in the number of additions. Some of our matrix multiplication routines involve thousands of additions (e.g., for an 8192-by-8192 matrix). Our experimental results demonstrate that these linear increases in noise do not undermine the overall accuracy of our system.

We provide the runtime and communication costs of the conversion protocols in Supplementary Table 2. Since these protocols perform a masked decryption of the input values, followed by a removal of the mask in the target scheme, their communication cost is comparable to that of collective decryption.

### Supplementary Note 4: Algorithm Design Techniques

Here, we describe our main strategies to design an efficient, secure, and federated protocol for end-to-end GWAS. These strategies may be helpful in developing other genomic analysis workflows using our hybrid framework. Recall that we have the flexibility to switch between MHE and secret sharing-based MPC schemes (Supplementary Note 3) to jointly utilize their computational primitives. Another important aspect of our federated setting is that each CP has access to their own unencrypted (plaintext) input data, which can be used to minimize the workload and communication throughout the computation.

**Accurately analyze data by emulating centralized computation without the need for data sharing.** A common workaround for analyzing distributed datasets that cannot be pooled together is to independently analyze each dataset and then to combine the results after the fact (e.g., via meta-analysis). While such an approach can be effective in large-data settings, it severely limits the types of analyses that can be performed and often requires assumptions about data distributions or simplification of the problem setting (e.g., exclusion of global principal components) in order to obtain a consistent definition of the intermediate results to be combined across multiple parties. Our work is instead based on the key observation that, using modern cryptographic tools such as MHE and MPC, a group of collaborators can perform joint computations as if they had access to the pooled data, but without actually sharing any private data among them. SF-GWAS is designed to enable this alternative approach to collaboration for end-to-end GWAS workflows. This is generally achieved by formulating the high-level algorithm for the centralized setting, then implementing the secure and federated version of each step using the available MHE and MPC routines. For example, in quality control, the desired computation in one step is to filter the SNPs based on a minor allele frequency (MAF) threshold. Analogous to the centralized setting, SF-GWAS computes the MAF globally for each SNP then compares to the threshold. SF-GWAS achieves this without sharing the input data by having the parties individually compute local statistics, securely aggregate them, then securely compare to the threshold using the MPC comparison routine. Overall, SF-GWAS closely emulates the centralized GWAS pipeline.

**Redesign the target algorithms to maximize use of low-cost operations and minimize communication by leveraging local plaintext data.** Prior works on secure GWAS based on either HE or MPC [1, 12] require sharing or centralizing the entire input dataset among the collaborating entities, albeit in an encrypted form. For modern large-scale GWAS datasets, this requirement leads to overwhelming communication and computation overheads (Fig. 2). SF-GWAS eliminates this overhead by adopting MHE, which allows the CPs to locally keep the original input datasets and share only smaller amounts of intermediate results via MHE and MPC routines to carry out the global computation. Not only does this vastly reduce communication by not sharing the input datasets, but it also enables SF-GWAS to maximize the use of low-cost operations involving the unencrypted plaintext input throughout the algorithm. Notably, large-scale matrix multiplications required in both PCA and LMM algorithms can now be formulated as ciphertext-plaintext multiplications, for which we implement specialized matrix multiplication routines with improved efficiency. We also restructure some of the computations to maximize the use of efficient cleartext operations. For example, we introduce a new subroutine (see Protocol 5) in PCA-based SF-GWAS to perform the SNP dosages standardization on-the-fly and thus keep the large-scale genotype matrix in plaintext during the entire protocol. In LMM-based SF-GWAS, we reformulate the federated ADMM protocol (see Protocols 11 and 12) using a matrix identity to perform the computationally-intensive matrix inversion on plaintext local data before combining the parties' local results.

**Switch between MPC and MHE routines for efficient secure computation.** The GWAS computation entails large-scale matrix and vector multiplications as well as the evaluation of non-polynomial functions, such as square root, division, and comparison (see Supplementary Notes 5 and 6). While the matrix/vector multiplications can be efficiently performed in MHE leveraging the packing and SIMD properties of the underlying cryptosystem (enabling parallelization), employing MPC routines for smaller-scale, non-polynomial operations is typically more efficient and more accurate than performing them using MHE. Note that, when computing the square root of a single element (e.g., performed once for each active column in the QR factorization algorithm), ciphertext packing is highly underutilized and adds a significant performance overhead. In addition, the evaluation of non-polynomial functions in MHE requires high-depth polynomial approximations for predefined intervals; in sophisticated and multi-step workflows such as GWAS, an appropriate choice of approximation interval is challenging to determine without incurring too much performance penalty by setting a wide range. In contrast, in MPC frameworks, there exist efficient protocols for computing the bit-length of a secret-shared value [13], which can be used to map the input to a common interval for accurate polynomial approximation. With the goal of using the best available routine for each type of operation in the protocol, SF-GWAS employs MHE to execute large-scale matrix operations and performs non-polynomial operations, as well as small-matrix operations, using secret sharing-based MPC routines.

**Vectorized encoding for efficiently composing linear algebra operations on encrypted matrices.** Linear algebra operations on large-scale matrices are frequently invoked in the GWAS workflow. For example, in PCA-based SF-GWAS, QR factorization is repeatedly performed in-between matrix multiplications during the power iteration procedure in PCA (Supplementary Note 5) and requires operations using each column of the matrix as a unit. Retrieving a column from a matrix of $m$ rows, where the rows are individually packed in ciphertexts, would require $m$ homomorphic multiplications, additions, and rotations, whereas it incurs no cost when the matrix is already encoded column-wise. Similarly in the LMM workflow, update equations for our iterative algorithms for ridge regression include a sequence of matrix multiplications whose efficiency depends on the choice of encoding (i.e., row-wise vs. column-wise). In fact, the overwhelming overhead of transforming encrypted matrices from one encoding to another would make SF-GWAS practically infeasible. Our MHE implementations of various components of GWAS are based on a consistent vectorized encoding scheme for encrypted matrices, and we tailor the operations to efficiently work with the chosen encoding without costly conversions.

### Supplementary Note 5: SF-GWAS Algorithm for PCA-based GWAS

Here, we describe the individual steps of the SF-GWAS workflow. We first present the details of the PCA-based GWAS pipeline and address the LMM-based workflow in Supplementary Note 6. We assume that before the protocol is executed, the collaborating entities have agreed upon the protocol parameters. These include: security parameters for MPC and MHE frameworks (e.g., base primes and the scaling factor for encoding continuous values); GWAS parameters (e.g., dataset dimensions, quality control filters, number of iterations in randomized PCA); and network parameters (e.g., IP addresses and port numbers). A complete list of parameters is documented in our software.

**Input data.** SF-GWAS supports two formats for the input data: (i) a custom binary matrix format, where the data is divided into blocks of genomic regions for block-wise processing; or (ii) the PLINK2 PGEN format, which is meant for large-scale datasets and can be conveniently preprocessed using PLINK2. Note that in our experiments, we used (i) for the three datasets from S-GWAS, and (ii) for eMERGE and UKB datasets. During the main protocol, SF-GWAS retrieves the individual-by-SNP genotype matrix from these input files in a streaming fashion to process them on-the-fly, without importing the whole dataset into memory. In addition to the genotype data, SF-GWAS takes in a covariate file (including a matrix where the rows represent individuals and columns represent different covariate features) and a phenotype file (including a vector of phenotypes for all individuals). Note that these files are locally kept unencrypted and each party provides a different set of input files to the SF-GWAS program to be jointly and securely analyzed over the network.

**Initial setup.** Secure TLS channels are established between pairs of parties (can be simplified to a hub-and-spoke network, if desired). For each channel, a key exchange protocol sets up a shared secret between pairs of parties, which is used to set the seed for the shared pseudorandom generators (PRGs). MHE key generation protocols are invoked to interactively generate a collective encryption key, a secret-shared decryption key, a re-linearization key, and a set of rotation keys for pre-determined values of coordinate shifts expected to be encountered throughout the algorithm (i.e., all powers of two up to the total number of slots in a ciphertext and all integers up to a small constant depending on the number of covariates/principal components).

**Protocol notation and subroutines.** For simplicity, all data variables are considered encrypted or blinded, either via MHE or secret sharing based on the context, by default. Cleartext variables are distinguished with an overline, e.g., $\bar{\mathbf{x}}$. Bold lowercase symbols indicate vectors and uppercase indicate matrices. The operators $\times$, $\cdot$, and $\bullet$ indicate matrix multiplication, element-wise product, and inner-product between two vectors, respectively.

Our protocols invoke the following MPC subroutines, which take the input values as secret shares and output the desired result also as secret shares.

- $\mathsf{MPC\text{-}Multiply}(\mathbf{x}, \mathbf{y})$ computes the element-wise fixed-point multiplication of two secret-shared vectors $\mathbf{x}$ and $\mathbf{y}$, involving a secure multiplication followed by a truncation procedure for rescaling. For simplicity, we also use this function to refer to its variants, e.g., when the second input is a public scalar $\bar{y}$.
- $\mathsf{MPC\text{-}Divide}(\mathbf{x}, \mathbf{y})$ computes the element-wise fixed-point division of a secret-shared vector $\mathbf{x}$ by a secret-shared vector $\mathbf{y}$.
- $\mathsf{MPC\text{-}Sqrt}(\mathbf{x})$ computes the square root of each element in a secret-shared vector $\mathbf{x}$.
- $\mathsf{MPC\text{-}SqrtInv}(\mathbf{x})$ computes the inverse square root of each element in a secret-shared vector $\mathbf{x}$.
- $\mathsf{MPC\text{-}Comparison}(\mathbf{x}, \bar{b})$ compares each element in a secret-shared vector $\mathbf{x}$ against a public value $\bar{b}$ and returns a Boolean vector indicating whether each element is greater than $\bar{b}$.

- $\mathsf{MPC\text{-}IsPositive}(\mathbf{x})$ tests the sign of each element of a secret-shared vector $\mathbf{x}$ and returns the result as a binary vector where ones correspond to non-negative values.
- $\mathsf{MPC\text{-}EigenDecomp}(\mathbf{X})$ computes the eigendecomposition of a secret-shared symmetric matrix $\mathbf{X}$ and returns the eigenvectors and associated eigenvalues.
- $\mathsf{MPC\text{-}Aggregate}(\bar{\mathbf{x}}_p)$ aggregates all parties' cleartext vectors implicitly by setting the local shares to the elements of the local vector, which represents secret sharing of the global sum $\sum_p \bar{\mathbf{x}}_p$.
- $\mathsf{MPC\text{-}MatrixInvSqrt}(\mathbf{X})$ computes $\mathbf{B}$ such that $\mathbf{B}^T\mathbf{B} = \mathbf{X}^{(-1)}$ using the eigendecomposition protocol $\mathsf{MPC\text{-}EigenDecomp}$, i.e., by scaling the eigenvectors by the inverse square root of the corresponding eigenvalues.

We refer to the S-GWAS publication [1] for details of the algorithms underlying these basic protocols. Our framework features improved implementations of the core non-polynomial operations Divide, Sqrt, SqrtInv, Comparison, and IsPositive, where we employ bit decomposition to convert the input secret shares into Boolean shares, which are compacted into integer values for efficient parallel computation. We then use optimized Boolean sharing-based bit-length and sign-test protocols (the latter based on the ReLU protocol of [3]) to obtain improved performance of these core routines over the prior framework of S-GWAS.

We also make use of the following standard MHE subroutines on ciphertext data.

- $\mathsf{MHE\text{-}Aggregate}(\mathbf{X}_p)$ aggregates all parties' encrypted $\mathbf{X}_p$ and shares the encrypted result, i.e., $\sum_p \mathbf{X}_p$, with all parties.
- $\mathsf{MHE\text{-}Duplicate}_m(\mathbf{x})$ replicates the first element of encrypted $\mathbf{x}$ to fill in the first $m$ positions of $\mathbf{x}$.
- $\mathsf{MHE\text{-}Rotate}_m(\mathbf{x})$ returns a cyclic-rotation of encrypted $\mathbf{x}$ by $m$ positions to the left.
- $\mathsf{MHE\text{-}ScalarMult}(s, \mathbf{x})$ multiplies all elements of encrypted $\mathbf{x}$ by an encrypted scalar value $s$.

We note that the protocols to switch between MHE and MPC as well as the MHE interactive bootstrapping protocol are invoked as needed and are omitted from protocol descriptions for simplicity.

Each of the collaborating parties $\mathsf{CP}_1, \ldots, \mathsf{CP}_k$ holds the corresponding portions of the horizontally distributed input genotype matrix $\bar{\mathbf{X}} \in \mathbb{R}^{(n\times m)} = (\bar{\mathbf{X}}_1, \ldots, \bar{\mathbf{X}}_k)$, covariate matrix $\bar{\mathbf{C}} \in \mathbb{R}^{(n\times c)} = (\bar{\mathbf{C}}_1, \ldots, \bar{\mathbf{C}}_k)$ and phenotype vector $\bar{\mathbf{y}} \in \mathbb{R}^{(n\times 1)} = (\bar{\mathbf{y}}_1, \ldots, \bar{\mathbf{y}}_k)$. The constant $n$ denotes the total number of individuals across all parties, $m$ the number of SNPs, and $c$ the number of covariates.

**Phase 1: Quality control (QC).** The federated approach of SF-GWAS allows individuals in the dataset to be filtered based solely on local calculations on the unencrypted input data. The pseudocode for our QC protocol is provided in Protocol 3. For the binary block format, SF-GWAS performs the sample filtering based on genotype missing rate and heterozygosity. For the PGEN format, SF-GWAS additionally takes a sample filter file as input, since programs like PLINK can be easily used to perform the filtering in advance. In both cases, SF-GWAS filters the SNPs based on missing rate, minor allele frequency (MAF), and Hardy-Weinberg equilibrium (HWE), all of which are computed globally including all parties' data. Unlike S-GWAS [1], SF-GWAS requires the parties to locally compute the aggregate statistics using the unencrypted data (see lines 2-3 in Protocol 3) before invoking MPC routines for joint filtering, which greatly improves computational efficiency. Note that, after each party $\mathsf{CP}_i$ has computed a local statistic $x_i$ for a particular SNP (e.g., the number of missing values), a secret sharing of the global statistic $[\sum_i x_i]$ can be immediately constructed by setting $[\sum_i x_i]_j = x_j$ for each $\mathsf{CP}_j$. Based on these global statistics, we apply filtering by calling MPC protocols for comparison and division (where the latter is required to compute quantities like the HWE test statistic based on the aggregated statistics, see lines 5-8 in Protocol 3). While a dataset with a large number of SNPs can incur a substantial computational cost in this step, SF-GWAS effectively parallelizes the MPC routines over blocks of SNPs to speedup the computation. The output of this step is a binary vector indicating which SNPs passed all QC filters, which is revealed to the parties so that subsequent analysis can be restricted to this smaller subset (line 10).

**Protocol 3** SF-GWAS: Quality Control (QC)

**Input:** Individual and SNP missing rate upper bounds, individual heterozygosity upper and lower bounds, minor allele frequency lower bound and Hardy-Weinberg equilibrium $\chi^2$ test statistic upper bound. For each party $p$, input genotypes $\bar{\mathbf{X}}_p^{(\hat{n}_p \times \hat{m})}$, covariates $\bar{\mathbf{C}}_p{}^{(c \times \hat{n}_p)}$, and phenotypes $\bar{\mathbf{y}}_p^{(\hat{n}_p)}$.
**Output:** For each party $p$, quality control-filtered dataset $\bar{\mathbf{X}}'_p$, $\bar{\mathbf{y}}'_p$, and $\bar{\mathbf{C}}'_p$.

1: Each party $p$:
2: Apply individual-level missingness and heterozygosity filters locally to obtain a filtered dataset.
3: Compute, in plaintext, missing value counts $\bar{\mathbf{m}}_p^{(\hat{m})}$, minor allele counts $\bar{\mathbf{a}}_p^{(\hat{m})}$, and genotype counts $\bar{\mathbf{G}}_p^{(\hat{m})}$ for all SNPs.
4: Invoke MPC-Aggregate on $\bar{\mathbf{m}}_p, \bar{\mathbf{a}}_p$, and $\bar{\mathbf{G}}_p$ to obtain secret shares of global aggregate counts $\mathbf{m}, \mathbf{a}$, and $\mathbf{G}$.
5: Invoke MPC-Comparison on $\mathbf{m}$ to compute the SNP missingness filter.
6: Invoke MPC-Comparison on $\mathbf{a}$ to compute the SNP MAF filter.
7: Invoke MPC-Divide and secure multiplications on $\mathbf{G}$ to compute the HWE $\chi^2$ statistics $\mathbf{z}$.
8: Invoke MPC-Comparison on $\mathbf{z}$ to compute the SNP HWE filter.
9: Securely multiply the three filters via MPC to take an intersection of passing SNPs and reveal the resulting filter.
10: Each party constructs a filtered input dataset $\bar{\mathbf{X}}'_p{}^{(n_p \times m)}$, $\bar{\mathbf{y}}'_p{}^{(n_p)}$, and $\bar{\mathbf{C}}'_p{}^{(c \times n_p)}$.

**Phase 2: Principal component analysis (PCA).** We follow the approach of S-GWAS [1] to implement a randomized PCA algorithm [14] for this step using our hybrid secure computation approach (Supplementary Note 4). Following standard practice, we perform PCA on a subset of SNPs with low levels of linkage disequilibrum (LD). While SF-GWAS imposes a minimum distance threshold on the QC-filtered SNPs to automatically obtain this reduced set, one may provide a different agreed-upon list of SNPs to use for PCA.

Our PCA protocol is summarized in Protocol 4. The algorithm begins by the computation of the average and inverse standard deviation for each column of the input data (see lines 1 to 5 in Protocol 4). The obtained global encrypted vectors are used throughout the protocol (lines 8, 9, 12, 16 and 19) to standardize the genotype matrix *on-the-fly*. Note that this procedure (introduced in Protocol 5) enables us to keep the genotype matrix in plaintext during the entire protocol and therefore to maximize the use of low-cost operations on local plaintext data (see Supplementary Note 4).

The genotype matrix is then multiplied with a random projection matrix to obtain a low-dimensional 'sketch' of the original matrix (line 7). The random projection matrix is independently sampled from a globally shared randomness, and the local projection is efficiently performed by each party using the plaintext data.

Subsequently, we invoke the *power iteration* procedure [1] (lines 14 to 23), which repeatedly multiplies the sketch with the full genotype matrix for an easier identification of the top principal components (PCs) in the subsequent steps. This step involves large-scale plaintext-ciphertext matrix multiplications and QR factorization of the multiplication results (for normalization purposes) at the end of each iteration. For both, the matrices are distributed across the parties, necessitating aggregation of local computation results in certain steps to emulate the intended global computation. We leverage an efficient block-wise matrix multiplication routine based on [11] to perform the local computation, and employ a novel MHE implementation of distributed QR factorization (described below in Protocol 6), which additionally uses secret sharing-based MPC routines for comparison and square root inverse operations. We set the number of power iterations to 20 by default, which we found to be consistently effective in all our experiments. This is in part because GWAS typically requires a small number of principal components (e.g., 10). A different number of iterations can be used if desired.

After the power iterations, eigendecomposition (line 25 in Protocol 4) is applied to the small covariance matrix of the resulting low-dimensional sketch, which we perform using our efficient MPC routines given the small size of the matrix (based on the protocol from [1]). The resulting eigenvectors are then multiplied with the sketch to construct the PCs of the original matrix (line 27). The final result includes ciphertexts encoding the top PCs of the genotype matrix, representing the inferred covariates of each sample capturing their genetic ancestry background. Further details of our secure federated algorithm for PCA can be found in our related publication [15].

---

**Protocol 4** SF-GWAS: Principal Component Analysis (PCA)

---

**Input:** Randomized PCA algorithm parameters $\rho = \psi + \alpha$ where $\psi$ is the desired number of principal components (PCs) and $\alpha$ the oversampling parameter, a public random sketch matrix $\mathbf{\Pi}^{(\rho\times n)} = (\bar{\mathbf{\Pi}}_1^{(\rho\times n_1)}, \ldots, \bar{\mathbf{\Pi}}_k^{(\rho\times n_k)})$ whose columns have a single non-zero entry at a random position with value $\pm 1$, and $t$ the number of power iterations. Each party $p$ holds a filtered plaintext genotype matrix $\bar{\mathbf{X}}_p^{(n_p\times \ell)}$ based on a reduced set of $\ell$ SNPs selected for PCA, where $n_p$ denotes the number of individuals in party $p$'s dataset.
**Output:** For each party $p$, an encrypted matrix $\mathbf{Q}_p^{(\psi\times n_p)}$ whose columns represent the projections of each individual onto the $\psi$ PCs computed jointly on the global genotype dataset $(\bar{\mathbf{X}}_1, \ldots, \bar{\mathbf{X}}_p)$.

1: Each party $p$:
2:    Compute, in plaintext, the local sum $\bar{\mathbf{a}}_p$ and the sum of squares $\bar{\mathbf{b}}_p$ of genotype values for the columns of $\bar{\mathbf{X}}_p$.
3: Invoke $\mathbf{a} \leftarrow \mathsf{MPC\text{-}Aggregate}(\bar{\mathbf{a}}_p)$ and $\mathbf{b} \leftarrow \mathsf{MPC\text{-}Aggregate}(\bar{\mathbf{b}}_p)$ to obtain the secret shares of the global sums.
4: Compute the averages by invoking $\mathbf{m} \leftarrow \mathsf{MPC\text{-}Multiply}(\mathbf{a}, 1/n)$ and $\mathbf{v} \leftarrow \mathsf{MPC\text{-}Multiply}(\mathbf{b}, 1/n)$. Note that $1/n$ is a public constant.
5: Invoke $\mathbf{s} \leftarrow \mathsf{MPC\text{-}SqrtInv}(\mathbf{v} - \mathsf{MPC\text{-}Multiply}(\mathbf{m}, \mathbf{m}))$ to compute the inverse standard deviation for columns of $\bar{\mathbf{X}}$.
6: Each party $p$:
7:    Compute local data projection in plaintext: $\bar{\mathbf{P}}_p^{(\rho\times\ell)} \leftarrow \bar{\mathbf{\Pi}}_p^{(\rho\times n_p)} \times \bar{\mathbf{X}}^{(n_p\times\ell)}$
8:    Correct for mean centering: $\mathbf{P}_p^{(\rho\times\ell)} \leftarrow \bar{\mathbf{P}}_p - \bar{\mathbf{\Pi}}_p \times \bar{\mathbf{1}}^{(n_p\times 1)} \times \mathbf{m}^{(1\times\ell)}$
9:    Multiply each row of $\mathbf{P}_p$ with $\mathbf{s}$ for standardization
10: Invoke $\mathbf{P} \leftarrow \mathsf{MHE\text{-}Aggregate}(\mathbf{P}_p)$
11: Each party $p$:
12:    Invoke $\mathbf{R}_p^{(\rho\times n_p)} \leftarrow \mathsf{MHE\text{-}MatrixMult\text{-}LazyNorm}(\mathbf{P}, \bar{\mathbf{X}}_p^T, \mathbf{a}, \mathbf{s}, \mathsf{false})$ // Protocol 5
13:    Invoke $\mathbf{R}_p^{(\rho\times n_p)} \leftarrow \mathsf{DQR}(\mathbf{R}_p)$ // Protocol 6
14: **for** $j = 0, \ldots, t-1$ **do** // Power iterations
15:    Each party $p$:
16:       Invoke $\mathbf{P}_p^{(\rho\times\ell)} \leftarrow \mathsf{MHE\text{-}MatrixMult\text{-}LazyNorm}(\mathbf{R}_p, \bar{\mathbf{X}}_p, \mathbf{a}, \mathbf{s}, \mathsf{true})$
17:    $\mathbf{P}^{(\rho\times\ell)} \leftarrow \mathsf{MHE\text{-}Aggregate}(\mathbf{P}_p)$
18:    Each party $p$:
19:       Invoke $\mathbf{R}_p^{(\rho\times n_p)} \leftarrow \mathsf{MHE\text{-}MatrixMult\text{-}LazyNorm}(\mathbf{P}, \bar{\mathbf{X}}_p^T, \mathbf{a}, \mathbf{s}, \mathsf{false})$
20:       **if** $j < t-1$ **do**
21:          Invoke $\mathbf{R}_p^{(\rho\times n_p)} \leftarrow \mathsf{DQR}(\mathbf{R}_p)$
22:       **end if**
23: **end for**
24: $\mathbf{Z}^{(\rho\times\rho)} \leftarrow \mathsf{MHE\text{-}Aggregate}(\mathbf{R}_p \times \mathbf{R}_p^T)$
25: $\mathbf{U}^{(\psi\times\rho)} \leftarrow \mathsf{MPC\text{-}EigenDecomp}(\mathbf{Z})[0 : \psi - 1, :]$ // Top $\psi$ eigenvectors
26: Each party $p$:
27:    Compute $\mathbf{Q}_p^{(\psi\times n_p)} \leftarrow \mathbf{U} \times \mathbf{R}_p$

---

The distributed QR factorization (Protocol 6) is a core subroutine for PCA which is repeatedly used (see lines 13 and 21 in Protocol 4) to obtain an orthogonal matrix from the input matrix. More precisely, the distributed and encrypted input matrix $\mathbf{A}$ is first repeatedly multiplied by the Householder matrix $\mathbf{H} = \bar{\mathbf{I}} - 2\mathbf{v}^T \times \mathbf{v}$ (see lines 1 to 32 in Protocol 6), where $\mathbf{v}$ is the Householder transformation (row) vector computed from the first row of $\mathbf{A}$. This transformation is recursively performed on the $(i, i)$ minors of $\mathbf{A}$ by discarding its first row and column. Here, the householder transformation (lines 3 to 18) transforms the input vector into a vector (of the same norm) with zeros in all coordinates except for the first. Following the standard technique, the norm of the input vector (computed in lines 3-5) is added to or subtracted from its first coordinate (line 10), depending on the sign of the first coordinate (determined in line 7) for numerical stability. Afterwards, the vector is normalized (line 17) to obtain the Householder transformation vector. The matrix $\mathbf{Q}$ is then computed (lines 33 to 43) and corresponds to the product of all Householder matrices $\mathbf{H}$.

Throughout the PCA execution, the parties interact to aggregate intermediate computation results among the parties (e.g., when summing up the partial matrix multiplication results to obtain the global result) as well as to refresh the ciphertexts (i.e., apply interactive bootstrapping) to restore the capacity for further multiplications.

**Protocol 5** MHE-MatrixMult-LazyNorm($\mathbf{M}, \bar{\mathbf{N}}, \mathbf{a}, \mathbf{s}, t$)

**Input:** An encrypted matrix $\mathbf{M}^{(a\times b)}$ and a cleartext matrix $\bar{\mathbf{N}}^{(b\times c)}$. A Boolean flag $t$ indicating the axis of operation: $\mathbf{a}$ and $\mathbf{s}$ are encrypted vectors including the means and inverse standard deviations, respectively, of the *columns* of $\bar{\mathbf{N}}$, if $t$ is true, and the *rows* of $\bar{\mathbf{N}}$, if $t$ is false.
**Output:** An encrypted matrix $\mathbf{R}^{(a\times c)}$ representing the result of multiplying $\mathbf{M}$ by the *standardized* $\bar{\mathbf{N}}$, i.e., mean-centered and divided by the standard deviations based on the values in $\mathbf{a}$ and $\mathbf{s}$. For efficiency, this is achieved in a lazy manner without directly standardizing the matrix.

**If** $t$ is true:
  Compute the ciphertext-plaintext matrix multiplication: $\mathbf{D}^{(a\times c)} \leftarrow \mathbf{M} \times \bar{\mathbf{N}}$
  Correct for mean centering: $\mathbf{E} \leftarrow \mathbf{D}^{(a\times c)} - ((\mathbf{M} \times \mathbf{1}^{(b\times 1)}) \times \mathbf{a}^{(1\times c)})$
  Multiply each row of $\mathbf{E}$ with $\mathbf{s}$ for standardization: $\mathbf{R}^{(a\times c)} \leftarrow \mathbf{E}^{(a\times c)} \times \mathbf{S}^{(c\times c)}$, where $\mathbf{S}$ is a diagonal matrix with $\mathbf{s}$ along the diagonal
**Else**:
  Multiply each row of $\mathbf{M}$ with $\mathbf{s}$ for standardization: $\mathbf{D}^{(a\times b)} \leftarrow \mathbf{M}^{(a\times b)} \times \mathbf{S}^{(b\times b)}$ where $\mathbf{S}$ is a diagonal matrix with $\mathbf{s}$ along the diagonal
  Compute the ciphertext-plaintext matrix multiplication: $\mathbf{E}^{(a\times c)} \leftarrow \mathbf{D} \times \bar{\mathbf{N}}$
  Correct for mean centering: $\mathbf{R}^{(a\times c)} \leftarrow \mathbf{E} - (\mathbf{D} \times \mathbf{a}^{(b\times 1)} \times \mathbf{1}^{(1\times c)})$
**End if**

**Protocol 6** Distributed QR Factorization (DQR($\cdot$))

**Input:** Each party $p$ holds an encrypted submatrix $\mathbf{A}_p$ of the global matrix $\mathbf{A}^{(\delta\times h)} = (\mathbf{A}_1^{(\delta\times h_1)}, \ldots, \mathbf{A}_k^{(\delta\times h_k)})$.
**Output:** Encrypted matrix $\mathbf{Q}^{(\delta\times h)}$ with orthogonal rows such that $\mathbf{A} = \mathbf{L} \times \mathbf{Q}$ for a lower-triangular matrix $\mathbf{L}$.

Each party initializes $\mathbf{H}_p \leftarrow \mathbf{0}^{(\delta\times h_p)}$
**for** $i = 0, \ \ldots, \ \delta - 1$ **do**
  // Compute Householder transformation vector $\mathbf{v}$
  Each party computes $z_p \leftarrow \mathbf{A}_p[0,:] \bullet \mathbf{A}_p[0,:]$
  $\|\mathbf{z}\|^2 \leftarrow$ MHE-Aggregate($z_p$)
  $\|\mathbf{z}\| \leftarrow$ MPC-Sqrt($\|\mathbf{z}\|^2$)
  Let $o$ and $\ell$ such that $\mathbf{A}[:,i] = \mathbf{A}_o[:,\ell]$
  $s \leftarrow$ MPC-IsPositive($\mathbf{A}_o[0,\ell]$)
  Assign $\mathbf{u}_p \leftarrow \mathbf{A}_p[0,:]$
  Party $o$:
    $u \leftarrow s \cdot \|\mathbf{z}\| + \mathbf{A}_o[0,\ell]$
    $\Delta \leftarrow u \cdot u - \mathbf{A}_o[0,\ell] \cdot \mathbf{A}_o[0,\ell]$
    $\mathbf{u}_o[\ell] \leftarrow u$
    Secret shares $\Delta$ with other parties
  $\|\mathbf{u}\|^2 \leftarrow \|\mathbf{z}\|^2 + \Delta$
  $\|\mathbf{u}\|^{-1} \leftarrow$ MPC-SqrtInv($\|\mathbf{u}\|^2$)
  Each party $p$:
    $\mathbf{v}_p \leftarrow$ MHE-ScalarMult($\|\mathbf{u}\|^{-1}, \mathbf{u}_p$)
    Assign $\mathbf{H}_p[i,:] = \mathbf{v}_p$
  // Update $\mathbf{A} \leftarrow \mathbf{A} - 2 \cdot \mathbf{A} \times \mathbf{v}^T \times \mathbf{v}$
  Each party $p$ computes $\mathbf{w}_p = \mathbf{A}_p \times \mathbf{v}_p^T$
  Invoke $\mathbf{w} \leftarrow$ MHE-Aggregate($\mathbf{w}_p$)
  Each party $p$:
    **for** $j = 0, \ \ldots, \ \delta - 1 - i$ **do**
      $\mathbf{A}_p[j,:] \leftarrow \mathbf{A}_p[j,:] - 2 \cdot$ MHE-ScalarMult($\mathbf{w}[j], \mathbf{v}_p$)
    **end for**
    // Remove first row and zero out column $i$ of $\mathbf{A}$
    **for** $j = \ 0, \ \ldots, \ \delta - i - 2$ **do**
      **if** $p = o$:
        $\mathbf{A}_p[j,:] \leftarrow (\mathbf{0}^{(1\times(\ell+1))}, \mathbf{A}_p[j+1, \ell+1:])$
      **else:**
        $\mathbf{A}_p[j,:] \leftarrow (\mathbf{A}_p[j+1,:])$
      **end if**
    **end for**
**end for**
Set $\mathbf{Q} = (\mathbf{Q}_1^{(\delta\times h_1)}, \ldots, \mathbf{Q}_P^{(\delta\times h_p)}) = [\bar{\mathbf{I}}^{(\delta\times\delta)} \ \ \mathbf{0}^{(\delta\times(h-\delta))}]$
**for** $i = \delta - 1, \ \ldots, 0$ **do**
  // Update $\mathbf{Q} \leftarrow \mathbf{Q} - 2 \cdot \mathbf{Q} \times \mathbf{v}^T \times \mathbf{v}$ for each Householder vector $\mathbf{v}$
  Each party $p$:
    $\mathbf{v}_p = \mathbf{H}_p[i,:]$
    $\mathbf{w}'_p = \mathbf{Q}_p \times \mathbf{v}_p^T$
  Invoke $\mathbf{w}' \leftarrow$ MHE-Aggregate($\mathbf{w}'_p$)
  Each party $p$:
    **for** $j = 0, \ \ldots, \ \delta - 1$ **do**
      $\mathbf{Q}_p[j,:] \leftarrow \mathbf{Q}_p[j,:] - 2 \cdot$ MHE-ScalarMult($\mathbf{w}'[j], \mathbf{v}_p$)
    **end for**
**end for**

**Phase 3: Association tests.** SF-GWAS supports both linear and logistic regression-based association tests as follows.

*Linear regression-based association tests.* SF-GWAS implements linear regression-based association tests by measuring the Pearson correlation coefficient between each genotype (minor allele dosages) and the phenotype with covariate correction. For binary traits, this is equivalent to the Cochran-Armitage trend test. To correct for confounding due to covariates, both the observed covariates (e.g., age and sex) and the principal components from the previous phase are projected out of the genotypes and phenotypes before computing the correlation (lines 1 to 10 in Protocol 7). This procedure involves computing the joint orthogonal basis of all covariates (another application of distributed QR factorization from the previous phase), then projecting the genotype vectors onto this basis (line 11). This latter step requires multiplication between the large-scale genotype matrix and the encrypted covariate basis (both distributed among the parties), which constitutes a bottleneck in this phase. Similar to the matrix multiplications in the PCA step, we leverage an efficient block-wise matrix multiplication routine, based on [11], to perform the required local computation of this step at the level of genomic blocks from the genotype matrix. Once the matrix multiplications are finished, the parties aggregate their local results to obtain the global multiplication results (line 12). Next, we compute the correlation coefficients through a sequence of MPC routines as in [1]. This involves invocations of square root inverse to compute the inverse standard deviations of each genotypes and the phenotype (lines 22 and 23); and a number of multiplications to calculate the correlation coefficient (line 24). The coefficients for all SNPs spanning the genome are then revealed as the final output of the algorithm. Note that, given the covariate-corrected correlation coefficient $r$, one can compute the corresponding $\chi^2$ statistic with one degree of freedom as $r^2(n-c)/(1-r^2)$, based on which a $p$-value can be obtained. $n$ and $c$ denote the total number of individuals and the number of covariates, respectively. Note that this mapping does not reveal any additional information other than $r$.

**Protocol 7** SF-GWAS: Linear Regression-based Association Tests

**Input:** Each party $p$ holds a subset of the plaintext input genotype matrix $\bar{\mathbf{X}}_p^{(n_p \times m)}$, a subset of the cleartext covariate matrix $\bar{\mathbf{C}}_p^{(c \times n_p)}$, and a subset of the cleartext phenotype vector $\bar{\mathbf{y}}_p^{(n_p \times 1)}$ and a subset of the encrypted PC covariate matrix $\mathbf{Q}_p^{(\psi \times n_p)}$, whose columns represent the projections of each individual onto the $\psi$ PCs computed jointly on the global genotype dataset.

**Output:** An encrypted vector $\mathbf{r}^{(1 \times m)}$ in which each element corresponds to the Pearson correlation association statistic for each of the $m$ variants.

1: Each party $p$ obtains $\mathbf{D}_p^{(c \times n_p)} \leftarrow \mathsf{DQR}(\bar{\mathbf{C}}_p)$
// Compute $\mathbf{E} = (\mathbf{I} - \mathbf{D}^T \times \mathbf{D}) \times \mathbf{Q}^T$
2: Invoke $\mathbf{U} \leftarrow$ MHE-Aggregate$(\mathbf{D}_p \times \mathbf{Q}_p^T)$
3: Each party $p$:
4: Computes $\mathbf{E}'_p \leftarrow \mathbf{Q}_p^T - (\mathbf{D}_p^T \times \mathbf{U})$
5: Collectively obtains $\mathbf{E}_p \leftarrow \mathsf{DQR}(\mathbf{E}'_p)$
6: Concatenate $\mathbf{Z}_p^{((c+\psi) \times n_p)} \leftarrow (\mathbf{D}_p, \mathbf{E}_p)$
// Project covariates out of $\mathbf{y}$:
$\mathbf{y}' = (\mathbf{I} - \mathbf{Z}^T \times \mathbf{Z}) \times \mathbf{y}$
7: Invoke $\mathbf{u} \leftarrow$ MHE-Aggregate$(\mathbf{Z}_p \times \mathbf{y}_p)$
8: Each party $p$ computes $\mathbf{y}'_p \leftarrow \mathbf{y}_p - (\mathbf{Z}_p^T \times \mathbf{u})$
// Compute association statistics
9: Each party $p$:
10: Concatenates $\mathbf{Z'}_p^{((c+\psi+1) \times n_p)} \leftarrow (\mathbf{Z}_p, \mathbf{y'}_p^T)$
11: Computes $\mathbf{W}_p{}^{((c+\psi+1) \times m)} \leftarrow \mathbf{Z}'_p \times \bar{\mathbf{X}}_p$
12: Invoke $\sigma_{xy}^{(1 \times m)} \leftarrow$ MHE-Aggregate$(\mathbf{W}_p[c+\psi, :])$
13: Each party $p$:
14: Initialize $\mathbf{b}_p \leftarrow \mathbf{0}^{(1 \times m)}$
15: **for** $i = 0, \ldots, c+\psi-1$ **do**
16: $\mathbf{b}_p \leftarrow \mathbf{b}_p + \mathbf{W}_p[i, :] \cdot \mathbf{W}_p[i, :]$
17: **end for**
18: $\bar{\mathbf{v}}_p^{(1 \times m)} \leftarrow \sum_i \bar{\mathbf{X}}_p[i, :] \cdot \bar{\mathbf{X}}_p[i, :]$
19: $\mathbf{s}_{x,p} \leftarrow \bar{\mathbf{v}}_p - \mathbf{b}_p$
20: Invoke $\mathbf{s}_x \leftarrow$ MHE-Aggregate$(\mathbf{s}_{x,p})$
21: Invoke $s_y \leftarrow$ MHE-Aggregate$(\mathbf{y}'_p \bullet \mathbf{y}'_p)$
22: Invoke $\sigma_x^{-1} \leftarrow$ MPC-SqrtInv$(\mathbf{s}_x)$
23: Invoke $\sigma_y^{-1} \leftarrow$ MHE-Duplicate$_m$(MPC-SqrtInv$(s_y)$)
24: Compute $\mathbf{r}^{(1 \times m)} \leftarrow \sigma_{xy} \cdot \sigma_x^{-1} \cdot \sigma_y^{-1}$

*Logistic regression-based association tests.* SF-GWAS evaluates the statistical significance of a SNP effect in the standard logistic regression model for GWAS using a score test approach. This strategy minimizes the burden of nonlinear computation under encryption by requiring only the baseline (null) covariate-only model to be estimated via logistic regression as opposed to training a separate model for every genetic variant being tested. Given the coefficients of the covariate-only model, the score test $t$-statistic for each SNP $i$ can be computed as

$$t = \frac{\tilde{\mathbf{g_i}}^T(\mathbf{y}-\mathbf{u})}{\sqrt{\tilde{\mathbf{g_i}}^T\mathbf{A}\tilde{\mathbf{g_i}}}},$$

where $\mathbf{y}$ is the phenotype vector, $\mathbf{u}$ the vector of estimated mean of the binary trait under the null model and $\mathbf{A} = \mathbf{diag}\{u_j(1-u_j)\}$ with $u_j$ the $j$-th element of $\mathbf{u}$ corresponding to the $j$-th individual. We denote $\tilde{\mathbf{y}} = (\mathbf{y}-\mathbf{u})$. The genotype residual vector for all individuals after adjusting for covariates is

$$\tilde{\mathbf{g_i}} = \mathbf{g}_i - \mathbf{D}^T(\mathbf{DAD}^T)^{(-1)}\mathbf{DA}\mathbf{g}_i,$$

where $\mathbf{g_i}$ is the genotype vector and $\mathbf{D}$ the matrix of all covariates (including the principal components). Therefore, the $t$-statistic for SNP $i$ can be expressed as

$$t = \frac{\mathbf{g}_i^T\tilde{\mathbf{y}} - \mathbf{g}_i^T\mathbf{AD}^T(\mathbf{DAD}^T)^{(-1)}\mathbf{D}\tilde{\mathbf{y}}}{\sqrt{\mathbf{g}_i^T\mathbf{A}\mathbf{g}_i - \mathbf{g}_i^T\mathbf{AD}^T(\mathbf{DAD}^T)^{(-1)}\mathbf{DA}\mathbf{g}_i}}. \tag{1}$$

SF-GWAS efficiently computes this statistic for all SNPs by first computing $\mathbf{A}$, which requires to train the null model, i.e., a logistic regression model trained with solely covariates as features and the phenotype as the label. The training of the null model is performed through the secure execution of Newton's Method, described in Protocol 8. Although second-order derivative methods like Newton's method often converge faster than first-order gradient-based methods, they have been avoided in previous secure federated solutions [16, 1] due to their inability to practically handle the secure execution of a matrix inversion. Instead, previous works typically rely on stochastic gradient descent (SGD) for its computational simplicity, each training iteration requiring only simple matrix-vector multiplications and the evaluation of the activation function. Conversely, each training iteration in Newton's method necessitates the computation of the gradient and the inverse of the Hessian matrix. SF-GWAS efficiently supports the matrix inversion by switching to the MPC domain with secret shares and leveraging our eigendecomposition protocol.

More precisely, Newton's method for logistic regression iteratively updates the parameter estimates by using first-order (i.e., gradient $L(\cdot)$) and second-order (i.e., Hessian $H(\cdot)$) derivatives of the log-likelihood function to find the maximum likelihood estimates. Given a matrix $\mathbf{Z}^{c\times n}$ with each column a sample of $c$ features, and a vector $\mathbf{y}^{(n\times 1)}$ of the corresponding labels, Newton's method iteratively finds the maximum-likelihood parameters $\mathbf{w}^{(c\times 1)}$ for the logistic GWAS null model by computing:

$$\mathbf{w}^{(t+1)} = \mathbf{w}^{(t)} - \left[H(\mathbf{w}^{(t)})\right]^{-1}\nabla L(\mathbf{w}^{(t)}) = \mathbf{w}^{(t)} - \left[\mathbf{ZAZ}^T\right]^{-1}\mathbf{Z}(\mathbf{y}-\mathbf{u}), \tag{2}$$

where $\mathbf{u}^{(n\times 1)} = \sigma(\mathbf{w}^T\mathbf{Z})$, with $\sigma(\cdot)$ indicating the sigmoid function, represents the vector of predicted probabilities (i.e., the vector of estimated mean of the binary trait under the null model), and $\mathbf{A}^{(n\times n)} = \mathbf{diag}(\mathbf{u})$ is a diagonal matrix with $\mathbf{u}$ on its diagonal.

Protocol 8 details our secure implementation of the Newton's method on a vertically-split, row-wise encrypted matrix $\mathbf{Z}^{(c\times n)}$ (i.e., each row is packed into one or more ciphertexts), which outputs an encrypted vector of estimated weights $\mathbf{w}^{(c\times 1)}$. The ComputeInvHessian subroutine is invoked in each iteration to compute the predicted probabilities (lines 9-10), the gradient (lines 11-13) and the inverse of the Hessian matrix (lines 14-19), based on the current parameters. The encrypted Hessian matrix $\mathbf{W} = \mathbf{ZAZ}^T$ is computed as the aggregation of the locally computed $\mathbf{W}_p$ (line 18). SF-GWAS then switches to the secret sharing domain to compute $\mathbf{W}^{(-1)}$ and its factorization $\mathbf{W}^{(-1)} = \mathbf{B}^T\mathbf{B}$ (line 19) based on eigendecomposition. This subroutine optionally returns additional variables for use in Protocol 9. The model parameters are updated in line 5 based on the gradient and the Hessian matrix.

We observed that Newton's method for the GWAS null model converges rapidly in a few epochs, e.g., less than 12 in all our experiments, whereas mini-batch SGD consistently requires more than 100 epochs, where an epoch represents a full pass over the training dataset over multiple iterations (Supplementary Fig. 10). For general applicability, we set the number of iterations for Newton's method to 15 by default.

**Protocol 8** Secure Federated Newton's Method for Logistic Regression (LR-NM)

**Input:** Each party $p$ holds a subset of the encrypted input matrix $\mathbf{Z}_p^{(c \times n_p)}$, and a subset of the cleartext label vector $\bar{\mathbf{y}}_p^{(n_p \times 1)}$. The learning parameter *iter* defines the number of epochs in Newton's Method. The ComputeInvHessian procedure returns the encrypted gradient and inverse of Hessian matrix, and the intermediate values that are used in Protocol 9.

**Output:** An encrypted vector of model coefficients $\mathbf{w}^{(c \times 1)}$.

1: Each party $p$:
2:   Initializes $\mathbf{w}^{(c \times 1)} \leftarrow \mathbf{0}^{(c \times 1)}$
3:   **for** $i = 0, \ldots, iter$ **do**
4:     $\mathbf{r}$, _, _, $\mathbf{W}^{(-1)}$, _, _ $\leftarrow$ ComputeInvHessian($\mathbf{Z}_p$, $\bar{\mathbf{y}}_p, \mathbf{w}$) // gradient $\mathbf{r}$, inverse Hessian $\mathbf{W}^{(-1)}$
5:     $\mathbf{w} \leftarrow \mathbf{w} + \mathbf{W}^{(-1)} \times \mathbf{r}$ // $\mathbf{w}$ is the same across all parties
6:   **end for**

7: **Procedure** ComputeInvHessian($\mathbf{Z}_p^{(c \times n_p)}$, $\bar{\mathbf{y}}_p^{(n_p \times 1)}, \mathbf{w}^{(c \times 1)}$)
8:   Each party $p$:
9:     Computes $\mathbf{i}_p^{(1 \times n_p)} \leftarrow \mathbf{w}^T \times \mathbf{Z}_p$
10:     Computes $\mathbf{u}_p^{(1 \times n_p)} \leftarrow \text{SigmoidApprox}(\mathbf{i}_p)$
11:     Computes $\tilde{\mathbf{y}}_p^{(1 \times n_p)} \leftarrow \bar{\mathbf{y}}_p - \mathbf{u}_p$
12:     Computes $\mathbf{r}_p^{(c \times 1)} \leftarrow \mathbf{Z}_p \times \tilde{\mathbf{y}}_p^T$
13:   Invoke $\mathbf{r} \leftarrow$ MHE-Aggregate($\mathbf{r}_p$)
14:   Each party $p$:
15:     Computes $\mathbf{A}_p^{(n_p \times n_p)} \leftarrow \text{diag}(\mathbf{u}_p - \mathbf{u}_p^2)$
16:     Computes $\mathbf{V}_p^{(c \times n_p)} \leftarrow \mathbf{Z}_p \times \mathbf{A}_p$
17:     Computes $\mathbf{W}_p^{(c \times c)} \leftarrow \mathbf{V}_p \times \mathbf{Z}_p^T$
18:   Invoke $\mathbf{W} \leftarrow$ MHE-Aggregate($\mathbf{W}_p$)
19:   Invoke $\mathbf{B}, \mathbf{W}^{(-1)} \leftarrow$ MPC-MatrixInvSqrt($\mathbf{W}$) // s.t. $\mathbf{B}^T\mathbf{B} = \mathbf{W}^{(-1)}$
20:   **return** $\mathbf{r}$, $\mathbf{V}_p$, $\mathbf{B}$, $\mathbf{W}^{(-1)}$, $\tilde{\mathbf{y}}_p$, $\mathbf{A}_p$

The next step is to compute the score test statistics for all SNPs based on the null model, as shown in Protocol 9. To perform this efficiently, SF-GWAS first computes the operations that are independent of the SNPs in Equation 1. It also computes only once all elements that are common between the numerator and denominator. Similarly as for the linear regression-based association tests, SF-GWAS first computes the joint orthogonal basis of all covariates (line 2 in Protocol 9). It then trains the null model (line 3) and evaluates it (line 4). The evaluation of the model is similar to another iteration of the training using the model parameters obtained before. This is performed using the ComputeInvHessian procedure (defined in Protocol 8). $\mathbf{W}^{(-1)} = (\mathbf{DAD}^T)^{(-1)}$ (the inverse of the Hessian matrix) and $\mathbf{V}_p = \mathbf{D}_p\mathbf{A}$ from Equation 1 are computed once in this procedure (line 4) and reused in the following steps. The subsequent matrix multiplications are ordered to minimize their complexity and the intermediate results are aggregated among the parties when they are of small dimensions (e.g., $\mathbf{d}$ in line 9). Until line 11, the operations are independent of the number of SNPs $m$. SF-GWAS performs the SNPs-dependent operations in chunks to enable parallelization and to limit the memory usage. We design the following steps to maximize ciphertext-plaintext multiplications (lines 12, 20, 24 and 25), which are more efficient than ciphertext-ciphertext. We also note that to efficiently compute the vector $\mathbf{b} = \mathbf{g}^T\mathbf{AD}^T(\mathbf{DAD}^T)^{(-1)}\mathbf{DAg} = \mathbf{g}^T\mathbf{VW}^{(-1)}\mathbf{V}^T\mathbf{g}$ (in the denominator of Equation 1), we first compute $\mathbf{H}_p$ such that $\mathbf{H}_p^T\mathbf{H}_p = \mathbf{b}$. $\mathbf{H}_p$ is obtained in line 12 through an efficient ciphertext-plaintext matrix multiplication, and $\mathbf{b}$ is efficiently computed by aggregating the squared rows of $\mathbf{H}$ (line 17) after aggregation among all parties (line 13). The final statistics for all SNPs are obtained by computing the inverse square root in the secret-sharing domain.

**Protocol 9** SF-GWAS: Logistic Regression-based Association Tests

**Input:** Each party $p$ holds a subset of the plaintext input genotype matrix $\bar{\mathbf{X}}_p^{(n_p \times m)}$, a subset of the cleartext covariate matrix $\bar{\mathbf{C}}_p^{(c \times n_p)}$, a subset of the cleartext phenotype vector $\bar{\mathbf{y}}_p^{(n_p \times 1)}$, a subset of the encrypted PC covariate matrix $\mathbf{Q}_p^{(\psi \times n_p)}$, whose columns represent the projections of each individual onto the $\psi$ PCs computed jointly on the global genotype dataset, and the number of epochs *iter* for Newton's Method (see Protocol 8).

**Output:** An encrypted vector $\mathbf{q}^{(1 \times m)}$ in which each element corresponds to the Score Test association statistic for each of the $m$ variants.

Each party $p$:
  Computes $\mathbf{D}_p^{((c+\psi) \times n_p)} \leftarrow [1, \mathsf{DQR}(\bar{\mathbf{C}}_p, \mathbf{Q}_p)[1:,:]]$
Invoke $\mathbf{w}^{((c+\psi)\times 1)} \leftarrow \mathsf{LR\text{-}NM}(\mathbf{D}_p, \bar{\mathbf{y}}_p, iter)$
Invoke $\_, \mathbf{V}_p, \mathbf{B}, \mathbf{W}^{(-1)}, \tilde{\mathbf{y}}_p, \mathbf{A}_p \leftarrow$ COMPUTE-INVHESSIAN$(\mathbf{D}_p, \bar{\mathbf{y}}_p, \mathbf{w})$
Each party $p$:
  Computes $\mathbf{U}_p^{(n_p \times (c+\psi))} \leftarrow \mathbf{V}_p^T \times \mathbf{W}^{(-1)}$
  Computes $\mathbf{E}_p^{(n_p \times (c+\psi))} \leftarrow \mathbf{V}_p^T \times \mathbf{B}^T$
  Computes $\mathbf{d}_p^{((c+\psi)\times 1)} \leftarrow \mathbf{D}_p \times \tilde{\mathbf{y}}_p$
Invoke $\mathbf{d} \leftarrow \mathsf{MHE\text{-}Aggregate}(\mathbf{d}_p)$
Each party $p$:
  Computes $\mathbf{o}_p^{(n_p \times 1)} \leftarrow \mathbf{U}_p \times \mathbf{d}$
  Computes $\mathbf{H}_p^{((c+\psi)\times m)} \leftarrow \mathbf{E}_p^T \times \bar{\mathbf{X}}_p$
Invoke $\mathbf{H} \leftarrow \mathsf{MHE\text{-}Aggregate}(\mathbf{H}_p)$
Each party $p$:
  Initializes $\mathbf{b}^{(1\times m)} = \mathbf{0}^{(1\times m)}$
  **For** $i = 0, \ldots, c+\psi-1$ **:**
    $\mathbf{b} \leftarrow \mathbf{b} + \mathbf{H}[i,:] \cdot \mathbf{H}[i,:]$
  **End for**
Each party $p$:
  Computes $\mathbf{x}_p^{(1\times m)} \leftarrow \mathbf{a}_p^{(1 \times n_p)} \times \bar{\mathbf{X}}_p^2$, with $\mathbf{a}_p$ the diagonal from $\mathbf{A}_p$
Invoke $\mathbf{x} \leftarrow \mathsf{MHE\text{-}Aggregate}(\mathbf{x}_p)$
Compute $\mathbf{e}^{(1\times m)} \leftarrow \mathbf{x} - \mathbf{b}$
Each party $p$:
  Computes $\mathbf{j}_p^{(1\times m)} \leftarrow \tilde{\mathbf{y}}_p \times \bar{\mathbf{X}}_p$
  Computes $\mathbf{s}_p^{(1\times m)} \leftarrow \mathbf{o}_p^T \times \bar{\mathbf{X}}_p$
Invoke $\mathbf{f}^{(1\times m)} \leftarrow \mathsf{MHE\text{-}Aggregate}(\mathbf{j}_p - \mathbf{s}_p)$
Invoke $\mathbf{q}^{(1\times m)} \leftarrow \mathsf{MPC\text{-}Mult}(\mathbf{f}, \mathsf{MPC\text{-}InvSqrt}(\mathbf{e}))$

We note that recent studies have suggested several approaches to further improve the power of logistic association tests in highly imbalanced case-control datasets—e.g., Firth or Saddle Point Approximation (SPA)-corrected logistic regression [17]. Further extending our secure federated techniques to support a wide variety of analysis needs is an important direction for future research.

### Supplementary Note 6: SF-GWAS Algorithm for LMM-based GWAS

In LMM-based GWAS, the effect of population structure and cryptic relatedness on the phenotype is captured as a *random* effect, whose covariance is determined by the genetic relatedness among individuals in the cohort. In the following we introduce LMM-based association studies and describe an efficient stacked ridge regression approach for LMMs called REGENIE [17]. We then describe the main challenges of supporting the LMM-based workflow in a federated manner and introduce our scalable algorithm for secure federated LMM-based GWAS.

**Linear mixed model (LMM) association studies.** We start by formally describing the LMMs. LMMs model the target phenotype vector $\mathbf{y}$ of length $n$ individuals using the following linear model

$$\mathbf{y} = \beta_{\text{test}} \mathbf{x}_{\text{test}} + \mathbf{C}\boldsymbol{\alpha} + \mathbf{g} + \mathbf{e},$$

where $\mathbf{x}_{\text{test}}$ is a (column) vector of minor allele dosages of the variant being tested across $n$ individuals, $\mathbf{C}$ is a $n$-by-$c$ matrix of observed covariates where $c$ denotes the number of covariates, $\mathbf{g}$ represents the ambient genetic effect, and $\mathbf{e}$ represents the environmental effect. Both $\mathbf{x}_{\text{test}}$ and $\mathbf{y}$ are standardized to have zero mean and unit variance. We let $n$ and $m$ be the number of individuals and the number of variants in the dataset, respectively.

In this model, $\beta_{\text{test}}$ and $\boldsymbol{\alpha}$ describe the fixed effect sizes associated with the tested variant and the covariates, respectively, whereas $\mathbf{g}$ and $\mathbf{e}$ are modeled as random effects (hence the term "mixed" model). Under the standard infinitesimal model, which posits that the genetic effect on phenotype consists of many small effect-size variants, we can express

$$\mathbf{g} = \mathbf{X}_{\text{LOCO}} \boldsymbol{\beta}$$

where $\mathbf{X}_{\text{LOCO}}$ is a $n$-by-$m_{\text{LOCO}}$ matrix consisting of the standardized genotypes of $m_{\text{LOCO}}$ variants used to model the genetic effect based on the standard leave-one-chromosome-out (LOCO) scheme, which excludes all variants in the same chromosome as the tested variant in order to avoid the effects of linkage disequilibrium. These ambient variants are associated with effect sizes $\boldsymbol{\beta} \sim \mathcal{N}(\mathbf{0}, (\sigma_g^2/m_{\text{LOCO}})\mathbf{I}^{(m_{\text{LOCO}} \times m_{\text{LOCO}})})$, inducing a distribution over the genetic effect as $\mathbf{g} \sim \mathcal{N}(0, \sigma_g^2 \mathbf{K})$. $\mathbf{K} = \mathbf{X}_{\text{LOCO}} \mathbf{X}_{\text{LOCO}}^T / m_{\text{LOCO}}$ is referred to as the genetic relatedness (or empirical kinship) matrix. The environmental effect is modeled as $\mathbf{e} \sim \mathcal{N}(0, \sigma_e^2 \mathbf{I}^{(n \times n)})$. Note that $\sigma_g$ and $\sigma_e$ represent the variances of the polygenic and environmental components. The goal of the association test is to test the null hypothesis $H_0 : \beta_{\text{test}} = 0$.

A standard technique [17, 18] is to project the covariates out of the phenotypes and genotypes to simplify the computation. This results in a modified model

$$\tilde{\mathbf{y}} = \beta_{\text{test}} \tilde{\mathbf{x}}_{\text{test}} + \tilde{\mathbf{X}}_{\text{LOCO}} \boldsymbol{\beta} + \mathbf{e}, \tag{3}$$

where $\tilde{\mathbf{y}} = \mathbf{P}\mathbf{y}$, $\tilde{\mathbf{x}} = \mathbf{P}\mathbf{x}$, and $\tilde{\mathbf{X}}_{\text{LOCO}} = \mathbf{P}\mathbf{X}_{\text{LOCO}}$ with $\mathbf{P} = \mathbf{I}^{(n \times n)} - \mathbf{C}(\mathbf{C}^T \mathbf{C})^{-1} \mathbf{C}^T$. Note that $\mathbf{P}$ projects a vector onto the null space of $\mathbf{C}$.

The LMM-based $\chi^2$ test statistic is given by

$$\chi^2 = \frac{(\tilde{\mathbf{x}}_{\text{test}}^T \mathbf{V}_{\text{LOCO}}^{-1} \tilde{\mathbf{y}})^2}{\tilde{\mathbf{x}}_{\text{test}}^T \mathbf{V}_{\text{LOCO}}^{-1} \tilde{\mathbf{x}}_{\text{test}}}, \tag{4}$$

where $\mathbf{V}_{\text{LOCO}} = \hat{\sigma}_g^2 \mathbf{K} + \hat{\sigma}_e^2 \mathbf{I}^{(n \times n)}$ given the estimates $\hat{\sigma}_g$ and $\hat{\sigma}_e$ of the variance parameters.

**Review of REGENIE: Efficient stacked ridge regression for LMMs.** Computing the LMM association statistics is a computationally expensive task, in part because the maximum likelihood estimation of the variance parameter $\sigma_g$ involves costly matrix operations involving the $n$-by-$n$ genetic relatedness matrix (GRM), which becomes prohibitively large for large-scale datasets. Much of the prior algorithmic development efforts have focused on speeding up the use of GRM, e.g. by exploiting a factorization of the matrix [19]. A recent algorithm called REGENIE [17] introduced a different strategy based on *stacked ridge regression*, resulting in significant scalability improvements for LMM-based association tests, while achieving comparable accuracy to existing state-of-the-art tools such as BOLT-LMM [18], fastGWA [20], and SAIGE [21]. Moreover, we discovered that REGENIE's approach is more amenable to efficient computation over distributed datasets, which leads to its practical performance in SF-GWAS.

In REGENIE [17], the whole-genome regression model in Equation 3 is estimated in two phases by first regressing out the contribution of $\tilde{\mathbf{X}}_{\text{LOCO}}$ from $\tilde{\mathbf{y}}$, then fitting $\beta_{\text{test}}$ on the residuals to test for association. To further reduce the cost of regression over the large genome-wide matrix $\tilde{\mathbf{X}}_{\text{LOCO}}$, REGENIE employs a stacked ridge regression approach in the following two steps, referred to as Level 0 and Level 1.

Level 0: The genotype matrix is first split into $B$ contiguous blocks of $T$ variants each:

$$\tilde{\mathbf{X}}_{\text{LOCO}} = (\tilde{\mathbf{X}}_{\text{LOCO}}^{(1)}, \tilde{\mathbf{X}}_{\text{LOCO}}^{(2)}, \ldots, \tilde{\mathbf{X}}_{\text{LOCO}}^{(B)}).$$

Then for each block $b \in [B]$ and different choices of the regularization parameter $\lambda_r \in \{\lambda_1, \ldots, \lambda_R\}$, where $R$ represents the number of regularization parameters being tested, the following solution to the ridge regression problem $\tilde{\mathbf{y}} \approx \tilde{\mathbf{X}}_{\text{LOCO}}^{(b)} \boldsymbol{\beta}$ is computed.

$$\hat{\boldsymbol{\beta}}_{\lambda_r}^{(b)} := ((\tilde{\mathbf{X}}_{\text{LOCO}}^{(b)})^T \tilde{\mathbf{X}}_{\text{LOCO}}^{(b)} + \lambda_r \mathbf{I}^{(n \times n)})^{-1} (\tilde{\mathbf{X}}_{\text{LOCO}}^{(b)})^T \tilde{\mathbf{y}}, \tag{5}$$

$$\hat{\mathbf{y}}_{\text{LOCO}}^{(b,r)} := \tilde{\mathbf{X}}_{\text{LOCO}}^{(b)} \hat{\boldsymbol{\beta}}_{\lambda_r}^{(b)}. \tag{6}$$

The $\hat{\mathbf{y}}_{\text{LOCO}}^{(b,r)}$ is referred to as the *predictors*, representing the best polygenic prediction of the phenotype within a given genomic region, accounting for genotype correlations.

Level 1: The local predictors from Level 0 are aggregated to form a $n$-by-$BR$ global feature matrix

$$\mathbf{W} := (\hat{\mathbf{y}}_{\text{LOCO}}^{(1,1)}, \ldots, \hat{\mathbf{y}}_{\text{LOCO}}^{(B,R)}). \tag{7}$$

Then another around of ridge regression is performed (with $K$-fold cross validation to choose the optimal regularization parameter $\eta$) to obtain the following genome-wide phenotype predictions:

$$\hat{\mathbf{y}}_{\text{LOCO}} := \mathbf{W}(\mathbf{W}^T\mathbf{W} + \eta\mathbf{I}^{(BR\times BR)})^{-1}\mathbf{W}^T\tilde{\mathbf{y}}. \tag{8}$$

Given this global predictor as a proxy for ambient genetic effect, the $\chi^2$ statistic (with one degree of freedom) for the variant being tested in Equation 4 can now be formulated as

$$\chi^2 = \frac{(\tilde{\mathbf{x}}_{\text{test}}^T(\tilde{\mathbf{y}} - \hat{\mathbf{y}}_{\text{LOCO}}))^2}{\hat{\sigma}_e^2 \cdot (\tilde{\mathbf{x}}_{\text{test}}^T\tilde{\mathbf{x}}_{\text{test}})}, \tag{9}$$

where $\hat{\sigma}_e^2 = \|\tilde{\mathbf{y}} - \hat{\mathbf{y}}_{\text{LOCO}}\|_2^2/(n-c)$ is the estimated variance of the environmental effect.

The above approach substantially improves the speed of LMM computation by decomposing the problem into $B$ separate local ridge regression problems, which can be performed in parallel in high performance computing environments. Importantly, by formulating the problem as a series of ridge regression tasks, the problem becomes more tractable for secure federated computation, an aspect we exploit in our design of SF-GWAS to achieve efficiency.

**Key challenges of federated LMM computation.** To motivate our novel algorithmic techniques, we first describe the computational challenges in distributing the LMM computation across multiple data holders. Recall that most existing LMM-based GWAS algorithms account for population structure by using the $n$-by-$n$ genetic relatedness matrix (GRM) $\mathbf{K}$, which intuitively captures how individuals within a dataset are related to one another. A core computational step in the estimation of variance parameters or the calculation of association statistics is calculating a quantity of the form

$$(\mathbf{K} + \lambda\mathbf{I}^{(n\times n)})^{-1}\mathbf{v} = (\tilde{\mathbf{X}}\tilde{\mathbf{X}}^T/m + \lambda\mathbf{I}^{(n\times n)})^{-1}\mathbf{v}, \tag{10}$$

for some length-$n$ vector $\mathbf{v}$. Because the size of $\mathbf{K}$ scales with the number of individuals in the dataset, for large-scale GWAS, this computation inherently incurs an overwhelming computational cost. As such, addressing this challenge has been the focus of recent algorithmic development efforts for LMM (e.g., BOLT-LMM [18]).

In our setting, the fact that the off-diagonal blocks of $\mathbf{K}$ describe relatedness between individuals in *different* collaborating sites introduces a unique difficulty in distributing the computation. When naïvely implemented, those interactive terms in $\mathbf{K}$ are bound to require heavy communication among parties to account for their contributions. Even recently proposed iterative approaches for efficiently solving this linear system of equations without the inverse (e.g., conjugate gradient descent used by BOLT-LMM [18]) presents a similar challenge, as it involves repeated multiplications of $\mathbf{K}$ with a candidate solution vector. Moreover, we note that $\mathbf{K}$ is typically defined over covariate-corrected genotypes $\tilde{\mathbf{X}} = \mathbf{P}\mathbf{X}$, where $\mathbf{P}$ denotes a projection matrix for removing the covariate effect, which introduces another layer of entanglement between the private datasets at different sites, making federated computation further challenging.

**Our approach: secure federated ridge regression with covariates.** For SF-GWAS, we approach this problem differently, following the approach of REGENIE [17]. Instead of using the notion of a GRM, we train ridge regression models for local genomic windows to use as a proxy for ambient genetic effect on phenotype. Correcting for these polygenic predictions by testing for a variant's association with the phenotype residuals, our approach implicitly accounts for population structure under the LMM formulation.

Our work identifies this alternative approach for LMMs as a key enabling factor for federated computation. Let $\tilde{\mathbf{X}}^{(b)}$ be a subset of columns from a genotype matrix corresponding to a genomic block with $m_b$ variants. The ridge regression problem for LMM reduces to computing the expression

$$((\tilde{\mathbf{X}}^{(b)})^T\tilde{\mathbf{X}}^{(b)} + \lambda\mathbf{I}^{(m_b\times m_b)})^{-1}(\tilde{\mathbf{X}}^{(b)})^T\tilde{\mathbf{y}}. \tag{11}$$

In contrast to Equation 10, we see that the inverse operation is for a matrix of size $m_b$-by-$m_b$, which is significantly smaller than $\mathbf{K}$ (note $m_b$ is typically 1000). Furthermore, for horizontally distributed $\tilde{\mathbf{X}}^{(b)}$, where each party $p$ holds a subset of rows in this matrix denoted $\tilde{\mathbf{X}}_p^{(b)}$, we have the following decomposition

$$(\tilde{\mathbf{X}}^{(b)})^T\tilde{\mathbf{X}}^{(b)} = \sum_{p=1}^{k}(\tilde{\mathbf{X}}_p^{(b)})^T\tilde{\mathbf{X}}_p^{(b)}. \tag{12}$$

This property allows SF-GWAS to distribute the computation more efficiently across the parties and maximally leverage plaintext data that is available locally. In the following section, we describe how we exploit this insight to design secure and federated algorithms for conjugate gradient descent (CGD) and alternating direction method of multipliers (ADMM) approaches for ridge regression, which are used by SF-GWAS to carry out the Level 1 and Level 0 steps of REGENIE, respectively.

As explained before (see Equations 5 to 8), stacked ridge regression approach to LMM involves solving the following two ridge regression problems in Levels 0 and 1.

$$\hat{\boldsymbol{\beta}}^{(b)}_{\lambda_r} := ((\tilde{\mathbf{X}}^{(b)}_{\text{LOCO}})^T\tilde{\mathbf{X}}^{(b)}_{\text{LOCO}} + \lambda_r\mathbf{I}^{(n\times n)})^{-1}(\tilde{\mathbf{X}}^{(b)}_{\text{LOCO}})^T\tilde{\mathbf{y}}, \qquad (\textbf{Level 0}) \qquad (13)$$

$$\hat{\mathbf{y}}_{\text{LOCO}} := \mathbf{W}(\mathbf{W}^T\mathbf{W} + \eta\mathbf{I}^{(BR\times BR)})^{-1}\mathbf{W}^T\tilde{\mathbf{y}}. \qquad (\textbf{Level 1}) \qquad (14)$$

The first is solved $KBR$ times for each pair of a block $b \in [B]$ and a regularization parameter $\lambda_r$ for $r \in [R]$ using $K$-fold cross validation, and the second is solved $KR$ times for each of $R$ values of $\eta$ using the same $K$-fold cross validation. The challenging step in both is the multiplication by the inverse matrix, which is infeasible to solve directly when the matrix is only available in encrypted form. This is unlike REGENIE, which explicitly solves for the inverse using eigenfactorization. Explicitly computing the inverse for a large, homomorphically encrypted matrix imposes a considerable computational burden, which we aim to avoid. To address this, we develop two algorithms described below.

**Secure federated conjugate gradient descent (CGD).** Conjugate gradient descent (CGD) [22] is a well-known iterative algorithm for solving a system of linear equations without explicitly constructing the inverse of the design matrix. Notably, BOLT-LMM heavily utilizes the CGD algorithm to avoid working with the inverse of GRM. A requirement of CGD is that the design matrix be positive definite and well conditioned; in our setting, the regularization term in ridge regression ensures this property [23]. Hence, CGD can be applied to any of the ridge regression problems in our task. The central step in CGD is a multiplication of a candidate solution vector with the design matrix (not the inverse), which lends itself to efficient distributed computation. We outline our federated CGD algorithm in Protocol 10.

---

**Protocol 10** Secure Federated Conjugate Gradient Descent (CGD) for Ridge Regression

---

**Input:** Number of parties $k$, horizontally distributed input matrix $\mathbf{A} = (\mathbf{A}_1, \ldots, \mathbf{A}_k)$, target vector $\mathbf{b}$, regularization parameter $\lambda$, number of iterations $\tau$. Input data can be either plaintext or ciphertext.

**Ouput:** A vector $\mathbf{x}$ that satisfies $(\mathbf{A}^T\mathbf{A} + \lambda\mathbf{I})\mathbf{x} \approx \mathbf{b}$.

Initialize: $\mathbf{x}_0 \leftarrow \mathbf{0}$, $\mathbf{y}_0 \leftarrow \mathbf{b}$, $\mathbf{r}_0 \leftarrow \mathbf{b}$
**for** $j \in \{0, \ldots, \tau - 1\}$ **do**
  Each party $p$ locally computes $\mathbf{z}_p \leftarrow \mathbf{A}_p^T\mathbf{A}_p\mathbf{y}_j$
  $\mathbf{z} \leftarrow$ MHE-Aggregate($\mathbf{z}_p$)
  $\mathbf{u} \leftarrow \mathbf{z} + \lambda\mathbf{y}_j$ // $\mathbf{u} = (\mathbf{A}^T\mathbf{A} + \lambda\mathbf{I})\mathbf{y}_j$
  $\alpha \leftarrow$ MPC-Divide($\mathbf{r}_j^T\mathbf{r}_j$, $\mathbf{y}_j^T\mathbf{z}$)
  $\mathbf{x}_{j+1} \leftarrow \mathbf{x}_j + \alpha\mathbf{y}_j$
  $\mathbf{r}_{j+1} \leftarrow \mathbf{r}_j - \alpha\mathbf{z}$
  $\beta \leftarrow$ MPC-Divide($\mathbf{r}_{j+1}^T\mathbf{r}_{j+1}$, $\mathbf{r}_j^T\mathbf{r}_j$)
  $\mathbf{y}_{j+1} \leftarrow \mathbf{r}_j + \beta\mathbf{y}_j$
**end for**
**return** $\mathbf{x}_\tau$

---

We highlighted in blue the key step enabling the federated approach; we securely execute the rest of the algorithm using our secure computation framework. As explained in the previous section, the fact that the design matrix $\mathbf{A}^T\mathbf{A}$ in our setting decomposes as a sum of local design matrices $\mathbf{A}_p^T\mathbf{A}_p$ allows this step to be performed independently, then aggregated after both multiplications (the MHE-Aggregate step). Thus, the required communication scales with the number of predictive features (variants), not the number of samples held by each party, thereby offering better scaling to large datasets. To apply this algorithm to encrypted datasets, each of the calculations, namely matrix-vector multiplication, inner products, and addition/subtraction, are implemented using HE routines, with the exception of division, for which we switch to a secret sharing-based MPC routine. Note that the MHE-Aggregate step involves each party broadcasting their share to others and adding up all shares (homomorphically), which does not involve any decryption. For efficiency, this procedure is implemented over a star network where a central coordinator aggregates all data and in turn relays the result to all parties.

**Secure federated ADMM for ridge regression with covariates.** Although CGD offers a natural federated solution for ridge regression, it is still computationally burdensome for large input matrices. For instance, consider applying CGD to Level 0 of REGENIE, where the input is given as

$$\mathbf{A} = \tilde{\mathbf{X}}^{(b)} = (\tilde{\mathbf{X}}_1^{(b)}, \ldots, \tilde{\mathbf{X}}_k^{(b)}). \tag{15}$$

When each party performs the following local computation in Protocol 10

$$\mathbf{z}_p \leftarrow (\tilde{\mathbf{X}}_p^{(b)})^T \tilde{\mathbf{X}}_p^{(b)} \mathbf{y}_j, \tag{16}$$

they first need to multiply $\mathbf{y}_j$ with $\tilde{\mathbf{X}}_p^{(b)}$, which has an output dimension of $n_p$ (number of individuals in party $p$'s dataset), followed by another multiplication with $(\tilde{\mathbf{X}}_p^{(b)})^T$, finally resulting in a vector of length $m_b$. This is due to the fact that $(\tilde{\mathbf{X}}_p^{(b)})^T \tilde{\mathbf{X}}_p^{(b)}$ cannot be precomputed in plaintext, since $\tilde{\mathbf{X}}_p^{(b)}$ requires covariate correction involving all parties covariate data. Note that $m_b$, the block size, is a user parameter typically set to a small value (e.g. 1000) whereas $n_p$ can grow much larger for large-scale datasets. Therefore, CGD does not benefit from any dimension reduction (to $m_b$) that is otherwise exhibited in the plaintext formulation.

SF-GWAS overcomes this challenge by leveraging the alternating direction method of multipliers (ADMM) technique [24], which is a powerful method for transforming convex optimization problems into distributed optimization problems that can be more efficiently solved. Intuitively, ADMM relaxes the global objective by decoupling the terms involving each individual dataset, which in turn can be jointly optimized using local update equations that, in our case, involve plaintext matrices of size $m_b$ as desired. Our techniques build upon a recent work in security literature [25], which introduced a secure multiparty ADMM algorithm for distributed linear regression. Our work extends this work to the setting where the design matrix must be covariate-corrected, which introduces additional challenges as we describe below.

Here we describe how we apply ADMM to the ridge regression problem in Level 0 of REGENIE. Recall that the ridge regression of $\tilde{\mathbf{y}}$ onto $\tilde{\mathbf{X}}^{(b)}$ with regularization parameter $\lambda$ can be equivalently formulated as the following optimization problem:

$$\text{minimize}_{\mathbf{w}} \quad \frac{1}{2}\|\tilde{\mathbf{X}}^{(b)}\mathbf{w} - \tilde{\mathbf{y}}\|_2^2 + \frac{1}{2}\lambda\|\mathbf{w}\|_2^2. \tag{17}$$

To apply ADMM, we first decouple the two terms using a slack variable $\mathbf{z}$ with an equality constraint as follows.

$$\text{minimize}_{\mathbf{w},\mathbf{z}} \quad \frac{1}{2}\|\tilde{\mathbf{X}}^{(b)}\mathbf{w} - \tilde{\mathbf{y}}\|_2^2 + \frac{1}{2}\lambda\|\mathbf{z}\|_2^2, \tag{18}$$

$$\text{s.t.} \quad \mathbf{w} - \mathbf{z} = 0. \tag{19}$$

Next, we note that the first objective term can be written as a sum of squared loss computed over each party's dataset as

$$\|\tilde{\mathbf{X}}^{(b)}\mathbf{w} - \tilde{\mathbf{y}}\|_2^2 = \sum_{p=1}^{P} \|\tilde{\mathbf{X}}_p^{(b)}\mathbf{w} - \tilde{\mathbf{y}}_p\|_2^2, \tag{20}$$

where we partition $\tilde{\mathbf{y}} = [\tilde{\mathbf{y}}_1, \ldots, \tilde{\mathbf{y}}_k]$ in the same manner as $\tilde{\mathbf{X}}^{(b)}$. Finally, further decoupling the $\mathbf{w}$ parameters across parties for distributed optimization, we obtain

$$\text{minimize}_{\mathbf{w}_p,\mathbf{z}} \quad \frac{1}{2}\sum_{p=1}^{P} \|\tilde{\mathbf{X}}_p^{(b)}\mathbf{w}_p - \tilde{\mathbf{y}}_p\|_2^2 + \frac{1}{2}\lambda\|\mathbf{z}\|_2^2, \tag{21}$$

$$\text{s.t.} \quad \mathbf{w}_p - \mathbf{z} = 0, \forall p. \tag{22}$$

The resulting iterative optimization procedure based on the standard ADMM derivation, for general local matrices $\mathbf{A}_1, \ldots, \mathbf{A}_k$, is shown in Protocol 11.

**Protocol 11** ADMM Algorithm for Ridge Regression (our adaptation of [25])

**Input:** Number of parties $k$, horizontally distributed input matrix $\bar{\mathbf{A}} = (\bar{\mathbf{A}}_1, \ldots, \bar{\mathbf{A}}_k)$, target vector $\mathbf{b} = (\mathbf{b}_1, \ldots, \mathbf{b}_k)$, regularization parameter $\lambda$, learning rate $\rho$, number of iterations $\tau$. An overline indicates a plaintext variable.
**Ouput:** A vector $\mathbf{z}$ that satisfies $(\mathbf{A}^T\mathbf{A} + \lambda\mathbf{I})\mathbf{z} \approx \mathbf{b}$.

Each party $p$:
  Locally computes $\bar{\mathbf{R}}_p^{-1} \leftarrow (\bar{\mathbf{A}}_p^T\bar{\mathbf{A}}_p + \rho\bar{\mathbf{I}})^{-1}$
  Initializes $\mathbf{w}_p^0 \leftarrow 0, \mathbf{u}_p^0 \leftarrow 0$ and a global vector $\mathbf{z}^0 \leftarrow 0$
**for** $j \in \{0, ..., \tau - 1\}$ **do**
  Each party $p$ locally computes $\mathbf{w}_p^{j+1} \leftarrow \bar{\mathbf{R}}_p^{-1}(\mathbf{b} + \rho\mathbf{z}^j - \mathbf{u}_p^j)$
  $\bar{\mathbf{w}}^{j+1} \leftarrow \textsf{MHE-Aggregate}(\mathbf{w}_p^{j+1})/k$
  $\bar{\mathbf{u}}^{j} \leftarrow \textsf{MHE-Aggregate}(\mathbf{u}_p^{j})/k$
  Each party $p$ locally computes:
    $\mathbf{z}^{j+1} \leftarrow (\rho\bar{\mathbf{w}}^{j+1} + \bar{\mathbf{u}}^j)/(\lambda/k + \rho)$
    $\mathbf{u}_p^{j+1} \leftarrow \mathbf{u}_p^j + \rho(\mathbf{w}_p^{j+1} - \mathbf{z}^{j+1})$
**end for**
**return** $\mathbf{z}^\tau$

However, we note that our given $\mathbf{A}_p = \tilde{\mathbf{X}}_p^{(b)}$ is not available in plaintext, as it is meant to be standardized and covariate-corrected based on the global matrix $\tilde{\mathbf{X}}^{(b)}$. Therefore, although each party has access to their own raw genotype martix $\mathbf{X}_p^{(b)}$, they are not able to precompute the following matrix shown in Protocol 11 in plaintext:

$$\mathbf{R}_p^{-1} = ((\tilde{\mathbf{X}}_p^{(b)})^T\tilde{\mathbf{X}}_p^{(b)} + \rho\mathbf{I}^{(m_b \times m_b)})^{-1}. \tag{23}$$

In SF-GWAS, we introduce a technique to resolve this issue by using the Woodbury matrix identity [26] to perform covariate correction in the computation of $\mathbf{R}_p^{-1}$ on the fly as follows. First, recall that

$$\tilde{\mathbf{X}}^{(b)} = (\mathbf{I}^{(n\times n)} - \mathbf{C}(\mathbf{C}^T\mathbf{C})^{-1}\mathbf{C}^T)\mathbf{X}^{(b)}\mathbf{S}^{(b)}, \tag{24}$$

where $\mathbf{C}$ is an $n$-by-$c$ covariate matrix, and $\mathbf{S}$ represents a diagonal matrix with inverse standard deviations for each column of $\mathbf{X}^{(b)}$. We consider the setting where $\mathbf{S}$ is jointly computed during quality control and shared among the parties to facilitate the LMM analysis, in line with existing policies (e.g., by the NIH) that allow sharing of genomic summary results; however, this variable can also be protected at a small additional computational cost if desired. We include an all-ones vector as a covariate in $\mathbf{C}$, which implicitly accounts for mean centering of $\mathbf{X}^{(b)}$. With one round of aggregation, we precompute a small $c$-by-$m_b$ matrix

$$\mathbf{H}^{(b)} = \mathbf{C}^T\mathbf{X}^{(b)} = \sum_{p=1}^{P} \mathbf{C}_p^T\mathbf{X}_p^{(b)}, \tag{25}$$

where each summand is computed locally using plaintext matrices then aggregated in an encrypted form. Next, noting that

$$\tilde{\mathbf{X}}^{(b)} = (\mathbf{X}^{(b)} - \mathbf{C}(\mathbf{C}^T\mathbf{C})^{-1}\mathbf{H}^{(b)})\mathbf{S}^{(b)}, \tag{26}$$

we are able to express $\mathbf{R}_p^{-1}$ (Equation 23) as

$$\mathbf{R}_p^{-1} = \mathbf{S}^{-1}([(\mathbf{X}_p^{(b)})^T\mathbf{X}_p^{(b)} + \rho\mathbf{S}^{-2}] + \mathbf{U}_p\mathbf{E}_p\mathbf{V}_p)^{-1}\mathbf{S}^{-1}, \tag{27}$$

for some matrices $\mathbf{U}_p, \mathbf{V}_p^T \in \mathbb{R}^{m_b \times 2c}$ and $\mathbf{E}_p \in \mathbb{R}^{2c\times 2c}$ (see Supplementary Note 9 for full derivation; note the inner dimension of $2c$). Finally, using the Woodbury identity and letting $\mathbf{Q} := (\mathbf{X}_p^{(b)})^T\mathbf{X}_p^{(b)} + \rho\mathbf{S}^{-2}$ to simplify the notation, we can expand the inverse as

$$(\mathbf{Q}_p + \mathbf{U}_p\mathbf{E}_p\mathbf{V}_p)^{-1} = \mathbf{Q}_p^{-1} - \mathbf{Q}_p^{-1}\mathbf{U}_p(\mathbf{E}_p^{-1} + \mathbf{V}_p\mathbf{Q}_p^{-1}\mathbf{U}_p)^{-1}\mathbf{V}_p\mathbf{Q}_p^{-1}. \tag{28}$$

We have successfully reformulated the computation of $\mathbf{R}_p^{-1}$ as one involving an *inverse of a plaintext matrix* $\mathbf{Q}_p$ and several matrix multiplications with a small inner dimension of $2c$. Note that the new composite inverse matrix

$$\mathbf{L}_p := (\mathbf{E}_p^{-1} + \mathbf{V}_p\mathbf{Q}_p^{-1}\mathbf{U}_p)^{-1} \tag{29}$$

can be efficiently computed using secure MPC protocols, using the eigenfactorization routine introduced in Supplementary Note 5 (MPC-EigenDecomp($\cdot$)).

As a result, we can compute a key matrix $\mathbf{Q}_p^{-1}$ completely locally in plaintext, then use Equations 27 and 28 to compute the multiplication with $\mathbf{R}_p^{-1}$ on the fly using the plaintext $\mathbf{Q}_p^{-1}$. Note that this step is the only expensive matrix multiplication in the ADMM algorithm, and as such our reformulated ADMM offers significant reduction in computational cost. Moreover, we emphasize that, aside from the precomputation of $\mathbf{H}^{(b)}$ and $\mathbf{Q}_p^{-1}$, none of the matrix operations in our ADMM algorithm scales with the number of individuals in the dataset, and thus scales very efficiently to datasets with many samples, as our results show. Our final ADMM algorithm for ridge regression with covariates, leveraging the Woodbury identity technique, is presented in Protocol 12 (changes with respect to Protocol 11 are shown in blue).

---

**Protocol 12** Secure Federated ADMM-Woodbury Algorithm for Ridge Regression with Covariates

---

**Input:** Number of parties $k$, horizontally distributed input matrix $\bar{\mathbf{X}} = (\bar{\mathbf{X}}_1, \ldots, \bar{\mathbf{X}}_k)$ and a covariate matrix $\bar{\mathbf{C}} = (\bar{\mathbf{C}}_1, \ldots, \bar{\mathbf{C}}_k)$, a diagonal matrix $\bar{\mathbf{S}}$ with inverse standard deviations of columns of $\bar{\mathbf{X}}$, target vector $\bar{\mathbf{b}} = (\bar{\mathbf{b}}_1, \ldots, \bar{\mathbf{b}}_k)$, regularization parameter $\lambda$, learning rate $\rho$, number of iterations $\tau$. An overline indicates a plaintext variable.

**Ouput:** A vector $\mathbf{z}$ that satisfies $(\mathbf{A}^T\mathbf{A} + \lambda\mathbf{I})\mathbf{z} \approx \mathbf{b}$, where $\mathbf{A} := (\mathbf{I} - \mathbf{C}(\mathbf{C}^T\mathbf{C})^{-1}\mathbf{C}^T)\mathbf{XS}$

Each party $p$ locally computes $\bar{\mathbf{H}}_p \leftarrow \bar{\mathbf{C}}_p^T\bar{\mathbf{X}}_p$
$\mathbf{H} \leftarrow$ MHE-Aggregate($\bar{\mathbf{H}}_p$)
Each party $p$:
    Initializes $\mathbf{w}_p^0 \leftarrow 0, \mathbf{u}_p^0 \leftarrow 0$ and a global vector $\mathbf{z}^0 \leftarrow 0$
    Locally computes $\bar{\mathbf{Q}}_p^{-1} \leftarrow (\bar{\mathbf{X}}_p^T\bar{\mathbf{X}}_p + \rho\bar{\mathbf{S}}^{-2})^{-1}$
Precompute $\mathbf{E}_p$, $\mathbf{V}_p$, $\mathbf{U}_p$, and $\mathbf{L}_p := (\mathbf{E}_p^{-1} + \mathbf{V}_p\bar{\mathbf{Q}}_p^{-1}\mathbf{U}_p)^{-1}$ in Equation 29 for all parties // MPC-EigenDecomp
**for** $j \in \{0, ..., \tau - 1\}$ **do**
  Each party $p$:
    Locally computes $\mathbf{h} \leftarrow \bar{\mathbf{Q}}_p^{-1}\bar{\mathbf{S}}^{-1}(\bar{\mathbf{b}} + \rho\mathbf{z}^j - \mathbf{u}_p^j)$
    Locally computes $\mathbf{w}_p^{j+1} \leftarrow \bar{\mathbf{S}}^{-1}(\mathbf{h} - \bar{\mathbf{Q}}_p^{-1}\mathbf{U}_p\mathbf{L}_p\mathbf{V}_p\mathbf{h})$
  $\mathbf{w}^{j+1} \leftarrow \frac{1}{k} \cdot$ MHE-Aggregate($\mathbf{w}_p^{j+1}$)
  $\mathbf{u}^j \leftarrow \frac{1}{k} \cdot$ MHE-Aggregate($\mathbf{u}_p^j$)
  Each party $p$ locally computes:
    $\mathbf{z}^{j+1} \leftarrow (\rho\mathbf{w}^{j+1} + \mathbf{u}^j)/(\lambda/k + \rho)$
    $\mathbf{u}_p^{j+1} \leftarrow \mathbf{u}_p^j + \rho(\mathbf{w}_p^{j+1} - \mathbf{z}^{j+1})$
**end for**
**return** $\mathbf{z}^\tau$

---

**Association testing.** Recall from Equation 9 that the association statistic for each variant $\mathbf{x}$ can be computed as

$$\frac{(\tilde{\mathbf{x}}^T(\tilde{\mathbf{y}} - \hat{\mathbf{y}}_{\text{LOCO}}))^2}{\hat{\sigma}_e^2 \cdot (\tilde{\mathbf{x}}^T\tilde{\mathbf{x}})}.$$

Let $\sigma_x$ be the standard deviation of $\mathbf{x}$, and

$$\mathbf{P} = \mathbf{I}^{(n\times n)} - \mathbf{C}(\mathbf{C}^T\mathbf{C})^{-1}\mathbf{C}^T.$$

We can express

$$\tilde{\mathbf{x}} = \sigma_x^{-1}\mathbf{P}\mathbf{x}.$$

Since we include a column of ones in the covariate matrix $\mathbf{C}$ for mean correction, we do not need to apply a separate mean centering to $\mathbf{x}$. Also, since the scaling factor $\sigma_x$ cancels between the denominator and the numerator, we can set it to one without affecting the result, i.e., let $\tilde{\mathbf{x}} = \mathbf{P}\mathbf{x}$.

Thus, it suffices to compute

$$s := \frac{\mathbf{x}^T \mathbf{P} \hat{\mathbf{y}}^*_{\text{resid,LOCO}}}{\hat{\sigma}_e \cdot \sqrt{\mathbf{x}^T \mathbf{P} \mathbf{x}}},$$

where $\hat{\mathbf{y}}^*_{\text{resid,LOCO}} = \tilde{\mathbf{y}} - \hat{\mathbf{y}}_{\text{LOCO}}$. We first directly calculate $\hat{\mathbf{y}}^*_{\text{resid,LOCO}}$ and $\hat{\sigma}_e = \|\tilde{\mathbf{y}} - \hat{\mathbf{y}}_{\text{LOCO}}\|_2 / \sqrt{n-c}$. Then, we obtain the numerator by first computing

$$\mathbf{w} := \mathbf{P} \hat{\mathbf{y}}^*_{\text{resid,LOCO}}$$

then computing $\mathbf{x}^T \mathbf{w}$. Note that $\mathbf{w}$ is computed once per chromosome and shared across all variants.

Next we consider $\mathbf{x}^T \mathbf{P} \mathbf{x}$ in the denominator. Exploiting the fact that $\mathbf{C}^T \mathbf{C}$ is a small constant-size matrix, we first obtain $\mathbf{R}$ such that $\mathbf{R}\mathbf{R}^T = (\mathbf{C}^T \mathbf{C})^{-1}$ using our MPC eigendecomposition routine. We can now express

$$\mathbf{P} = \mathbf{I} - \mathbf{C}\mathbf{R}\mathbf{R}^T \mathbf{C}^T.$$

With

$$\mathbf{u} := \mathbf{R}^T \mathbf{C}^T \mathbf{x}, \tag{30}$$

we can compute

$$\mathbf{x}^T \mathbf{P} \mathbf{x} = \mathbf{x}^T \mathbf{x} - \mathbf{u}^T \mathbf{u}. \tag{31}$$

We further consolidate the computation above as follows. After computing $\mathbf{w}$, we take a single pass over the input genotype matrix to compute the following terms for each genotype $\mathbf{x}$:

$$\mathbf{x}^T \mathbf{x}, \ \mathbf{C}^T \mathbf{x}, \ \mathbf{w}^T \mathbf{x}.$$

Note that the first two are computed in plaintext and easily converted to secret shares, where each party sets their own share to the computed plaintext value such that the implicit sum of shares simply becomes the overall sum (i.e., using MPC-Aggregate). The third term involves a ciphervector $\mathbf{w}$ and thus can be treated as ciphervector-plainmatrix multiplication, which is also utilized in the PCA-based workflow. Given these terms, we can compute $\mathbf{u}$, then $\mathbf{x}^T \mathbf{P} \mathbf{x}$ using Equations 30 and 31 via MPC. Finally, we multiply the numerator with the inverse of the denominator (taking $\hat{\sigma}_e$ into account) to obtain the association statistics.

**Overall workflow for LMM-based GWAS.** Here we summarize the workflow of the LMM-based SF-GWAS workflow. We first apply the same quality control procedure as in the PCA-based workflow to obtain a filtered dataset for each party. Next, joint PCA is optionally performed to obtain fixed-effect covariates based on top principal components. Corresponding to Level 0 of REGENIE, we divide the genome into blocks, then for each block run our secure and federated ADMM-Woodbury algorithm (Protocol 12) to obtain local phenotype predictors for different values of the regularization parameter. For Level 1 of REGENIE, we aggregate the local predictors into a combined feature matrix and use our CGD algorithm (Protocol 10) to obtain the LOCO genome-wide phenotype predictions for each chromosome. We repeat this estimation procedure for cross validation and measure prediction accuracy on a hold-out set to select the best variance parameter for the genetic effect. With the final set of genome-wide LOCO predictors, we follow the steps in the previous section to compute association statistics for all variants.

**Notes on computational complexity.** SF-GWAS greatly reduces the runtime of a direct implementation of REGENIE in a federated setting. By using our improved ADMM-Woodbury algorithm, we delegate large matrix inverse operations to be performed locally in plaintext. Since in practice the overhead of cryptographic operations greatly overshadows that of plaintext computation, in our complexity analysis we only consider homomorphic operations over the encrypted data.

The implementation of ADMM-Woodbury is separated into two components, the precomputation of a small matrix inverse in the Woodbury identity and the main ADMM iterations. For the precomputation of the $\mathbf{L}_p$ matrix (Equation 29), each party performs a $m_b$-by-$m_b$ matrix multiplication around $2c$ times where

$c$ is the number of covariates, where $m_b$ is the blocksize. Each main iteration is dominated by the work of multiplying a $m_b$-by-$m_b$ plaintext matrix $\mathbf{Q}_p^{-1}$ twice with a ciphervector, combined with cipher-cipher multiplications with precomputed matrices $\mathbf{U}_p$, $\mathbf{L}_p$, and $\mathbf{V}_p$. Like in the plaintext setting, matrix vector multiplication in HE scales linearly with the size of the matrix. Since ridge regression in Level 0 is computed for each block ($B$), cross-validation fold ($K$), and regularization parameter ($R$), the complexity of Level 0 is $O(KBRm_b^2\tau_a)$ where $\tau_a$ is the number of iterations in ADMM. Since $m = Bm_b$ by definition, this can be expressed as $O(m_bKRM\tau_a)$. We note that this is a much better asymptotic runtime than naïvely using CGD for Level 0. Each iteration of CGD scales linearly with the size of the matrix being multiplied, which is $n$-by-$m_b$ in our case. Therefore, the total runtime of Level 0 with CGD is $O(m_bKBRn\tau_c)$, or equivalently $O(KRmn\tau_c)$, which is a factor of $n/m_b$ larger than our ADMM-Woodbury solution. For a dataset of $n = 10^5$ and $m_b = 10^3$, this amounts to a factor of 100 improvement using ADMM-Woodbury.

For Level 1, since we evaluate the CGD subroutine lazily without explicitly constructing the design matrix, we distribute the work in a way where each party multiplies their respective plaintext genotype matrix with a vector. Since the size of the matrix is $n$-by-$BR$, and there are $KR$ different ridge regressions that must be performed, the runtime complexity of Level 1 is $O(KR^2nB\tau_c)$, where $\tau_c$ is the number of iterations in CGD (typically 30), and $K$ and $R$ are small numbers (default values of 5 in REGENIE).

Lastly, to compute association statistics, much of the work can be done in plaintext. The chromosome-specific LOCO residual vectors (observed phenotypes minus the LOCO predictors of genetic effect) must be covariate corrected and then multiplied by the genotype matrix for the corresponding chromosome. This is equivalent to one matrix-vector multiplication between the full genotype matrix (plaintext) and a residual vector (ciphertext). While this leads to an asymptotic complexity of $O(nm)$, in practice since only one such multiplication is needed and because cipher-plain multiplications are considerably more efficient than cipher-cipher multiplications, computing association statistics is the quickest of the three levels of computation.

**Numerical stability.** There are several hyperparameters and statistics to take note of with SF-GWAS's use of iterative methods for ridge regression that factor into the numerical stability and convergence time of SF-GWAS. Our ADMM-Woodbury algorithm for Level 0 depends on the hyperparameter $\rho$, which represents the step size used in the ADMM iterations (i.e., learning rate). We have found that in practice, an effective $\rho$ can be determined as a function of the number of individuals, since the genomic block size is typically chosen within a fixed range of 1K to 5K variants for REGENIE. In our experiments, we set $\rho$ to the number of individuals in the local dataset divided by the number of cross-validation folds.

Another hyperparameter of interest for both CGD and ADMM-Woodbury is the number of iterations. The convergence of both methods are generally dependent on the condition number of the input matrix [23, 27] and thus rely heavily on the magnitude of the regularization parameter relative to the input matrix. Therefore, we implemented an adaptive approach that checks for convergence at regular intervals and terminates when a suitable solution has been obtained. We also employed a warm start strategy whereby solutions from a higher value of the regularization parameter is used to initialize the run with a lower value, which considerably improved convergence overall.

We also note that precision is challenging to maintain in general in secure computation protocols given their dependence on fixed-point representation of continuous numbers and the possibilities of numerical under/overflow. Throughout our algorithm, we carefully managed the range of data values by scaling intermediate results accordingly (e.g., dividing by $\sqrt{n}$ when computing the sum of $n$ values for large $n$). As our results show, the combination of these strategies allow SF-GWAS to obtain accurate analysis results.

**Parallel processing of multiple phenotypes.** A key feature of REGENIE [17] aimed at computational efficiency is the support for concurrent analysis of multiple phenotypes. Although our SF-GWAS implementation currently analyzes one phenotype at a time, the modular design of our software allows for modifications to efficiently analyze multiple phenotypes. The speedup typically associated with large matrix operations (exploited by REGENIE) is not directly applicable in our setting due to the properties of encrypted operations. Nevertheless, SF-GWAS uses optimized encrypted matrix operations, which helps reduce the computational cost of analyzing multiple phenotypes. Moreover, across all our protocols, any step that is unrelated to the phenotype (e.g., PCA on genotypes) can be kept as is and executed only once—only the computations involving the phenotype need to be updated to process multiple phenotypes in parallel (e.g., the steps that depend on $\bar{\mathbf{y}}_p$ in Protocols 7 and 9).

### Supplementary Note 7: Security of SF-GWAS

The security of SF-GWAS is based on the honest-but-curious (semi-honest) adversarial model, where the collaborating entities follow the protocol faithfully as given, but may try to infer private information based on all of the information provided to them during the protocol. This is in contrast to the malicious security setting, where adversaries can freely deviate from the given protocols. We further discuss the choice of our security model and possible system extensions later in this section.

Another key assumption in our security model is that at least one party does not collude (i.e., share data) with others. If all parties collude, then the secret shares in either the MHE or MPC schemes can be combined to reveal the secret. The use of an auxiliary party (trusted dealer; $\mathsf{CP}_0$) in our MPC framework introduces an additional requirement that this party does not collude with any of the main parties. However, since the use of MPC in SF-GWAS is limited to aggregate-level intermediate analysis results, a violation of this latter requirement does not immediately lead to a leakage of genotype or phenotype data. There is still a residual risk that some information about the private input data (e.g., membership in a study cohort [28]) could be inferred from the leaked aggregate-level data in the event of a collusion. This arguably poses a significantly smaller risk than the leakage of raw input data, as reflected in the NIH Genomic Data Sharing Policy, which in principle allows the sharing of aggregate genomic statistics. We emphasize that, without collusion, our system keeps even the intermediate results confidential.

Given the above security assumptions, SF-GWAS formally ensures the confidentiality of each private input dataset from entities other than the respective data holder, except for what can be inferred from the necessary outputs of the analysis, including the quality control filter and association statistics. The LMM-based workflow additionally outputs the genomic heritability parameter chosen by cross-validation and the convergence status of iterative algorithms (i.e., CGD and ADMM) at fixed intervals. Optionally, global SNP variances can be shared among parties to further speed up the LMM-based workflow, as incorporated in our experiments.

The security strength of cryptosystems is usually represented in terms of the number of bits. For example, 128-bit (computational) security means that $2^{128}$ operations are required asymptotically to break the security of an encryption scheme. SF-GWAS is based on MHE security parameters, as specified in Supplementary Note 2, that provide 128-bit (computational) security. Several routines in MHE and MPC (e.g., MHE-MPC conversion and the truncation protocol in MPC) additionally rely on the notion of statistical security, which means that a potential leakage of private information is information-theoretically bounded in terms of the statistical distance from the uniform random distribution. For these routines, the SF-GWAS security parameters achieve 128-bit statistical security for both MHE and MPC, accounting for at least $2^{30}$ such operations via a union bound.

The main application setting of our methods involves academic researchers and entities who share scientific interests in running a joint analysis, but are prevented from doing so due to data sharing policies and regulations. Thus, these parties share an incentive to follow the protocols correctly to obtain accurate results, and their primary concern is what could be revealed to each other during the process. Moreover, we envision our protocols being provided as automated workflows on data analysis platforms, where users have limited means to modify the program during execution. The role of a trusted dealer could also be provided by such platforms as an automated service, for example, by creating an ephemeral virtual machine on-the-fly to generate and share the necessary correlated randomness. Therefore, we believe that our honest-but-curious security model without collusion provides a formal guarantee that is also practically meaningful. Our choice of the security model also ensures that the protocols can be applied efficiently to large-scale datasets, as demonstrated by our results.

Nevertheless, we recognize the value of stronger security protections even against malicious adversaries. To this end, one could introduce mechanisms for parties to verify each other's inputs and computations, e.g., based on zero-knowledge proofs in HE [29, 30] and authenticated secret shares in MPC [31]. Incorporating these extensions into SF-GWAS while preserving its practical performance is a meaningful future direction.

### Supplementary Note 8: Runtime Estimation and Complexity Analysis

In this section, we provide practical guidelines for estimating the required runtime of SF-GWAS for a given dataset. Our estimates are based on the e2-highmem-16 virtual machines (VMs) in the Google Cloud Platform (GCP [32]) running in the same zone. Note that the initial setup (e.g., key generation) usually takes less than a minute and is independent of the dataset size. The primary computational bottlenecks of SF-GWAS are the matrix multiplications involving the large genotype matrix; because these steps are parallelized, providing SF-GWAS with more vCPUs on larger VM types will lead to faster runtimes.

For interested readers, we also provide a theoretical complexity analysis of our protocols in Supplementary Tables 3-5, where we report the costs of the main analysis routines concretely in terms of the number of invocations of building-block cryptographic operations for both MHE and MPC. The runtime and communication costs of the individual operations based on a range of settings, representing different network communication speeds and the number of parties, are summarized in Supplementary Table 2.

**Quality control.** The bottleneck in this phase is the invocation of MPC protocols for division and comparison on vectors of length equal to the total number of SNPs in the dataset. Given enough network bandwidth, one can efficiently parallelize this routine by assigning a separate thread with a separate set of network channels to work on a portion of the vector at a time; this feature is implemented in SF-GWAS. Using 12 threads on each VM, the estimated runtime for this phase is around 2-4 minutes for every 500K SNPs. We observed similar performance for both two-party settings (S-GWAS datasets) and six/seven-party settings (eMERGE and UKB). Note that for 93 million imputed SNPs in UKB, this step took 4.5 hours.

**Principal component analysis.** In addition to the dataset dimensions, the runtime of PCA depends on several factors, including the number of power iterations (20 by default, recommended in most settings); the number of principal components (PCs) to extract (typically 5 or 10); oversampling parameter (typically 10; represents how many components to additionally extract and later discard to enhance approximation accuracy). Note that given our distance-based SNP pruning strategy for PCA, we used about 23-27K SNPs for PCA in all our datasets. Based on our experiments with the default parameters, we observed a linear scaling with approximately 1.5 to 3 hours of runtime per 10K individuals. Increasing the size of the SNP set is expected to linearly scale the runtime by the same factor (Supplementary Fig. 5). For the largest UKB data with six parties and a total of 276K individuals, the runtime of PCA for extracting 10 PCs was 44 hours.

**Association tests.** The runtime of this phase is largely determined by the time it takes to multiply either the orthogonal basis of the covariates (including PCs) for linear regression-based association tests, or a matrix derived from it with the same dimensions for the logistic regression-based version, with the local plaintext genotype matrix. This can be parallelized across genomic blocks, and each matrix multiplication can also be sped up by assigning more threads to it. Note that parties with different numbers of data samples will have different runtimes for this step, so the overall runtime depends on the slowest party to finish performing the matrix multiplications. Depending on the size of the local dataset, each party can leverage a different VM type to reduce their local runtime. For large datasets like UKB, we recommend running SF-GWAS in two stages: the first stage using e2-highmem-16 VMs in GCP (16 vCPUs and 128 GB RAM) or similar to perform everything up to (and including) the PCA step, then the second stage for association tests using more powerful VMs as needed. Our UKB analysis utilized n2-highmem-64 (64 vCPUs with 512 GB of RAM) and n2-highmem-128 (128 vCPUs with 864 GB of RAM) VMs depending on the local dataset size. We recommend allocating 8 GB RAM per vCPU and assigning 8 threads per block being processed; e.g., using the n2-highmem-64 VM, we processed 8 blocks in parallel—a parameter that can be configured in SF-GWAS. A ballpark estimation of runtime can be obtained by taking 0.5 to 1 hour of runtime per 500K SNPs (passing QC), 10K individuals, and 15 covariates (assuming linear scaling in each of these dimensions), then dividing by the number of blocks that are processed in parallel. Note that, for UKB data with 13,515,893 QC-passed SNPs, 276K individuals in total, and 36 covariates (including PCs), the overall runtime of this step was 77.8 hours leveraging a combination of aforementioned high-performance VMs.

**LMM-based workflow.** The runtime of the LMM-based workflow on the smaller lung cancer dataset was 2.8 days (68.3 hours). Most of the runtime could be attributed to Level 0 (61.4 hours, compared to 6.33 hours of Level 1), which trains a large number of local ridge regression models across genomic blocks. On the UKB dataset, the total runtime was 6 days (145 hours). Here, Level 0 (69.3 hours) and Level 1 (73.7 hours) had comparable runtimes due to the much larger number of individuals in the dataset. UKB data (array genotypes) includes 1.5x the number of variants and around 44.5x the number of individuals compared to the lung cancer data. As described in Supplementary Note 6, our ADMM-Woodbury algorithm for Level 0 precomputes the inverse of a $m_b$-by-$m_b$ matrix in plaintext for each block (where $m_b$ is the block size) and performs all subsequent operations using this matrix, effectively removing dependence of runtime on the number of individuals $n$, except for the cost of precomputation. This leads to the comparable Level 0 runtimes between the two datasets despite the much greater size of UKB. For Level 1, our runtime scales approximately linearly in both $n$ and the number of genomic blocks. The 11.6x increase in Level 1 runtime for UKB accounts for the greater $n$ (44.5x) offset by the greater number of cores (4x) used for the UKB experiment. All other steps, including association testing, have considerably smaller runtime compared to these two steps (Levels 0 and 1). Using similar computing resources as our UKB experiment (e.g., n2-highmem-64 for each party), the overall runtime of the LMM workflow can therefore be approximated as the sum of approximately one day for every 200K variants for Level 0, and around 6 hours for every 100K individuals and 200K variants for Level 1. Note that once Level 1 is performed, the resulting genome-wide phenotype predictions can be used to test the association of a larger set of variants if desired. The runtime for the association testing portion can be estimated in a similar way as in the PCA-based workflow.

### Supplementary Note 9: Derivation of Our ADMM-Woodbury Algorithm

The ADMM-Woodbury algorithm for ridge regression (Protocol 9) requires the computation of the inverse of the large scale encrypted matrix $\mathbf{R}$:

$$\mathbf{R}_p^{-1} = ((\tilde{\mathbf{X}}_p)^T \tilde{\mathbf{X}}_p + \rho \mathbf{I}^{(m_b \times m_b)})^{-1}. \quad (32)$$

For simplicity, we drop the superscript $b$ in the following. We note that here $\tilde{\mathbf{X}}_p$ is not available in plaintext, as it is meant to be standardized and covariate-corrected based on the global matrix $\tilde{\mathbf{X}}_p$. We instead write $\tilde{\mathbf{X}}_p$ as a combination of the local cleartext (non-standardized) input $\mathbf{X}$, covariate matrix $\mathbf{C}$ (augmented with an all-ones vector for mean centering), and a diagonal matrix $\mathbf{S}$ including inverse standard deviations for each column of $\mathbf{X}$. Using the Woodbury matrix identity [26], we can then reformulate $\mathbf{R}_p^{-1}$ in terms of the inverse of a plaintext matrix $\mathbf{Q}_p$ and several matrix multiplications with small inner dimensions:

Note that $\tilde{\mathbf{X}}$ can be written as

$$\tilde{\mathbf{X}} = (\mathbf{X} - \mathbf{C}(\mathbf{C}^T\mathbf{C})^{-1}\mathbf{H})\mathbf{S}, \quad (33)$$

where the small $c$-by-$m_b$ matrix $\mathbf{H} = \mathbf{C}^T\mathbf{X} = \sum_{p=1}^{P} \mathbf{C}_p^T \mathbf{X}_p$ is obtained with one round of aggregation. Therefore, by combining Equations 32 and 33, we have

$$\begin{aligned}
\mathbf{R}_p^{-1} &= (((\mathbf{X} - \mathbf{C}(\mathbf{C}^T\mathbf{C})^{-1}\mathbf{H})\mathbf{S})^T(\mathbf{X} - \mathbf{C}(\mathbf{C}^T\mathbf{C})^{-1}\mathbf{H})\mathbf{S} + \rho\mathbf{I}^{(m_b \times m_b)})^{-1} \\
\mathbf{S}\mathbf{R}_p^{-1}\mathbf{S} &= ((\mathbf{X}^T - \mathbf{H}^T(\mathbf{C}^T\mathbf{C})^{-1}\mathbf{C}^T)(\mathbf{X} - \mathbf{C}(\mathbf{C}^T\mathbf{C})^{-1}\mathbf{H}) + \rho\mathbf{S}^{-2})^{-1} \\
&= (\mathbf{X}^T\mathbf{X} + \rho\mathbf{S}^{-2} - \mathbf{H}^T(\mathbf{C}^T\mathbf{C})^{-1}\mathbf{C}^T\mathbf{X} - \mathbf{X}^T\mathbf{C}(\mathbf{C}^T\mathbf{C})^{-1}\mathbf{H} + \mathbf{H}^T(\mathbf{C}^T\mathbf{C})^{-1}\mathbf{H})^{-1} \\
&= (\mathbf{X}^T\mathbf{X} + \rho\mathbf{S}^{-2} - \mathbf{X}^T\mathbf{C}(\mathbf{C}^T\mathbf{C})^{-1}\mathbf{H} - \mathbf{H}^T(\mathbf{C}^T\mathbf{C})^{-1}(\mathbf{C}^T\mathbf{X} - \mathbf{H}))^{-1}.
\end{aligned}$$

By denoting

$$\mathbf{U} = \begin{bmatrix}\mathbf{H}^T & \mathbf{X}^T\mathbf{C}\end{bmatrix}^{(m_b \times 2c)}, \qquad \mathbf{E} = \begin{bmatrix}-\mathbf{C}^T\mathbf{C}^{(c \times c)} & 0 \\ 0 & -\mathbf{C}^T\mathbf{C}^{(c \times c)}\end{bmatrix}^{-1}, \text{ and } \qquad \mathbf{V} = \begin{bmatrix}\mathbf{C}^T\mathbf{X} - \mathbf{H} \\ \mathbf{H}\end{bmatrix}^{(2c \times m_b)},$$

we can rewrite $\mathbf{R}_p^{-1}$ as

$$\mathbf{R}_p^{-1} = \mathbf{S}^{-1}([\mathbf{X}_p^T\mathbf{X}_p + \rho\mathbf{S}^{-2}] + \mathbf{UEV})^{-1}\mathbf{S}^{-1}. \tag{34}$$

Applying the Woodbury identity with $\mathbf{Q} = \mathbf{X}_p^T\mathbf{X}_p + \rho\mathbf{S}^{-2}$, we obtain

$$\mathbf{R}_p^{-1} = \mathbf{S}^{-1}(\mathbf{Q} + \mathbf{UEV})^{-1}\mathbf{S}^{-1} = \mathbf{S}^{-1}[\mathbf{Q}^{-1} - \mathbf{Q}^{-1}\mathbf{U}(\mathbf{E}^{-1} + \mathbf{VQ}^{-1}\mathbf{U})^{-1}\mathbf{VQ}^{-1}]\mathbf{S}^{-1}, \tag{35}$$

where $(\mathbf{E}^{-1} + \mathbf{VQ}^{-1}\mathbf{U})^{-1}$ is the inverse of a very small constant-size matrix of size $2c \times 2c$.

### References


[1] Hyunghoon Cho, David J Wu, and Bonnie Berger. Secure genome-wide association analysis using multiparty computation. *Nature biotechnology*, 2018.

[2] Marcella Hastings, Brett Hemenway, Daniel Noble, and Steve Zdancewic. Sok: General purpose compilers for secure multi-party computation. In *2019 IEEE symposium on security and privacy (SP)*, pages 1220–1237. IEEE, 2019.

[3] Eleftheria Makri, Dragos Rotaru, Frederik Vercauteren, and Sameer Wagh. Rabbit: Efficient comparison for secure multi-party computation. In *International Conference on Financial Cryptography and Data Security*, pages 249–270. Springer, 2021.

[4] Christian Mouchet, Juan R. Troncoso-pastoriza, Jean-Philippe Bossuat, and Jean Pierre Hubaux. Multiparty homomorphic encryption from ring-learning-with-errors. In *Proceedings on Privacy Enhancing Technologies Symposium*, 2021.

[5] Jung Hee Cheon, Andrey Kim, Miran Kim, and Yongsoo Song. Homomorphic encryption for arithmetic of approximate numbers. In *Proceedings of the International Conference on the Theory and Application of Cryptology and Information Security (ASIACRYPT)*, 2017.

[6] Lattigo: A Library For Lattice-Based Homomorphic Encryption in Go. `https://github.com/tuneinsight/lattigo`, (accessed: July 2022).

[7] Yvo G Desmedt. Threshold cryptography. *European Transactions on Telecommunications*, 5(4):449–458, 1994.

[8] Andrey Kim, Antonis Papadimitriou, and Yuriy Polyakov. Approximate homomorphic encryption with reduced approximation error. Cryptology ePrint Archive, Paper 2020/1118, 2020. URL `https://eprint.iacr.org/2020/1118`.

[9] Kyoohyung Han and Dohyeong Ki. Better Bootstrapping for Approximate Homomorphic Encryption. In *CT-RSA*, 2020.

[10] David Froelicher, Juan R Troncoso-Pastoriza, Jean Louis Raisaro, Michel A Cuendet, Joao Sa Sousa, Hyunghoon Cho, Bonnie Berger, Jacques Fellay, and Jean-Pierre Hubaux. Truly privacy-preserving federated analytics for precision medicine with multiparty homomorphic encryption. *Nature communications*, 12(1):1–10, 2021.

[11] Xiaoqian Jiang, Miran Kim, Kristin Lauter, and Yongsoo Song. Secure Outsourced Matrix Computation and Application to Neural Networks. In *Proceedings of the 2018 ACM SIGSAC CCS*, pages 1209–1222, 2018.

[12] Marcelo Blatt, Alexander Gusev, Yuriy Polyakov, and Shafi Goldwasser. Secure large-scale genome-wide association studies using homomorphic encryption. *Proceedings of the National Academy of Sciences*, 117(21):11608–11613, 2020.

[13] Morten Dahl, Chao Ning, and Tomas Toft. On Secure Two-Party Integer Division. In *Inter. Conf. on Financial Cryptography and Data Security*, 2012.

[14] Nathan Halko, Per-Gunnar Martinsson, and Joel A Tropp. Finding structure with randomness: Probabilistic algorithms for constructing approximate matrix decompositions. *SIAM review*, 53(2):217–288, 2011.

[15] David Froelicher, Juan R Troncoso-Pastoriza, Apostolos Pyrgelis, Sinem Sav, Joao Sa Sousa, Jean-Philippe Bossuat, and Jean-Pierre Hubaux. Scalable privacy-preserving distributed learning. In *Proceedings on Privacy Enhancing Technologies Symposium*, volume 2, pages 323–347, 2021.

[16] David Froelicher, Juan R. Troncoso-Pastoriza, Apostolos Pyrgelis, Sinem Sav, Joao Sa Sousa, Jean-Philippe Bossuat, and Jean-Pierre Hubaux. Scalable Privacy-Preserving Distributed Learning. *Proceedings on Privacy Enhancing Technologies Symposium*, 2021.

[23] *Iterative Solution of Large Sparse Systems of Equations*. Springer, Cham, Switzerland, 2016.

[24] Stephen Boyd, Neal Parikh, and Eric Chu. *Distributed optimization and statistical learning via the alternating direction method of multipliers*. Now Publishers Inc, 2011.

[25] Wenting Zheng, Raluca Ada Popa, Joseph E Gonzalez, and Ion Stoica. Helen: Maliciously secure coopetitive learning for linear models. In *Proceedings of IEEE Symposium on Security and Privacy (SP)*, pages 724–738, 2019.

[26] William W Hager. Updating the inverse of a matrix. *SIAM review*, 31(2):221–239, 1989.

[27] Robert Nishihara, Laurent Lessard, Benjamin Recht, Andrew Packard, and Michael I. Jordan. A General Analysis of the Convergence of ADMM. *International Conference on Machine Learning*, 32, 02 2015.

[28] Nils Homer, Szabolcs Szelinger, Margot Redman, David Duggan, Waibhav Tembe, Jill Muehling, John V Pearson, Dietrich A Stephan, Stanley F Nelson, and David W Craig. Resolving individuals contributing trace amounts of dna to highly complex mixtures using high-density snp genotyping microarrays. *PLoS genetics*, 4(8):e1000167, 2008.

[29] Vadim Lyubashevsky, Ngoc Khanh Nguyen, and Gregor Seiler. Practical lattice-based zero-knowledge proofs for integer relations. *ACM CCS*, 2020.

[30] Rupeng Yang, Man Ho Au, Zhenfei Zhang, Qiuliang Xu, Zuoxia Yu, and William Whyte. Efficient lattice-based zero-knowledge arguments with standard soundness: construction and applications. *CRYPTO*, 2019.

[31] Ivan Damgård, Valerio Pastro, Nigel Smart, and Sarah Zakarias. Multiparty computation from somewhat homomorphic encryption. In *Annual Cryptology Conference*, pages 643–662. Springer, 2012.

[32] Google Cloud Computing Services. `https://cloud.google.com/`, (accessed: August 2022).
